# Apical-to-basal graded ROS metabolism in intact *Hydra* leads to distinct levels of injury-induced ROS signaling in apical and basal regenerating tips

**DOI:** 10.1101/2022.10.04.510867

**Authors:** Nenad Suknovic, Silke Reiter-Karam, Osvaldo Chara, Wanda Buzgariu, Denis Martinvalet, Brigitte Galliot

## Abstract

After mid-gastric bisection, *Hydra* regenerates a head from the lower half and a basal disc from the upper one. What signals elicit two distinct regenerative responses in bisected *Hydra* remains unknown. A mathematical modeling approach based on quantitative data linked to MAPK activation and injury-induced cell death predicts an immediate release of a locally restricted short-lived signal in apical-regenerating tips. We found that Reactive oxygen species (ROS) fulfill this role as evidenced by the injury-induced production of hydrogen peroxide (H_2_O_2_), three-fold higher in apical-regenerating tips than in basal ones. By contrast, mitochondrial superoxide (mtO_2_^.-^) is similarly produced on each side of the cut, playing a positive role on wound healing as mtO_2_^.-^ scavenging delays healing whereas knocking-down *Super Oxide Dismutase* (*SOD*) leads to mtO_2_^.-^ accumulation and acceleration of wound-healing. In intact *Hydra*, the ROS-processing enzyme activities are inversely graded along the body column, basal-to-apical for SOD and apical-to-basal for catalase, explaining the asymmetrical levels of H_2_O_2_ after bisection. High H_2_O_2_ levels trigger injury-induced cell death via paracrine signaling in apical-regenerating tips, where NOX4 and CYBB enzymes amplify them. Hence, the asymmetrical regulation of H_2_O_2_ levels immediately after amputation is crucial to activate two distinct regenerative responses in *Hydra*.

## INTRODUCTION

Organisms that elicit whole-body regeneration or appendage regeneration respond to amputation by achieving a complex cellular remodeling that relies on the combination of cell death, cell dedifferentiation, cell proliferation and cell differentiation^1–3^. One major question is how the injured organism discriminates between a simple wound healing response and a wound healing response combined to the activation of a patterning genetic program that leads to the 3D reconstruction of the missing structures. The freshwater *Hydra* polyp offers an interesting paradigm as bisection along its body column drives wound healing, a symmetrical process on each side of the cut, as well as distinct molecular and cellular responses in the regenerating tips, leading to regeneration of the apical part (head) from the lower part and regeneration of the basal region (foot) from the upper part.

*Hydra* is made of two cell-layers, epidermis and gastrodermis, and its cells derive from three stem cell populations that cannot replace each other, the epidermal and gastrodermal epithelial stem cells (eESCs, gESCs), and the multipotent interstitial stem cells (ISCs)^4–6^. Upon mid-gastric bisection, apical-regenerating (AR) tips exhibit an immediate wave of cell death that affects cells from the interstitial lineage^7^. The dying cells transiently release signals such as Wnt3 that trigger the mitotic division of the surrounding cells paused in G2 and the up-regulation of *Wnt3* expression in the gESCs (**Fig.1A**). This cascade of events that rapidly launches the apical regeneration program after mid-gastric bisection, is based on the asymmetric activation of the MAPK/RSK/CREB pathway following amputation, which triggers cell death in AR tips^8–10^.

**Fig. 1.**
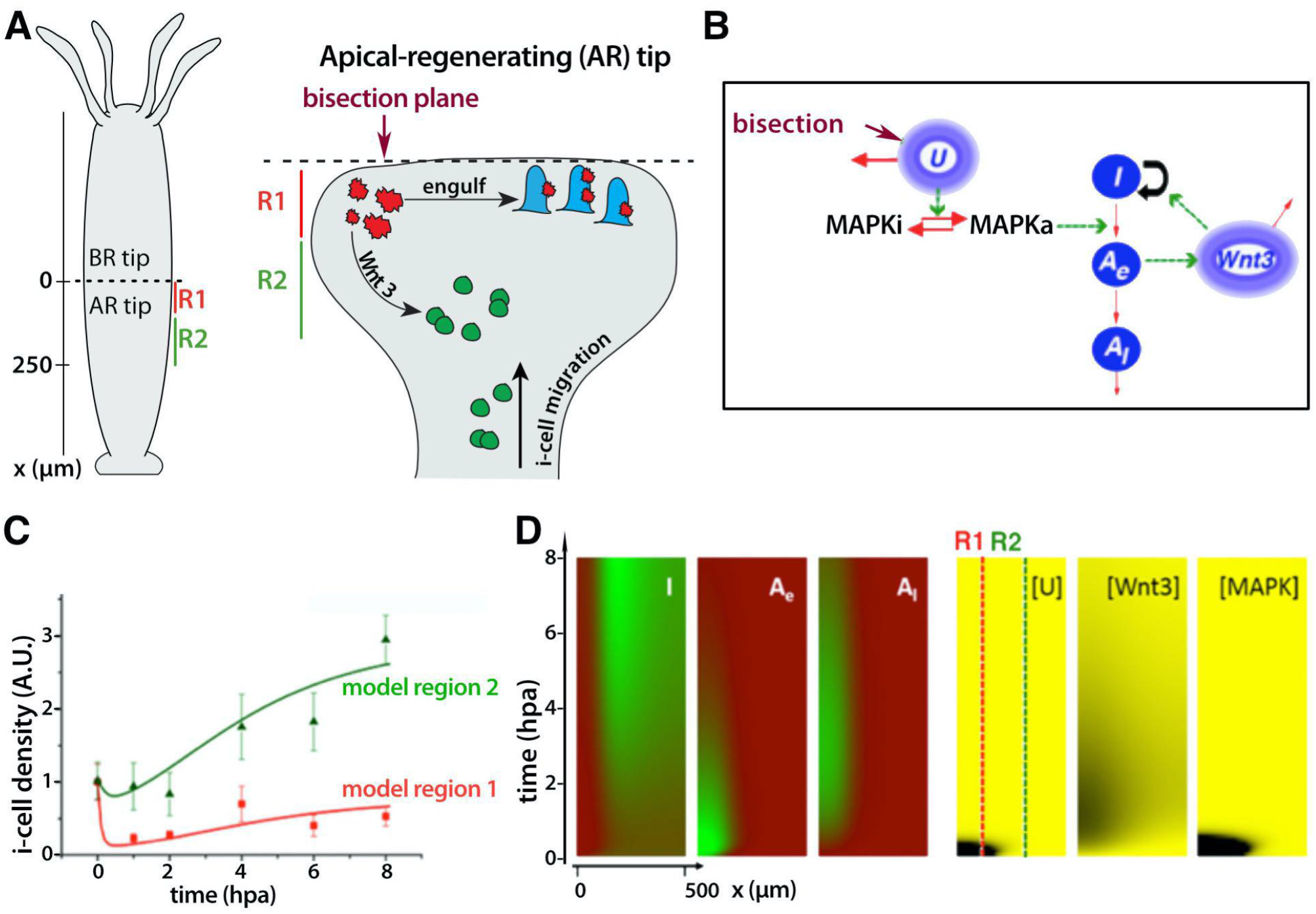
Mathematical Modeling of the immediate injury-induced signaling that triggers cellular remodeling in head regenerating tips. **(A)** Left: Schematical representation of *Hydra* where the amputation plane is located at *x* = 0. R1: region-1 (0-100 µm to the amputation plane), R2: region 2 (100-200 µm underneath). Right: Cellular remodeling occurring in apical-regenerating (AR) tips during the first hours after mid-gastric bisection as reported in ref.7, ref.10. Cell death takes place under the bisection plane within minutes. The dying cells (red) transiently release Wnt3 that activates β-catenin signaling in the surrounding cells, leading interstitial progenitors (green) to synchronously undergo cell division and gastrodermal Epithelial Stem Cells (gESCs) to express Wnt3. In parallel, injury signals attract interstitial progenitors towards the wound. **(B)** Schematic view of the mathematical model depicting the cellular network, the intracellular and extracellular signaling in AR tips. “*U*” corresponds to yet unidentified signal released at the amputation plane, which, by activation of the MAPK pathway (MAPKi transformed into MAPKa), induces death of interstitial cells (*I*) with early (*A*_*e*_) and late (*A*_*l*_) apoptotic cells. The Wnt3 signal (*W*) released by the early apoptotic cells triggers the proliferation of surrounding interstitial cells (*I*). **(C)** The model is fitted to previous experimental data: Dots and error bars represent experimental averages and standard deviations (SDs) of interstitial cells densities calculated from ref.7, continuous curves depict the best-fit model simulation. Red squares and green triangles correspond to values recorded in region-1 and region 2 respectively. For details, see Methods section and **Supplementary Figure S1. (D)** Space-time patterns reflecting the model dynamics. The intensity of green (black) color in each rectangular panel represents the cell density (signal concentration) where the background is red (yellow). Space and time go from left (amputation plane) to right and from bottom (time of head amputation) to top, respectively. This model, based on quantitative data of cell death and cell densities obtained in standard and pharmacologically altered conditions, predicts the activity of a putative injury-induced signal [*U*] restricted in time and space. hpa: hours post-amputation.

The aim of this study was double, to characterize the signals released at the time of injury to identify those that contribute to wound healing on one hand and those that drive the immediate asymmetric activation of MAPK signaling after bisection on the other hand. To do so, we first applied a mathematical modelling approach on quantitative data generated in previous studies on injury-induced cell death and MAPK activation^7,10^ and we deduced the temporal and spatial properties of a predicted injury-induced signal specific to apical regeneration. Next, we tested whether Reactive Oxygen Species (ROS) would fulfill the expected characteristics of this mathematically predicted injury-induced signal. ROS appear as suitable candidate molecules as both superoxide (O_2_^.-^) and hydrogen peroxide (H_2_O_2_) are immediately produced upon injury^11^ in the absence of any transcriptional response, in both vertebrate and invertebrate wound healing contexts such as in zebrafish larvae^12^, *Drosophila* embryo^13,14^ or *C. elegans*^15^. In addition to the wound-healing process, ROS signaling also contributes to complex regenerative processes such as *Drosophila* gut^16^, *Drosophila* wing^17,18^, *Xenopus* tadpole tail^19^, juvenile axolotl tail^20^, adult zebrafish fin^21^.

At high concentrations, H_2_O_2_ is able to trigger cell death either by activating the apoptosis signal-regulated kinase ASK^18,22,23^ and/or through MAPK phosphatase inactivation and subsequent JNK activation^17,24–26^. Here, we investigated the ROS molecules produced in *Hydra* bisected at mid-gastric position, and the role they play during apical and basal regeneration. We focused on three aspects: (i) the spatial and temporal production of ROS signals during *Hydra* regeneration; (ii) the enzymatic activities that regulate ROS metabolism in intact and regenerating animals; (iii) the role of ROS signals on wound healing, injury-induced cell death and apical regeneration. We show that O_2_^.-^ and H_2_O_2_ fulfill the criteria of immediate injury-induced signals, O_2_^.-^ being required for wound healing and high levels of H_2_O_2_ for apical regeneration.

## RESULTS

### Mathematical modelling prediction of an immediate injury-induced signal in Hydra regenerating tips

We hypothesize that animals submitted to mid-gastric bisection release or produce a diffusing signal, yet unidentified and notated *U*, which would activate the MAPK pathway. We tested the hypothesis by developing a mathematical model comprising three components: the cells, the extracellular signaling and the intracellular signaling (**Fig. 1B**). The peculiar space response of the interstitial cells following the amputation-induced apoptosis indicates that the mathematical model should incorporate the space dimension. Since the only relevant direction is the distance perpendicular to the amputation plane (**Fig. 1A**) the mathematical model involves not just ordinary differential equations (ODE) but also one-dimensional partial differential equations (PDEs). In the absence of quantitative information on the interaction between cells and signaling, the model is constrained by a number of plausible assumptions. According to the model, mid-gastric bisection releases or produces the signal *U*, which diffuses and undergoes lytic degradation (**Eq. 1**: 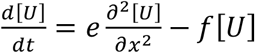). For simplicity, it is assumed that the signal is linearly degraded, which means that the effective concentration of the signal would be lower than the *Km* of a putative enzyme responsible for its degradation.

The link between the signal *U* and the MAPK pathway is proposed as follows: the signal *U* activates the phosphorylation of the inactive forms of the MAPK pathway (*M*_*i*_). It is considered that the reaction of activation is bilinear with the concentration of the substrate, the non-phosphorylated or inactive form of the enzymes (*M*_*i*_) and the signal (*U*) (**Eq. 2** in the Materials and Methods section). The backward reaction rate is assumed linear in the phosphorylated or activated signal (Ma, **Eq. 2**: 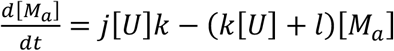).

The activated form of MAPK (*M*_*a*_) induces apoptosis by triggering the apoptotic cascade (via caspases) in the interstitial cell lineage. The simplest way to model this process is to assume that the rate of change of the apoptotic cell density is proportional to the interstitial cell density and the concentration of activated MAPK. It was previously described that three stages of apoptotic cells appear sequentially after mid-gastric bisection in *Hydra*^7^. However, the first two stages (formerly called early and advanced apoptotic cells) are kinetically equivalent suggesting that both stages could be modeled as a single stage hereafter constituted by the early apoptotic cells (*A*_*e*_). Hence, in the model *M*_*a*_ induces apoptosis by linearly decreasing the density of interstitial cells (*I*), which is the source of the density of the early apoptotic cells (*A*_*e*_, **Equations 3** and **4**). That is, the number of interstitial cells (*I*) is reduced and the number of early apoptotic cells (*A*_*e*_) is augmented in the same proportion (**Eq. 3**: 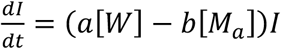 and **Eq. 4**: 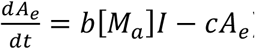). These cells are in turn linearly transformed in late apoptotic cells, which are also linearly depleted (*A*_*l*_, **Eq. 5**: 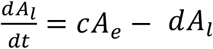). The early apoptotic cells release Wnt3 (*W*), which diffuses and is linearly degraded (*W*, **Eq. 6**: 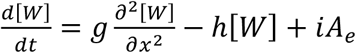) while it promotes the mitotic division of the neighbor interstitial cells (**Eq. 3**).

The model was fitted to previous experimental data of cell dynamics after mid-gastric bisection as shown in (**Fig. 1C**). The density of interstitial, early apoptotic and late apoptotic cells was calculated at different time points in two space regions, Region-1, i.e. 0 - 100 μm close to the amputation plane, and, Region-2 corresponding to the 100 - 200 μm area underneath. By using the procedure detailed in Supplementary Methods, the model was successfully fitted to the experimental time courses of densities of interstitial, early apoptotic and late apoptotic cells in Region 1 and 2 reported in regenerating *Hydra* (**Fig. 1C, 1D, Supplementary FigS1**). The predicted pattern of the hypothetical signal *U* is localized within approximately 150 µm from the amputation plane and it vanishes after one hour (**Fig. 1D**). Although more diffuse, the model-predicted pattern of Wnt3 (**Fig. 1D**) is also in agreement with the previously reported distribution in *Hydra* after amputation.

### Conservation of the SOD and catalase enzymatic machinery in Hydra

ROS metabolism arose early in evolution with H_2_O_2_ functioning as a cell-autonomous or non-cell-autonomous second messenger in bacteria, plants and metazoans^27^. There are two major sources of superoxide (O_2_^.-^) production in the cell, either by the mitochondrial Electron Transport Chain (ETC) complexes^28^ or by the membrane NADPH Oxidase (NOX) enzymes^29,30^ (**Fig. 2A**). O_2_^.-^ is a highly reactive and short-lived molecule that upon dismutation by superoxide dismutases (SOD) is transformed into H_2_O_2_, a relatively stable ROS molecule even when extra-cellular. Before initiating an analysis of ROS signaling in *Hydra*, we performed a structural and phylogenetic analysis of the components of the enzymatic machinery involved in ROS metabolism and the results, summarized in **Fig. 2** are shown in **Supplementary Fig. S2-S7**.

**Fig. 2.**
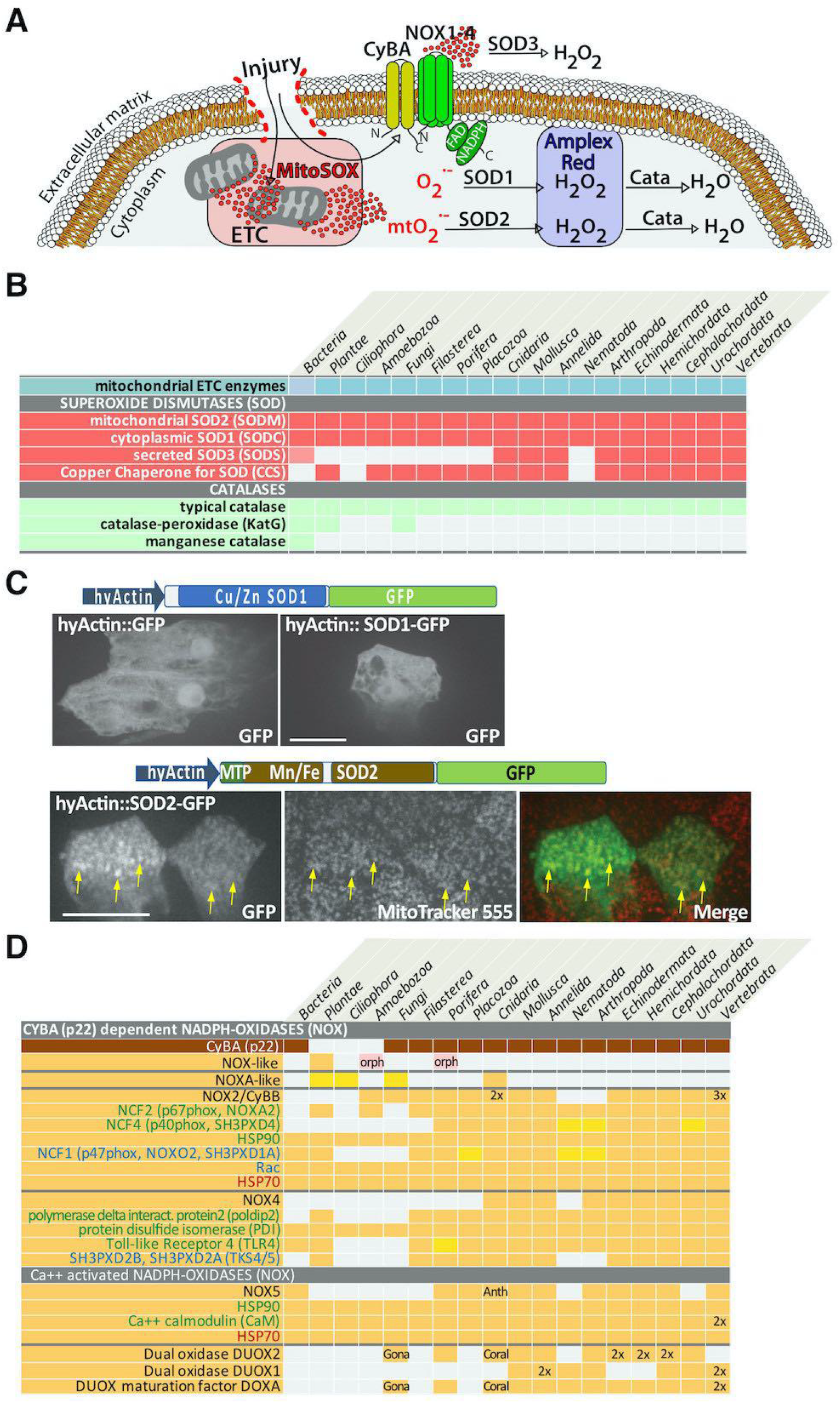
Evolutionary conservation of the enzymatic machinery involved in ROS metabolism. **(A)** Schematic view of superoxide (O_2_^.-^) and H_2_O_2_ production, either from the mitochondria (mtO_2_^.-^) via the electron transport chain (ETC), or from the membrane via NADPH Oxidase (NOX) activity; mtO_2_^.-^ is detected with MitoSOX and H_2_O_2_ with Amplex Red. **(B)** Schematic view of the phylogenetic distribution of the SOD enzymes. Accession numbers are listed in **Table-S1**. Structural and phylogenetic analyses of SOD sequences are given in **Supplementary Figures S2, S3** and **S4. (C)** Live imaging of eESCs transiently expressing the *HyActin*::SOD1-GFP and *HyActin*::SOD2-GFP constructs two days after electroporation. Note the cytoplasmic and mitochondrial (yellow arrows) localization of SOD1-GFP and SOD2-GFP respectively. Scale bars: 25 µm. **(D)** Schematic view of the phylogenetic distribution of the NOX enzymes and the components of the NOX2/CYBB, NOX4 and NOX5 complexes. Structural and phylogenetic analyses of NOX sequences are available in **Supplementary Figures S5, S6** and **S7**. Accession numbers of NOX sequences are listed in **Table-S2**. The NOX2/CYBB sequence duplicated in the last common ancestor of eumetazoans to give rise to the NOX4 family, while NOX1 and NOX3 are vertebrate-restricted NOX2/CYBB paralogs (3x). Anth: sequences only found in anthozoans among cnidarians; Coral: sequences only found in some corals; Gona: sequences only found in the fungus *Gonapodya prolifera*; orange background: orthologous or highly related sequence; pink background: orphan sequence (orph); yellow background: divergent sequence.

In vertebrates, three gene families encode the superoxide dismutases, *SOD1* and *SOD3* that encode the Copper Zinc-dependent SOD (CuZnSOD) enzymes, which are active in the cytoplasm and the extra-cellular space respectively, and *SOD2* that encodes the manganese-dependent SOD (MnSOD) enzyme, localised in the mitochondria^31,32^ (**Fig. 2, Supplementary Fig. S2**). *SOD1* and *SOD2* are found in Bacteria, Plantae, Fungi, Amoebozoa (*Dictyostelium*), Filasterea (*Capsaspora*) and Metazoa, whereas *SOD3* is only found in Cnidaria and Bilateria, suggesting a duplication of *SOD1* in the last common ancestor (LCA) of eumetazoans (**Fig. 2B, Table-S1, Supplementary Fig. S3**). The Copper Chaperone of SOD (CSS) is a factor necessary for Copper integration into SOD1^33^, *CSS* genes are found in Fungi, Plantae, Filasterea and Metazoa but not in Bacteria indicating an origin in the eukaryote LCA (**Supplementary Fig. S3A**).

*Hydra* expresses orthologs to each of these *SOD* genes (**Table-S1, Supplementary Fig. S4**), with *SOD1* and *SOD2* expressed at higher levels in the central body column, ubiquitously in the three stem cell populations but no temporal regulation during regeneration. By contrast, *SOD3*, which encodes a signal peptide, is expressed at higher levels at the apical and basal extremities of intact animals and almost exclusively in gESCs; *SOD3* is up-regulated during the early phase of apical and basal regeneration, subsequently maintained at high levels of expression (**Supplementary Fig. S4D**). To verify the expected cellular localization of SOD1 and SOD2, we produced and transiently expressed hyACT:SOD1-GFP and hyACT:SOD2-GFP in *Hydra* cells and found two distinct sub-cellular localizations, predominantly cytoplasmic for SOD1 and mitochondrial for SOD2 as anticipated from their respective structure (**Fig. 2C**). As ROS-detoxifying enzyme, *Hydra* expresses a single *catalase (cata)* gene (**Supplementary Fig. S2F**), at high levels along the body axis and predominantly in ESCs, with limited modulations during regeneration (**Supplementary Fig. S4**).

### Three CYBA-dependent NADPH-oxidases expressed in Hydra

Concerning the NADPH Oxidase (NOX) enzymes, the vertebrate NOX1, NOX2/CYBB, NOX3, NOX4 are known to be active only through their interaction with the critical membrane Cytochrome b-245 light chain (CYBA), that associates with their Cytochrome b-245 heavy chain (CYBB) domain to form functional NADPH oxidases and generate superoxide; by contrast the NOX5, DUOX enzymes are Ca^2+^ dependent^34–36^. In *H. vulgaris*, we identified orthologs to NOX2/CYBB, NOX4 and NOXA-like families, but no NOX5 and DUOX orthologs although present in two anthozoan species (**Fig. 2D, Supplementary Fig. S5**). The putative NOX2/CYBB, NOX4 and NOXA-like proteins exhibit a similar structure, a Ferric oxidoreductase domain in the N-moiety and a Nox-Duox-like_FAD_NADP domain in the C-moiety (**Supplementary Fig. S5B, S5C**).

The phylogenetic analysis of metazoan and non-metazoan NOX-related sequences identifies six NOX families (**Fig. 2D, Table-S2, Supplementary Fig. S6**): (1) the *NOX5* family found in Bacteria, Porifera, few coral cnidarian species (*Dengi, Xenia*), Placozoa (*Trichoplax*) and Bilateria including Vertebrata that appears monophyletic; (2) a *NOX-like* family present only in Plantae; (3) the *NOXA-like* family present in Ciliaphora (*Tetrahymena*), Fungi, Viridiplantae (green algae), Cnidaria but lost in Bilateria; (4) the *NOX2/CYBB* family that encodes the Cytochrome-b-245 heavy chain-like, present in Amoebozoa, Fungi, Filasterea and Metazoa, implying an origin that predates Unikonts; we also confirmed that *NOX1* and *NOX3* are *NOX2/CYBB* paralogs restricted to vertebrates; (5) the *CYBB/NOX2*-related *NOX4* family found only in Metazoa, implying a duplication event in the LCA of metazoans; (6) the *DUOX* family that encodes proteins characterized by an EF-hand domain and Ca-binding motifs, found only in Metazoa with two distinct sub-families, DUOX2 present in most metazoan phyla including Porifera, few cnidarian species, Annelida, Arthropoda, Mollusca, Echinodermata, Hemichordata, Cephalochardata, Urochordata but absent in Vertebrata, and DUOX1 restricted to Bilateria including Vertebrata suggesting a duplication of DUOX2 in the LCA of bilaterians (**Supplementary Fig. S6C**). We also verified through reciprocal blast hits that components of the NOX2/CYBB, NOX4 and NOX5 complexes^35^ are expressed in metazoans (**Fig. 2D**).

The analysis of the expression patterns of the *H. vulgaris NOX* genes was deduced from the RNAseq profiles^37^, showing along the body axis an expression predominantly apical for *NOXA-like, NOX2/CYBB* and *CYBA*, predominantly basal for *NOX4* (**Supplementary Fig. S7A**). All four genes are almost exclusively expressed in epidermal ESCs and their expression levels is not modified after elimination of interstitial cells (**Supplementary Fig. S7B, S7C**). During the first two days of regeneration, *NOXA-like* expression is not modulated, while *NOX2/CYBB* and *CYBA* are up-regulated in the immediate phase, reaching a plateau level after 8 hours. At 36 hours post-amputation (hpa), in the late phase of basal regeneration, *CYBA* and *NOX4* are up-regulated.

### Symmetrical levels of mtO_2_^.-^ versus asymmetrical levels of H_2_O_2_ in apical and basal regenerating tips

To test whether ROS molecules would fit the expected features of the injury-induced *U* signal predicted by the model (**Fig. 1D**), we applied two different ROS detection methods: the MitoSOX superoxide indicator that emits a red fluorescent signal when oxidized by mitochondrial superoxide (mtO_2_^.-^)^38^ and the Amplex UltraRed reagent to quantify H_2_O_2_ levels, a method widely-used in human leukocytes^39^, human cancer cells^40^, mice^41^, *D. melanogaster*^42^ and plants^43^. We applied these methods on wounded and regenerating *Hydra* (**Fig. 3A-H**) and to identify the cells that produce mtO_2_^.-^, we used transgenic lines that express the LifeAct-GFP reporter in eESCs and gESCs respectively^44,45^. We detected mtO_2_^.-^ signals in both apical- and basal-regenerating (AR, BR) tips, mostly in gastrodermal ESCs as confirmed by the orthogonal maximum projections along the z-y axis (**Fig. 3B**), with ∼80% of MitoSox dots in the gESCs in both AR and BR tips (**Fig. 3C**).

**Fig. 3.**
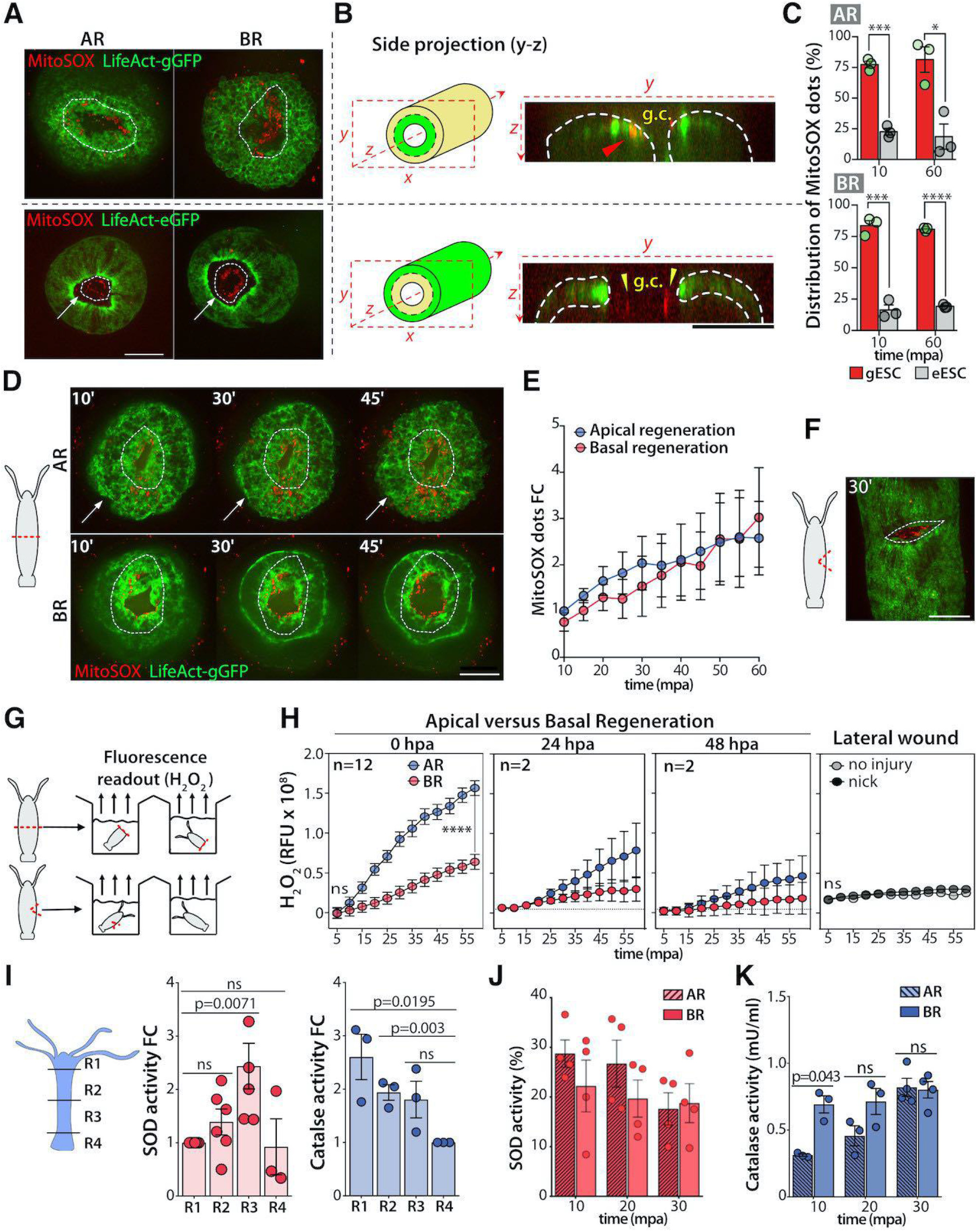
Production of mitochondrial superoxide (O_2_^.-^) and hydrogen peroxide (H_2_O_2_) in intact, injured and regenerating Hv_AEP2 animals. **(A)** Top projection views of apical- and basal-regenerating tips (AR, BR) from transgenic animals constitutively expressing LIfeAct-GFP in gastrodermal (LifeAct-gGFP) or epidermal (LifeAct-eGFP) ESCs, labeled with MitoSOX (red) 20 minutes post-amputation (mpa). White dotted lines indicate wounded areas; white arrows point to the wound edges enriched with F-actin cables only visible in the epidermal epithelial cells. **(B)** Side projection views of live AR halves from LifeAct-gGFP and LifeAct-eGFP animals imaged at 20 mpa. **(C)** Distribution of MitoSOX dots between eESCs and gESCS imaged on AR and BR tips from LifeAct-eGFP animals at 10 and 60 mpa. **(D, E)** Detection and quantification of MitoSOX dots in AR and BR tips from LifeAct-gGFP animals. White dotted lines as above, white arrows point to the mesoglea. **(F)** MitoSOX+ dots detected in a lateral wound of a LifeAct-eGFP animal (see **Supplementary Fig.S8**). White dotted line indicates the lateral nick area. Scale bars in C, D, F, H = 200 µm. **(G)** Scheme depicting the Amplex Red procedure used to quantify H_2_O_2_ levels in AR and BR halves and in intact or wounded animals. (**H**) H_2_O_2_ production measured at indicated conditions for 60 minutes, n: number of independent experiments, ****= p-value < 10-^4^, unpaired t-test. **(I)** Superoxide dismutase (SOD) and catalase (cata) enzymatic activities in four distinct regions dissected along the body axis of intact animals. SOD values are normalized on the R1 value and cata values on the R4 value. **(J, K)** SOD and catalase activities in AR and BR tips dissected at 10, 20 and 30 mpa. Each dot represents value from an independent experiment. Error bars correspond to SEM values; ns: not significant.

The analysis of the temporal and spatial variations in mtO_2_^.-^ levels during the immediate phase of apical and basal regeneration shows that MitoSox signals initially detected as early as 10 minutes after mid-gastric bisection, form a ring-like structure at the bisected planes of both upper and lower halves (**Fig. 3D**). Then, during the first hour post-amputation, mtO_2_^.-^ progressively accumulates in the AR- and BR-tips. Since MitoSOX is a cumulative type of cellular dye, we could only evidence a rather sustained increase. When we measured the total number of MitoSOX dots every five minutes from 5 to to 60 minutes post-amputation (mpa) in AR and BR tips, we recorded similar levels of mtO_2_^.-^ on each side of the cut, reflecting a similar up-regulation in both contexts (**Fig. 3E**). Next, we monitored the mtO_2_^.-^ levels in wounded but non-regenerating tissues: We injured LifeAct-eGFP animals laterally, making a notch with a scalpel at mid-body position and detected MitoSox dots (**Supplementary Fig. S8)**. As in AR and BR tips, the MitoSoxdots are located outside the epidermal GFP+ layer, consistent with a source in the gastrodermal epithelial cells (**Fig. 3F**).

Next, we quantified the levels of H_2_O_2_ in the upper and lower halves using the Amplex UltraRed assay. We did not evidence any significant difference in H_2_O_2_ production between apical and basal regeneration during the first 10 minutes, but soon after, we recorded a significant increase in H_2_O_2_ levels from AR halves, reaching at 60 mpa a three-fold difference with the level measured in BR halves (**Fig. 3G, 3H**). In animals injured with a lateral notch, H_2_O_2_ levels remain stable, indicating that, in contrast to mtO_2_^.-^, the increase of H_2_O_2_ level is regeneration-specific (**Fig. 3H**). We speculated that the asymmetric H_2_O_2_ levels in AR and BR tips might reflect an asymmetric constitutive distribution of enzymatic activities involved in ROS production and/or ROS degradation.

### Constitutive asymmetrical SOD and catalase activities along the body axis

To measure the global activities of the SOD and catalase enzymes in intact animals, we dissected four regions along the body axis of *Hv_AEP2* polyps, (i) apical, (ii) upper body column, (iii) lower body column and (iv) basal, and prepared protein extracts from each of them (**Fig. 3I**). We recorded maximal levels of SOD activity in the lower body column, significantly higher than in the apical region, whereas the catalase activity appears apical-to-basal graded, lower in the lower body column when compared to the apical region and undetectable in the peduncle/basal region. All together, these analyses show that in homeostatic conditions SOD activity appears maximal basally, whereas catalase activity is maximal apically, with an inverted apical-to-basal graded distribution along the body axis. These asymmetric homeostatic distributions along the body axis potentially explain how, after bisection, higher levels of H_2_O_2_ are obtained in the lower half, i.e., more production and less degradation, opening the possibility of distinct ROS signaling levels on each side of a mid-gastric cut (**Fig. 3I**).

To identify how asymmetrical H_2_O_2_ levels rapidly form in AR and BR tips, we measured in *Hv_AEP2* the SOD and catalase activities in regenerating tips dissected at early time-points. According to the constitutively asymmetric distribution of catalase activity along the body axis of intact animals, we were anticipating a higher SOD activity and a lower catalase activity in AR than in BR tips. Indeed, we found SOD activity slightly higher in AR than in BR tips at 10 and 20 minutes post-amputation (mpa) although this difference is not significant (**Fig. 3J**) and catalase activity significantly higher in BR tips than in AR tips at 10 mpa (**Fig. 3K**).

In summary of these two sections, the two ROS molecules identified as immediate injury-induced signals in bisected *Hydra* exhibit distinct regulations on each side of the cut, with mtO_2_^.-^ produced symmetrically by gESCs on either side of the bisection plane, and H_2_O_2_ submitted to asymmetric regulation. Despite the lack of a direct method to quantify membrane superoxide (mbO_2_^.-^) in *Hydra* tissues, we suspect NOX enzymes that produce mbO_2_^.-^to contribute to the asymmetric H_2_O_2_ production.

### Superoxide scavenging and oxidase inhibition lower H_2_O_2_ production

To elucidate the respective roles of mtO_2_^.-^ and H_2_O_2_ during regeneration, we first tested a series of pharmacological inhibitors like the cell-permeable iron chelator Tiron, commonly used as a superoxide scavenger^46–48^, or the flavoprotein inhibitor diphenyleneiodonium (DPI) that prevents the activity of several oxidases including the NOX and ETC enzymes^35^, or the specific NOX inhibitor VAS2870, or the catalase inhibitor 3-aminotriazole (ATZ)^49^ (**Fig. 4A**). For all these drugs, we performed toxicological tests to select the highest non-toxic concentrations, 15 mM for Tiron, 10 µM for DPI and 75 mM for ATZ (**Supplementary Fig. S9**). For VAS-2870, we never detected any toxic or biological effect and concluded that this drug is not active in *Hydra*.

**Fig. 4.**
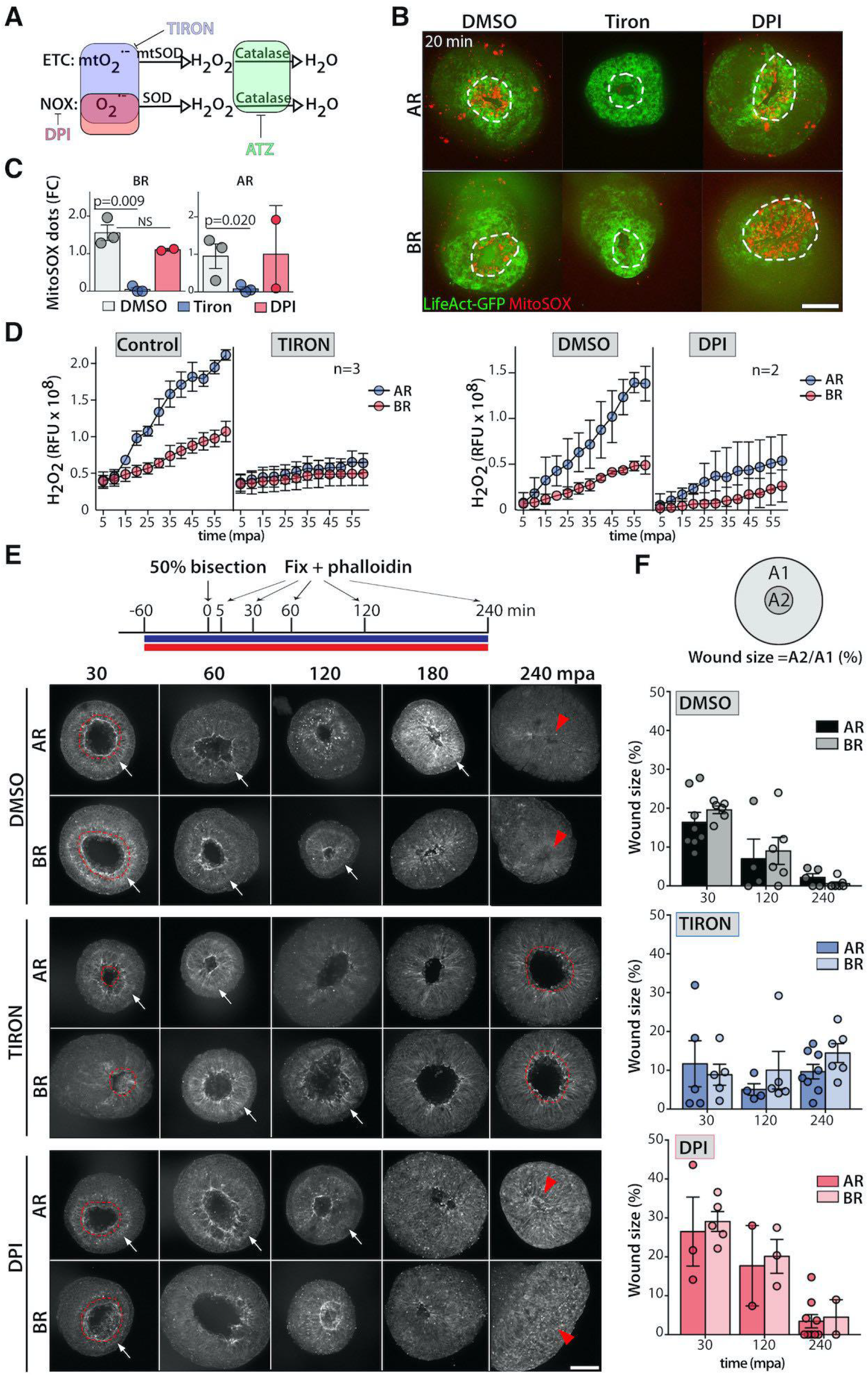
Pharmacological scavenging of superoxide inhibits wound healing. **(A)** Schematic view of the antioxydant activity of the Tiron and DPI drugs, Tiron acting as a superoxide (O_2_^.-^) scavenger and DPI inhibiting intracellular NADPH Oxidase (NOX) enzymes. **(B, C)** Detection of mitoO_2_^.-^ (red) at 20 mpa in AR and BR halves of animals either left untreated (DMSO) or exposed to Tiron or DPI for one hour before amputation. White dotted line indicates the wounded area. **(D)** H_2_O_2_ levels in AR and BR halves of animals exposed to Tiron or to DPI. **(E)** Monitoring of the wound closure process after mid-gastric bisection in DMSO, Tiron- or DPI-treated animals stained with phalloidin at indicated time points. Red dotted line indicates the wounded area, red arrowheads the healed wounds, white arrows the mesoglea. **(F)** Quantification of wound closure. At each time-point, the wound size was determined as the ratio between the gastric opening (A2) over the total surface (A1). Each dot corresponds to one animal wound. Scale bar: 200 µm. See **Supplementary Figure S10**.

Next, we measured the MitoSOX signal in *Hv_AEP2* animals exposed to Tiron for one hour prior to amputation (**Fig. 4B-C**). As expected, we noted in Tiron-treated animals a drastic decrease in MitoSOX signal in AR and BR tips, confirming that superoxide scavenging efficiently lowers mtO_2_^.-^ levels. We also found the H_2_O_2_ levels significantly reduced, confirming that Tiron efficiently scavenges all sources of O_2_^.-^ (**Fig. 4D**). In contrast, in bisected DPI-treated animals, we did not record any change in mtO_2_^.-^ levels (**Fig. 4B, 4C**) but we found the H_2_O_2_ levels significantly reduced (**Fig. 4D**), indicating that DPI mostly affects NOX enzymes and mbO_2_^.-^ production. These results indicate that NOX enzymes also contribute to H_2_O_2_ production in regenerating tips.

### Pharmacological and genetic modulations of ROS levels impact on wound healing

To monitor the effect of lowering mtO_2_^.-^ or mbO_2_^.-^ levels on wound closure process, we treated the animals with Tiron or DPI as above, collected the regenerating halves at five distinct time-points between 30 and 240 mpa, fixed and labeled them with Phalloidin, imaged the wound plane and measured the wound size (**Fig. 4E, 4F, Supplementary Fig. S10**). In untreated animals, the wound closure is rapid, symmetrical in AR and BR halves, with a two-fold decrease in the wound size between 30 and 120 mpa, and a complete closure recorded in 100% of animals at four hpa. In Tiron-treated animals, the wound remains open up to four hpa while in DPI-treated animals there is no delay in wound healing. These results suggest that wound healing relies on mtO_2_^.-^ rather than on mbO_2_^.-^ production.

Next, we silenced *SOD1, SOD2* or *SOD3* by electroporating five times the animals with siRNAs and found the transcript levels down-regulated by 55% for *SOD1* and *SOD2*, by 40% for *SOD3* (**Fig. 5A, 5B, Table-S3**). In *SOD1*(RNAi) animals we recorded a four-fold increase in mtO_2_^.-^ levels at 10 mpa, a level maintained elevated over 70 mpa (**Fig. 5C**), and H_2_O_2_ levels dramatically decreased in both AR and BR halves (**Fig. 5D**). In *SOD2*(RNAi) and *SOD3*(RNAi) animals, H_2_O_2_ levels were also decreased but to a lesser extent (**Fig. 5D**). These results reflect the efficiency of RNAi-induced inhibition of SOD activity. Next, we tested the impact of high mtO_2_^.-^ levels on wound healing: We recorded a faster wound healing process in both AR and BR tips of *SOD1*(RNAi) animals, as observed at 120 mpa (**Fig. 5E, 5F, Supplementary Fig. S11A**). We obtained a similar result in *SOD2*(RNAi) and *SOD3*(RNAi) animals (**Fig. 5G-5J, Supplementary Fig. S11B, S11C**), confirming that superoxide plays a positive role in wound healing.

**Fig. 5.**
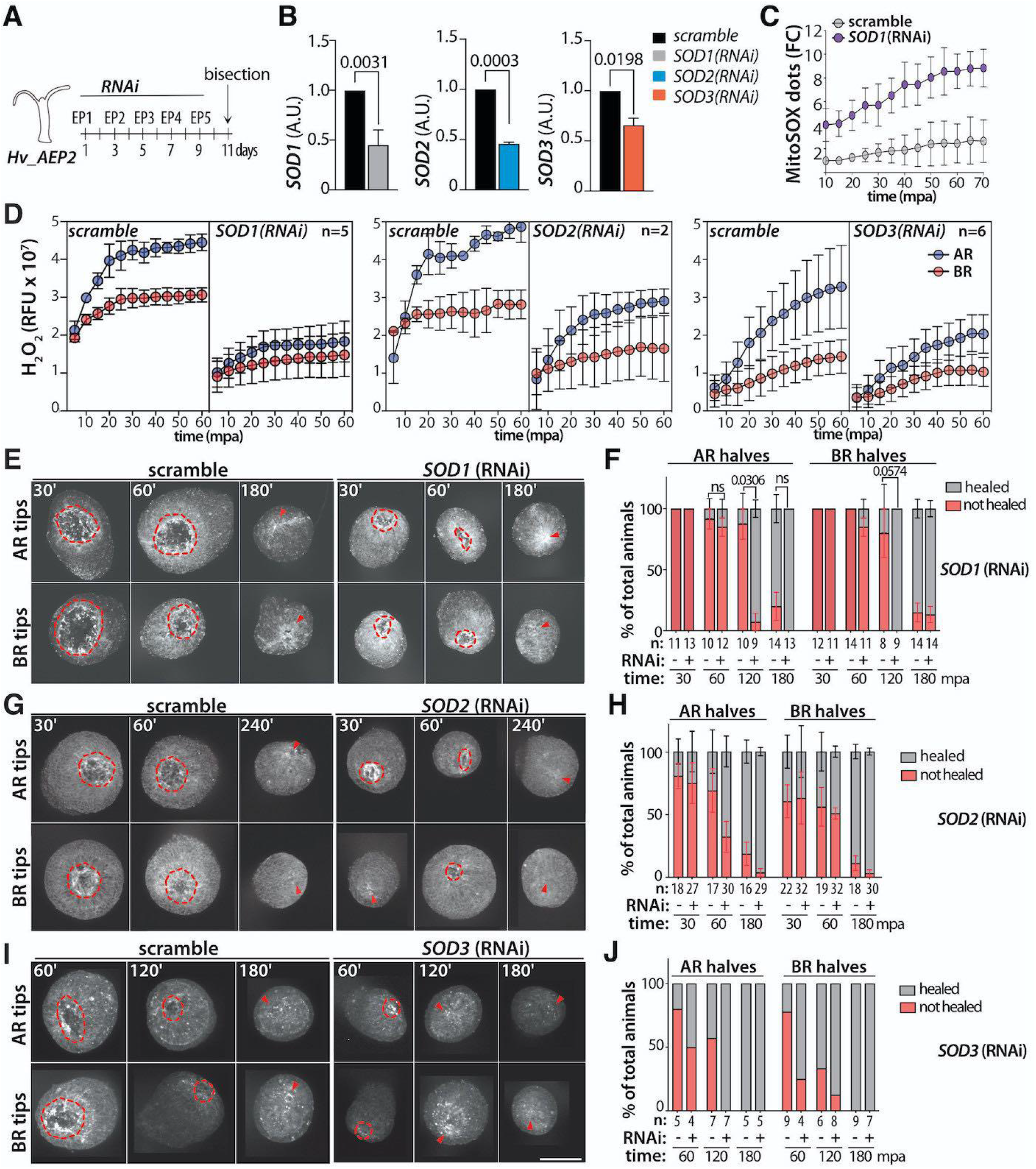
Wound healing in Hv_AEP2 animals knocked-down for SOD1, SOD2 or SOD3. **(A)** Schematic view of the RNAi procedure relying on 4 or 5 successive electroporations (EP) of siRNAs and an amputation two days after the last EP. **(B)** Q-PCR analysis of *SOD1, SOD2* and *SOD3* transcript levels two days after EP5 for *SOD1(RNAi)*, two days after EP4 for *SOD2(RNAi)* and *SOD3(RNAi)*. **(C)** Quantification of MitoSOX dots in AR wounds of animals treated and bisected as in B and pictured from 10 to 70 mpa. **(D)** H_2_O_2_ levels in AR and BR halves of animals electroporated with scramble, *SOD1, SOD2* and *SOD3* siRNAs. Imaging **(E, G, I)** and quantification **(F, H, J)** of the wound healing process in AR and BR halves of animals knocked-down for *SOD1* **(E, F)**, *SOD2* **(G, H)**, *SOD3* **(I, J)**. Animals were bisected two days after EP5 for *SOD1*(RNAi), two days after EP4 for *SOD2(RNAi)* and *SOD3(RNAi)*. Scale bar = 200 µm. Red dotted line indicates the open wounded area, red arrowheads the healed wounds. Error bars correspond to SEM values. Statistics were done via unpaired t-tests, ns= non significant. See **Supplementary Figure-S11**.

### Pharmacological and genetic modulations of ROS levels impact on apical regeneration

To investigate the role of ROS signaling on *Hydra* apical regeneration, we monitored the kinetics of apical regeneration in animals exposed to Tiron or DPI, given either over the immediate-early phase (from one hour before bisection up to 24 hpa), or during the early-late phase of regeneration (from 24 up to 48 hpa) (**Fig. 6A, Supplementary FigS12**). While the immediate-early Tiron treatment significantly delays apical regeneration, the early-late Tiron treatment does not, a result that highlights the important role(s) played by O_2_^.-^ in the immediate phase of regeneration. Similarly, DPI applied during the immediate-early phase greatly delays apical regeneration whereas a treatment given after 24 hpa does not affect it. When we focused on the 50% apical regeneration cutoff value in the early treatment experimental group, we noticed that the delay for Tiron-treated animals compared to control is around 24 hours. But in the case of DPI-treated animals this is significantly longer, around 70 hours (≈3 days). This difference is not observed in the late treatment group. Additionally, approximately 65% of DPI-early treated animals do not fully regenerate their heads even at 140 hours after the cut (**Fig. 6A, 6B Supplementary FigS12**). We also tested the impact of knocking-down *SOD1* on apical regeneration. Despite higher mtO_2_^.-^ and lower H_2_O_2_ levels, we did not record any modification in the regeneration kinetics or the morphology of the regenerated structure when compared to the control scramble(RNAi) animals (**Supplementary Fig. S13**). These results indicate that a partial *SOD1* silencing does not efficiently reduce H_2_O_2_ levels over several days when compared to DPI treatment.

**Fig. 6.**
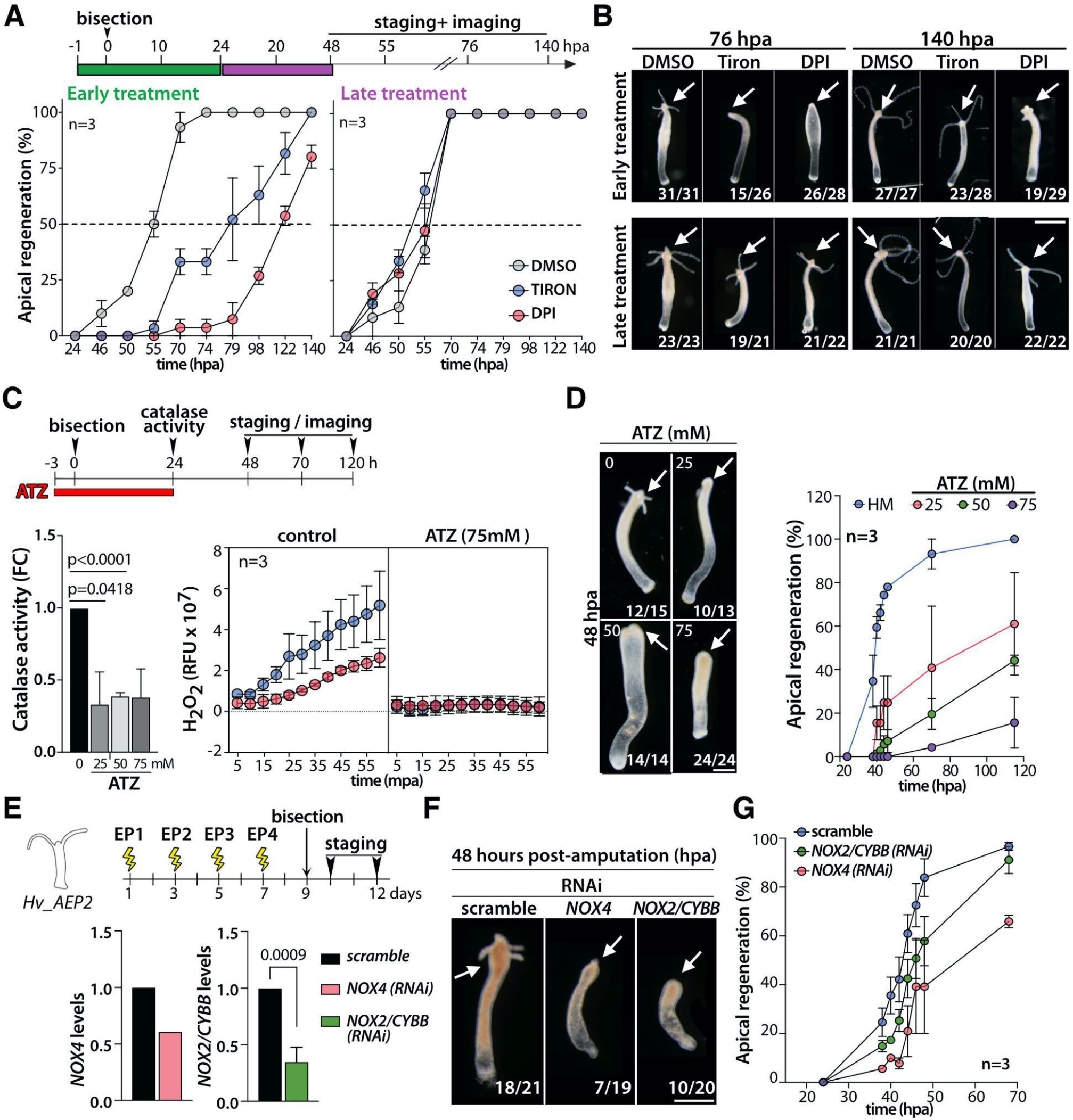
Impact of pharmacological and genetic modulations of ROS metabolism on apical regeneration. **(A)** Kinetics of apical regeneration in animals exposed to Tiron or DPI during the early (green bar) or early-late (purple bar) phases of regeneration. **(B)** Apical morphology of Tiron- and DPI-treated regenerating animals pictured at 76 and 140 hpa. Arrows indicate the apical regenerating extremity. **(C)** Catalase activity and H_2_O_2_ levels after 27 hours ATZ treatment as depicted in the scheme. **(D)** Dose-dependent inhibition of apical regeneration in ATZ-treated animals, pictured here at 48 hpa. **(E)** Scheme depicting the RNAi procedure to knock-down *NOX4* or *NOX2/CYBB*, and expression levels measured by Q-PCR on the bisection day. **(F)** Delayed apical regeneration in *NOX4(RNAi)* or *NOX2/CYBB(RNAi)* animals pictured at 48 hpa. **(G)** Kinetics of apical regeneration in scramble, *NOX2/CYBB(RNAi)* and *NOX4(RNAi)* animals. In panels A, C, D, E, G, error bars corresponding to SEM values. In B, D, F, scale bars = 500 µm. See **Supplementary Figures S12** (panel B), **S14** (panel D) and **S15** (panel G).

We also tested the impact of catalase inhibition by treating the animals with ATZ for 3 hours before amputation and 24 hours after. This treatment decreases by 60% the level of catalase activity (**Fig. 6C**), and a 3-hour ATZ treatment before amputation suffices to prevent any up-regulation of H_2_O_2_ levels immediately after bisection (**Fig. 6C**). This result is surprising as in the absence of H_2_O_2_ catabolic activity, we would rather expect an increase in H_2_O_2_ levels. We also noted a dose-dependent inhibition of apical regeneration in ATZ-treated animals (**Fig. 6D, Supplementary Fig. S14**), consistent with a positive role of H_2_O_2_ in apical regeneration. However, a reduction of *catalase* expression after siRNAs did not modify the regeneration kinetics or the morphology of the regenerated structure (**Supplementary Fig. S13**), again pointing to the inefficiency of a partial silencing.

The predominant expression of *NOX* genes in eESCs suggests a role for these enzymes in H_2_O_2_ production. To test the function of NOX enzymes in regenerating *Hydra*, we designed siRNAs targeting *NOX2/CYBB, NOX4* or *NOXA-like* and electroporated them four times in intact *Hv_AEP2* animals, which were subsequently bisected two days after the last electroporation (**Fig. 6E**). On the bisection day, we measured a ∼70% reduction of *NOX2/CYBB* expression and ∼50% for *NOX4* (**Fig. 6E**). In three independent experiments, we observed a 24-hour delay in apical regeneration after *NOX2/CYBB*(RNAi). At 48 hpa, the apical regeneration was successful in 86% animals after scramble (RNAi), 63% animals after *NOX4(*RNAi), 62% animals after *NOXA-like*(RNAi), and 50% animals after *NOX2/CYBB(*RNAi) (**Fig. 6F, Supplementary Fig. S15**). At 72 hpa, apical regeneration was achieved in only 60% *NOX4*(RNAi) animals (**Fig. 6G**). We noted that pharmacological treatments (Tiron, DPI, ATZ) have a more severe impact on apical regeneration compared to *SOD*(RNAi) or *NOX*(RNAi) experiments where gene silencing is only partial.

The results from these wound healing and regeneration experiments suggest that high levels of H_2_O_2_ facilitate the immediate phase of apical regeneration. Also, the fact that Tiron and DPI differently impact wound healing and apical regeneration suggests that mtO_2_^.-^ and mbO_2_^.-^ play distinct roles during *Hydra* regeneration; mtO_2_^.-^ playing a key role on wound healing but a limited one on apical regeneration, and mbO_2_^.-^ being necessary to reach high H_2_O_2_ levels, essential for apical regeneration.

### ROS signaling supports CREB phosphorylation and cell death in apical-regenerating tips

Next, as H_2_O_2_ is a rather stable messenger molecule, able to cross membranes and promote paracrine signaling, we explored the possibility of paracrine ROS signaling between the gastrodermal and epidermal layers in regenerating *Hydra*. As support for this hypothesis, we noted that mtO_2_^.-^ is predominantly detected in gastrodermal ESCs, while cells submitted to injury-induced cell death belong to the interstitial lineage, located in the epidermal layer. To investigate whether injury-induced ROS signals impact MAPK activation, we tested CREB phosphorylation and injury-induced cell death in AR tips (**Fig. 7A)**. To test whether CREB phosphorylation is altered when ROS signals are scavenged, we first immunodetected the phosphorylated form of CREB (pCREB) in the AR tips of Tiron-treated animals. As previously shown^9^, we identified in control animals taken at 30 mpa numerous pCREB positive nuclei in the most superficial region of untreated lower halves, whereas pCREB positive nuclei are rare after Tiron treatment (**Fig. 7B**).

**Fig. 7.**
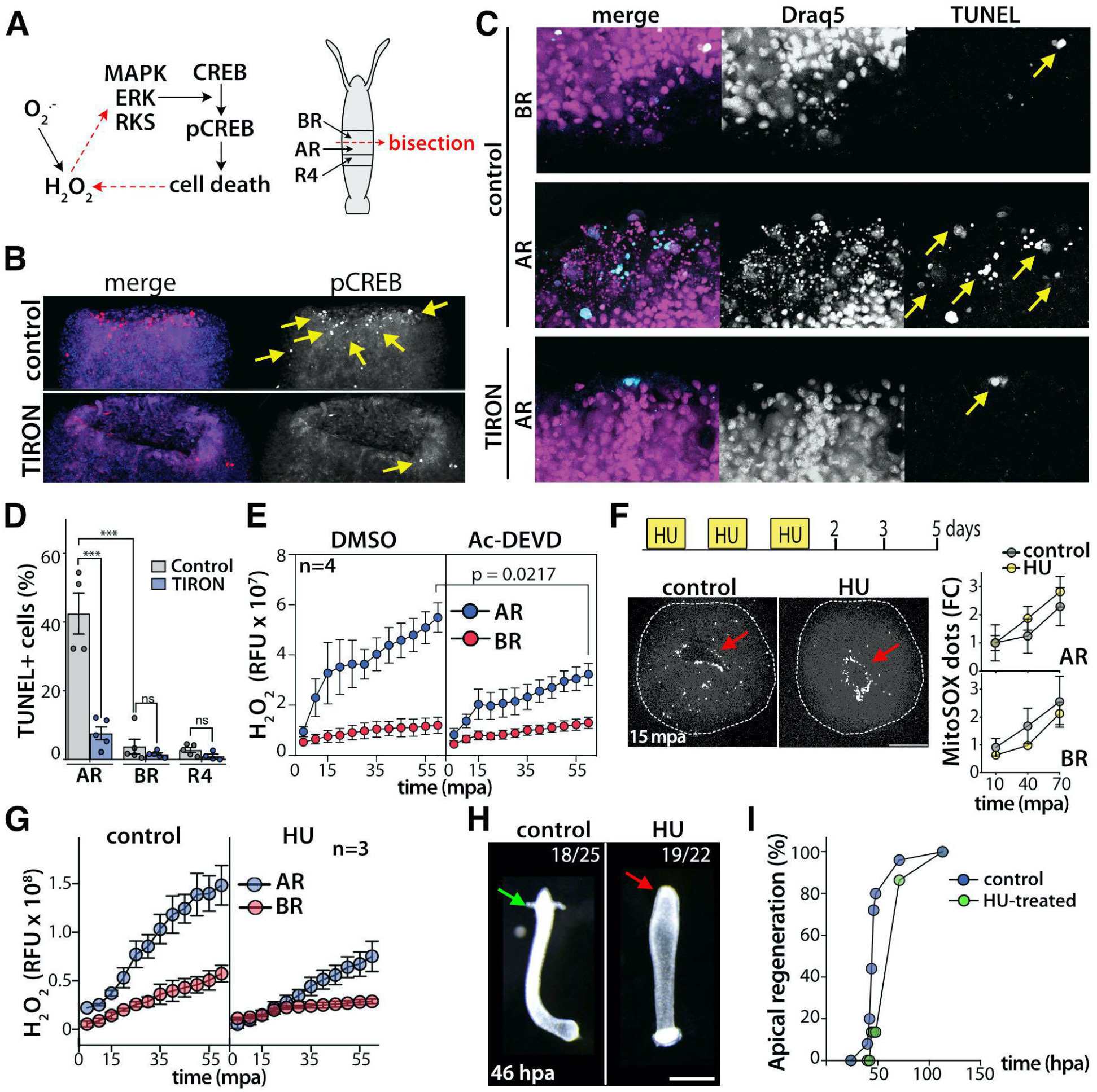
High levels of injury-induced H_2_O_2_ production in AR tips and the massive death of interstitial cells. **(A)** Putative regulation played by ROS signals on MAPK/CREB signaling in AR tips after mid-gastric bisection. Schematic view of the AR and BR tips, and R1 region in bisected *Hydra*. **(B)** Immunodetection of pCREB in AR tips from untreated (control) or Tiron-treated *Hv_Basel* animals fixed at 30 mpa. **(C)** Tiron-induced inhibition of cell death in AR halves of Tiron-treated *Hv_Basel* animals fixed at 30 mpa, labeled with TUNEL (arrows) and stained with Draq5. Note that nuclei are no longer fragmented and cells no longer TUNEL+ after Tiron treatment. **(D)** Three regions were dissected as in depicted in Fig1A. Graph showing the quantification of TUNEL positive cells in macerated regenerating tips dissected 30 min after bisection. *Hv_Basel* animals were either untreated (Control) or pulse-treated with 15 mM Tiron. Mean ± SEM, n= 4-5, p>0.01. One-way ANOVA with Tukey’s post hoc test, ****p <0.0001. **(E)** Impact of caspase inhibition induced by Ac-DEVD on H_2_O_2_ levels in AR and BR halves in *Hm_105* animals. **(F, top)** Scheme depicting HU treatment of *Hv_AEP2* animals to induce the elimination of interstitial cells. **(Bottom)** MitoSOX staining of control and HU-treated animals, bisected and imaged live at 15 mpa. The outer border of the polyp is shown with a white dotted line, MitSOX signals correspond to white dots (red arrow). Quantification of the MitoSOX signal is shown on the right. **(G)** H_2_O_2_ levels in AR and BR halves from control or HU-treated *Hv_AEP2* animals treated as in F and bisected on day 5. **(H)** Apical-regenerating animals imaged at 46 hpa, either untreated (green arrow) or HU-treated (red arrow). Scale bar: 500 µm. **(I)** Kinetics of apical regeneration of HU-treated *Hv_AEP2* animals. In panels D, E, F and G, error bars correspond to SEM values. See **Supplementary Figures S16** and **S17**.

To test the impact of ROS signaling on injury-induced cell death, we performed TUNEL labeling on AR and BR halves exposed or not to Tiron as described above, fixed 30 minutes after bisection (**Fig. 7C**). We noted a cell death zone containing TUNEL positive cells with fragmented nuclei in AR tips of control animals (**Fig. 7C**, middle panel, yellow arrows). Fragmented nuclei and TUNEL+ cells were rare in the BR tips (**Fig. 7C**, upper panel) and absent in AR tips from animals exposed to Tiron (**Fig. 7C**, lower panel). When we quantified the number of TUNEL positive cells, we found over 40% dying cells in AR tips of control animals at 30 mpa, in agreement with previous results^7^ and less than 10% after Tiron exposure (**Fig. 7D**). This result supports the hypothesis that ROS signaling contributes to injury-induced cell death on a paracrine mode, with H_2_O_2_ diffusing from gESCs to the epithelial layer and triggering there, interstitial cell death while epithelial cells resist to it. We also analyzed the impact of injury-induced cell death on ROS production. To do so, we treated animals with the caspase-3 inhibitor Ac-DEVD-CHO and measured ROS levels after bisection. We found the injury-induced increase in H_2_O_2_ levels in AR tips significantly lower, confirming a likely paracrine feedback loop between injury-induced cell death and ROS production (**Fig. 7E**). These results indicate that there are two sources of H_2_O_2_ supply: one from the gESCs and the other from the dying interstitial cells.

To investigate the putative role of H_2_O_2_ diffusing from interstitial dying cells, we depleted the animals from their interstitial cells by treating them with hydroxyurea (HU)^50^ (**Fig. 7F**). Briefly, HU treatment that blocks DNA replication, selectively kills in several days the fast-cycling cells of the interstitial lineage without affecting the gastrodermal or epidermal ESCs, which are highly resistant to cell death. We measured the levels of mtO_2_^.-^ and H_2_O_2_ in animals bisected 5 days after in a three-course HU treatment and we recorded similar levels of mtO_2_^.-^ in AR and BR tips from complete and interstitial-depleted animals indicating that interstitial cells do not contribute to mtO_2_^.-^ production (**Fig. 7F, Supplementary Fig. S16**). However, we recorded a lower injury-induced up-regulation of H_2_O_2_ levels in AR tips from HU-treated animals, supporting the scenario where interstitial dying cells contribute to ROS signaling from via H_2_O_2_ production in physiological conditions (**Fig. 7G**). However, as previously shown^50^, apical regeneration is only transiently delayed after HU treatment (**Fig. 7H, 7I, Supplementary Fig. S17**). All together, these experiments indicate that injury-induced ROS signals contribute to induce cell death of interstitial cells, which themselves release H_2_O_2_ and thus contribute to the amplification of ROS signaling in neighboring epithelial cells (**Fig. 8A-8C, Table-S4**).

**Fig. 8.**
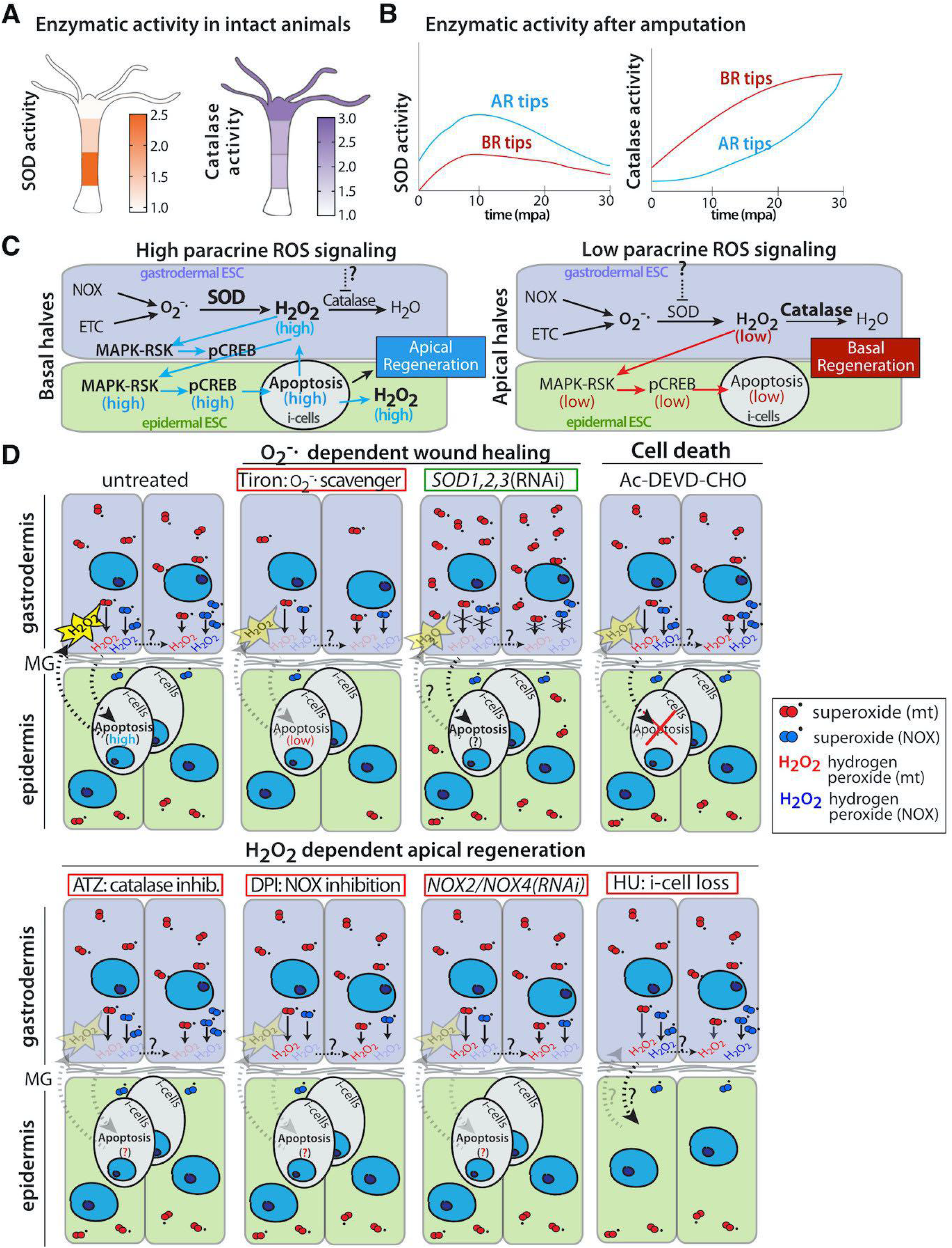
Summary scheme about two distinct ROS signaling in apical- and basal-regenerating tips. **(A)** Levels of SOD (left) and catalase (right) activities along the body axis of intact *Hydra*. **(B)** Modulations of SOD (left) and catalase (right) activities in apical- and basal-regenerating (AR, BR) tips during the first 30 minutes after bisection. **(C)** Model of injury-induced paracrine ROS signaling between gastrodermal and epidermal layers after mid-gastric bisection, high in AR tips and low in BR tips. In AR tips, high levels of SOD and low levels of catalase generate high levels of H_2_O_2_, which as paracrine signal, triggers MAPK/ERK/RSK activation followed by the death of interstitial-derived cells (grey) located in the epidermis. Dying cells release H_2_O_2_ that amplifies the process, a necessary condition for efficient apical regeneration. In BR tips, SOD levels are low and catalase levels rather high generating low levels of H_2_O_2_, explaining the observed low level of injury-induced cell death. **(D)** Pharmacological and genetic modulations of ROS signaling in apical-regenerating *Hydra*. In untreated *Hydra*, mitochondrial (red) and membrane (blue) superoxide are predominantly produced in gESCs (purple). Upon **Tiron** treatment, superoxide is scavenged, H_2_O_2_ is no longer produced, wound healing is inhibited, injury-induced cell death is abolished and AR is delayed. When ***SOD1, SOD2* or *SOD3*** are partially knocked-down, superoxide dismutation is reduced leading to lower levels of H_2_O_2_ production and superoxide accumulates. As a consequence, wound healing is speed-up, whereas AR and BR are not significantly modified. The **Ac-DEVD-CHO** treatment that inhibits caspase activity, blocks cell death and prevents H_2_O_2_ amplification. Upon **ATZ** treatment, catalase is inhibited, H_2_O_2_ levels surprisingly found severely reduced, leading to AR but not BR blockade. Upon **DPI** treatment, NOX enzymes are inhibited, mbO_2_^.-^ and H_2_O_2_ levels greatly reduced leading to AR blockade. When ***NOX4*** or ***NOX2/CYBB*** are knocked-down, NOX-dependent H_2_O_2_ production is reduced and AR is delayed. When interstitial cells are eliminated upon **HU** treatment, injury-induced cell death is reduced, H_2_O_2_ levels are not amplified and AR is delayed.

## DISCUSSION

Here we present a series of experimental evidence showing that two different types of ROS molecules are generated on either side of the bisecting plane in regenerating *Hydra*; one symmetrical, mitochondrial superoxide (mtO_2_^.-^), which promotes wound healing by acting primarily in a cell autonomous manner, and the other asymmetrical, hydrogen peroxide (H_2_O_2_), which acts in a paracrine mode to initiate apical regeneration, released first by gastrodermal epithelial cells to trigger death of surrounding interstitial cells, and then by dying cells of the interstitial lineage (**Fig. 8C, 8D**).

### Mitochondrial superoxide acts as a key signal for wound healing in Hydra

MtO_2_^.-^ is a short-lived molecule, active at the mitochondrial or cytosolic level but unable to cross cell membranes and to act as a paracrine signal^51^. Three types of evidence support the fact that in *Hydra* mtO_2_^.-^ directly orchestrates wound healing: (i) its similar immediate production after any type of wounding, followed or not by regeneration, (ii) the negative impact of O_2_^.-^ scavenging on wound healing while it is limited on regeneration, (iii) the positive impact of mtO_2_^.-^ accumulation on wound healing as observed when *SOD1, SOD2 or SOD3* are knocked-down. This wound healing function of mtO_2_^.-^ appears evolutionarily conserved as also reported in *C. elegans*^15^, *Drosophila*^52^ and mice^53^.

In *C. elegans*, a large ventral epidermal cell can be pierced and used as a wound healing model. In this context, RhoA, a member of the GTPase family that regulates cytoskeletal dynamics, is a direct target of mtROS^15^. The key mediator of cytoskeletal rearrangement is the Rho-associated kinase (ROCK), which once activated by Rho, promotes cytoskeletal rearrangement through its substrate myosin light chain (MLC). In *Drosophila* embryos, the dorsal closure provides a wound healing model where different levels of mtROS mediate different levels of ROCK activity and subsequently low and high tissue tension^52^. All components of the Rho/ROCK/MLC cascade are expressed in *Hydra*, but their activation by mtO2^.-^ during wound healing remains to be shown. In mammals, mtO_2_^.-^ is produced in response to hypoxia, acting positively on wound healing when transiently present but negatively when chronically present as observed in chronic wounds^53^. In *Hydra*, the hypoxia-inducible factor 1-alpha (<u>HIF1A</u>) is expressed at high levels along the body axis and in all stem cell populations, but its role is unknown.

In *Hydra*, mtO_2_^.-^ production appears to rely predominantly on the gastrodermal ESCs, where it can be detected at least for 60 minutes after injury. This observation suggests that digestive cells play the leading role in ROS-dependent wound healing while epidermal ESCs are activated either via a ROS-independent process and/or a ROS-dependent non-cell autonomous mechanism. Mitochondrial ROS in gESCs might activate signaling pathways via the redox-sensitive transcription factors such as NFE2F/Nrf, JunD or fra-l1^54^ and thus indirectly trigger non-cell autonomous effects. The initial role of gastrodermal cells in wound healing is in line with previous histological analyses showing that the gastrodermis heals first^55^.

### Hydrogen peroxide acts as a paracrine signal for injury-induced cell death and apical regeneration in Hydra

H_2_O_2_ has a longer half-life than O_2_^.-^ and can diffuse over relatively long distances in the extra-cellular space, thus forming gradients and acting in a paracrine fashion as demonstrated in zebrafish regeneration^12^. In *Hydra*, H_2_O_2_ production after bisection relies on immediate dismutation of O_2_^.-^ as scavenging O_2_^.-^ equally abolishes H_2_O_2_ production in AR and BR tips. Furthermore, the graded and inverted distribution along the axis of intact *Hydra* of SOD (basal to apical) and catalase (apical to basal) enzymatic activities explains the asymmetric levels of H_2_O_2_ on each side of the wound after bisection, high in the AR tips and low in the BR tips (**Fig. 8A, 8B**). These results support the mathematical model proposed in this study, with H_2_O_2_ corresponding to the U signal predicted by the model, capable of triggering asymmetric signaling and specific molecular and cellular events necessary to regenerate the head. This scenario is confirmed by the fact that CREB phosphorylation and injury-induced cell death in AR tips are suppressed when Tiron treatment scavenges O_2_^.-^ and thus prevents H_2_O_2_ production. Furthermore, cell death is observed over a distance of 100 µm from the amputation plane^7^, which is consistent with the expected diffusion of H_2_O_2_ as measured in amputated fins of zebrafish larvae^12^.

This work also identifies a secondary source of H_2_O_2_ production in *Hydra* halves regenerating their head, in line with the massive wave of interstitial cell death that takes place in apical-regenerating tips. Indeed, the observed increase in H_2_O_2_ levels in apical-regenerating animals no longer takes place in animals treated with Ac-DEVD-CHO, an inhibitor of effector caspases. This evidence is strengthened by the fact that H_2_O_2_ levels are also lowered after elimination of the interstitial cells with HU treatment, thus removing the potential source of dying cells (**Fig. 8D**). Both approaches indicate that interstitial cell death is central to the loop that ensures sustained high levels of H_2_O_2_ during the immediate phase of apical regeneration. A similar ROS – cell death loop was described in *Drosophila* imaginal discs, where depending on the ROS levels, two different cellular processes can be launched through the activation of Ask1 and Jun proteins: 1) cell proliferation/survival when ROS levels are low and 2) cell death when ROS levels are high. In the latter, high levels of ROS ensure high Jun activity, which in turns produces more ROS via a positive feedback loop^17,18^.

Beside its role on cell death, H_2_O_2_ also acts as a chemoattractant as in the zebrafish larvae where H_2_O_2_ released in the immediate post-injury phase recruits leucocytes that migrate towards the wound^12^. In the zebrafish larva regenerating its tail, H_2_O_2_ causes the reposition of notochord cells near the damage site, while these cells in turn secrete Hedgehog ligands required for regeneration^56^. In *Hydra*, interstitial progenitors do migrate towards the wound^57,7,58^, however this migration is seemingly not affected when animals are treated with Tiron (**Supplementary Fig. S18**). H_2_O_2_ also acts as a signaling molecule as in *Xenopus* tadpoles where H_2_O_2_ is a necessary signal to activate Wnt signaling^19^. In *Hydra* regenerating its head, this is also the case, at least indirectly, as injury-induced cell death triggers an immediate Wnt release visible around apoptotic bodies, followed by *Wnt3* up-regulation in gESCs^7^.

### ROS signaling via the MAPK/ERK pathway in regeneration

In multiple contexts, oxidative stress activates MAPK signaling by modulating ERK, JNK, p38, MEK activities^59^, which occurs through inhibition of protein phosphatase^24–26^. This study shows that in *Hydra* injury-induced H_2_O_2_ is an important player in injury-induced activation of the MAPK/ERK pathway, which in turns leads to phosphorylation of the CREB transcription factor, a molecular cascade necessary for apical but not basal regeneration^60^ (**Fig. 8C**). *Hydra* does express numerous components of the MAPK pathway including <u>ERK/MAPK1</u>, <u>JNK/MAPK10</u>, <u>p38/MAPK14</u>, <u>MEK/MAP2K</u>1, <u>MEK/MAP2K5</u> and <u>MAP3K15</u>, an Ask1-domain containing kinase. *Hydra* also expresses an *<u>AKT3</u>*-related gene in its epithelial cells. In the immediate phase after bisection, a large number of redox-sensitive proteins exhibit an impulse transient up-regulation^37,54^ suggesting that injury-induced ROS signaling rapidly triggers a transcriptional response. We identified five distinct transcription factors that exhibit such immediate transient up-regulation (ref-1, junD, ATF1/CREB, Fral-1, NFE2l/Nrf) and further studies will clarify their role in this immediate post-injury period when the apical organizer gets established. A recent study shows that treating *Hydra* with the antioxidant glutathione leads to a decrease in JNK, p38 and ERK phosphorylation in regenerating tips and alters apical regeneration as observed 72 hours after mid-gastric bisection^61^.

These results are consistent with studies showing that in bilaterians injury-induced ROS trigger regeneration by activating MAPK/ERK kinases, JNK activation in adult zebrafish regenerating its fin^21^, ERK1/2 phosphorylation in zebrafish regenerating its heart where injury-induced H_2_O_2_ destabilizes the redox-sensitive phosphatase Dusp6^62^. In mice, sustained elevated levels of H_2_O_2_ activate the ERK pathway, which promotes hepatocyte proliferation necessary for liver regeneration^63^. In planarians, H_2_O_2_ can rescue regeneration defects due to the MAPK-ERK silencing^64^. In *Drosophila* larvae regenerating their wings, Ask1 acts as a redox-sensitive protein that interacts with the insulin pathway kinase Akt in cells dedicated to regenerate, while inducing high JNK activity in cells committed to enter apoptosis^18^. These results obtained in distinct phyla suggest that the similar temporal chain of events that takes place after injury, injury-induced production of ROS, activation of the MAPK/ERK signaling pathway, localized cell death, release of growth factors by the dying cells and compensatory proliferation, likely reflects an ancestral sequence of events to regenerate and survive environmental threats.

## MATERIALS AND METHODS

### Mathematical modeling and fitting to previous experimental results

The mathematical model comprises three layers: the cells, the extracellular signaling and the intracellular signaling and is composed of the following one-dimensional partial differential equations and ordinary differential equations:

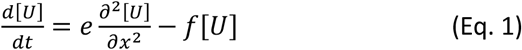

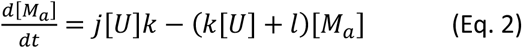

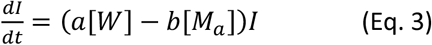

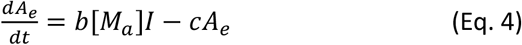

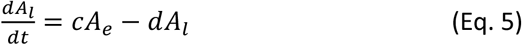

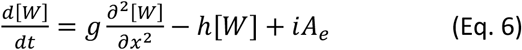

The model initial conditions are:

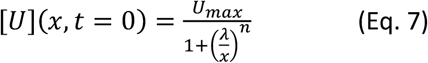

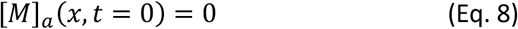

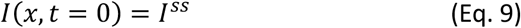

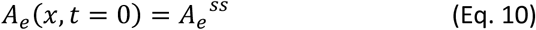

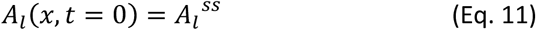

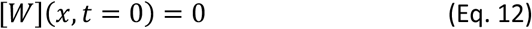

It is assumed that all cell densities start from the homogenous steady state I^ss^, A_e_^ss^ and A_l_^ss^ present in *Hydra* before amputation and zero-flux boundary conditions. In the model, mid-gastric bisection releases or produces the signal *U*, which diffuses and undergoes lytic degradation (**Eq. 1**). This signal activates the MAPK pathway (*M*_*i*_) by means of a bilinear reaction with the concentration of the substrate, the non-phosphorylated or inactive form of the enzymes (*M*_*i*_) and the signal (*U*) (**Eq. 2**). The backward reaction rate is assumed linear in the phosphorylated or activated signal (*M*_*a*_, **Eq. 2**).

The parameters involved in Equations 1-12 are the diffusion coefficient of *U* (*e*), the lytic degradation of *U* (*f*), the diffusion coefficient of *W* (*g*), the lytic degradation of *W* (*h*), the coupling between *W* production and density of early apoptotic cells (*i*), the kinetic rate constant of the transformation of *M*_*i*_ in *M*_*a*_ (*k*), the kinetic of the reverse reaction (*l*), the total concentration of *M* (*M*_*i*_ + *M*_*a*_ =*j*), the rate of proliferation of interstitial cells (*a*), the rate of apoptosis of these cells – this is also the rate of early apoptotic cells (*b*), the rate at which early apoptotic cells become late apoptotic cells (*c*), the rate at which late apoptotic cells are eliminated (*d*), the initial maximal concentration of *U* at the amputation plane (*U*_*max*_), the initial spatial inflection point of *U* (*λ*), and the Hill coefficient *n*. Simulations were performed in a C++ code in which the coupled ordinary and partial differential equations are numerically integrated on a one-dimensional regular grid using a Runge-Kutta time-stepping scheme of 4^th^ order and a finite difference approximation with six neighbors for spatial derivatives where the time and space discretization steps are 0.006 minutes and 0.005 mm, respectively.

### *Hydra* culture

*Hydra vulgaris (Hv)* from the *Basel (Hv_Basel), Hm-105* or *AEP2* (*Hv_AEP2*) strains were cultured at 18°C in Hydra medium (HM: 1 mM NaCl, 1 mM CaCl_2_, 0.1 mM KCl, 0.1 mM MgSO_4_, 1 mM Tris pH 7.6). The Endo-LifeAct-GFP and Ecto-LifeAct-GFP AEP transgenic strains^44,45^ are a kind gift from Bert Hobmayer (Innsbrück, Austria) and are named in this study LifeAct-gGFP and LifeAct-eGFP respectively. Animals were fed 3-4 times a week with freshly hatched *Artemia* nauplii and starved for one full day prior any experiment.

### Phylogenetic analyses

Phylogenetic reconstruction of the SOD and NOX families was performed as reported in ref^65^. Briefly, *Hydra* SOD- and NOX-related sequences identified on HydrAtlas were used to retrieve related sequences from species representative of metazoan and non-metazoan phyla (**Table-S1, Table-S2**). These sequences were subsequently aligned using the software Muscle Align (www.ebi.ac.uk/Tools/msa/muscle/) and the phylogenetic trees were built with PhyML3.0^66^ using the LG substitution model with 4 transition/transversion ratio and 6 to 8 substitution rate categories. The robustness of the branch support was tested with the aLRT SH-like fast likelihood method. The definitive trees was designed with the iTOL tool^67^.

### Pharmacological treatments

To prevent superoxide production, animals were exposed to the (NADPH) oxidase (NOX) inhibitor DPI (Diphenyleneiodonium chloride, BML-CN240, Enzo Life Sciences) 10 µM final prepared from a 10 mM stock solution diluted in DMSO. To scavenge superoxide, animals were treated with Tiron (4,5-Dihydroxy-1,3-benzenedisulfonic acid disodium salt monohydrate Sigma-Aldrich, N°172553) 15 mM final, prepared from a 0.5 M stock solution diluted in dH_2_O. To inhibit catalase, animals were exposed to 3-aminotriazole (ATZ, Sigma A8056-10G) 75 mM final prepared from a 0.5 M stock solution diluted in dH2O. To block apoptosis, animals are treated with Ac-DEVD-CHO (Enzo, ALX-260-030-M001) 10 µM final prepared from a 10 mM stock solution diluted in DMSO. To eliminate interstitial cells, *Hv_AEP2* animals were treated as in ref.^50^, subjected to three 24-hour long cycles of hydroxy-urea (HU, abcr AB117335) 5 mM final diluted in HM, each cycle separated by a 12 hour release period during which the animals were maintained in HM. During HU treatment, animals were fed normally but not washed until the end of the 24-hour period. Control animals were fed and washed in the same manner.

### Live imaging of mitochondrial superoxide (mtO_2_^.-^)

For MitoSOX labeling (M36008, Molecular Probes) a 5 µM solution was prepared in 1 ml HM immediately before use. Three *Hv_AEP2* animals were incubated for 60 minutes in MitoSOX labeling, then bisected at 50% level or lateraly ‘‘nicked’’ and mounted with the wound up-side in a 35 mm imaging dish (Cellvis, D35-20-1.5-N) for inverted microscopy in 1% low-melting agarose (Bio-Rad, 161-3113) containing 2% Urethane (Sigma, U2500). Bisection, mounting and solidification of agarose takes in total around 10 minutes, corresponding to the first imaging time-point after amputation. Imaging was done using the hybrid inverted spinning disk confocal microscope Marianas 3i. In case of drug treatment, animals were incubated with MitoSOX together with either DPI (10 µM) or Tiron (15 mM) for 60 minutes before bisection and processed as above. After treatment, animals were extensively washed in HM, mounted in distinct wells of the same 4-chamber imaging dish (Cellvis, D35C4-20-1.5-N) to allow simultaneous imaging. All images were analyzed with Imaris software using the ‘‘spot’’ tool for quantification. Minimal size of the detected spot was set at 5 µm with channel intensity set to minimum 70. For each condition, the number of dots was counted with the same macro at the indicated time-points and the fold change calculated relative to the value obtained in basal-regenerating tips of untreated animals taken 10 minutes after amputation.

### Hydrogen peroxide (H_2_O_2_) quantification in live *Hydra*

The Amplex UltraRed (Invitrogen, A36006) solution containing 100 µM Amplex UltraRed and 0.2 U/ml of Horseradish peroxidase (HRP) was freshly prepared prior to each assay. Ten *Hv_AEP2* animals were bissected and the upper and lower halves placed into separate tubes prefilled with HM. Ten halves were then transferred with 50 µl of HM into a black 96-well fluorescence microplate, with each well pre-filled with 50 µl of Amplex UltraRed solution. After gentle mixing, the microplate was placed in the Victor X5 plate reader (Perkin Elmer) for continuous fluorescence reading in five-minute increments using the red channel. The background value obtained by measuring a well containing 50 µl of Amplex UltraRed solution diluted in 50 µl HM, was subtracted from each result. To measure H_2_O_2_ when catalase is inhibited, 10 animals were incubated for 3 hours in 75 mM ATZ. To measure H_2_O_2_ when apoptosis is blocked, 30 animals were treated for 1 hour with 10 µM Ac-DEVD-CHO. After drug treatment, animals were washed intensively in HM, bisected and subjected to the H_2_O_2_ quantification procedure as previously described.

### SOD and Catalase activity assays

Briefly, 15 *Hv_AEP2* animals were bisected in four regions (apical, upper gastric, lower gastric and basal) and regenerating tips dissected at 10, 20 and 30 mpa. Whole cell extracts were then prepared in parallel, using 1:4 phosphate buffer from the Total SOD kit (Sigma-Aldrich, kit 19160) for SOD activity, the Catalase kit buffer (Abcam, kit ab83464) for catalase activity. After measuring the protein concentration via Bradford assay, 2 µg protein extracts were used for each condition according to the supplier guidelines.

### Peroxidase staining

After relaxing the animals in 2% urethane/HM for one minute, animals were fixed for 2 hours at RT in 4% PFA prepared in HM (pH = 7.5), then washed 3× 10 min in PBS, incubated in 500 μL DAB (SIGMAFAST™ 3,3’-Diamino-benzide) solution for 10 min, washed 3x 10 min in PBS. The DAB solution is freshly prepared with 1 tablet of DAB dissolved in 10 mL of PBS and filtered through a 0.22 μm filter. 5 mL of the filtered solution was added to 5 mL of PBS together with 20 μL Triton X-100 (0.2%) and 1 μL 30% H_2_O_2_ solution.

### Gene silencing via RNA interference

To knock-down genes of interest, *Hv_AEP2* animals were electroporated with small interfering RNA (siRNAs). For each targeted gene, three distinct siRNAs were designed (see **Table-S3**), produced by Qiagen, resuspended in ddH_2_O at 40 µM and stored at -20°C. 25 animals per condition were briefly washed in HM, incubated for 45 min in Milli-Q water, then placed in 200 µl 10 mM sterilized HEPES solution (pH 7.0) and transferred into a 0.4 cm gap electroporation cuvette (Cell Projects Ltd). Animals were electroporated with a scramble control siRNA or an equimolecular mixture of siRNAs (siRNA-a, siRNA-b, siRNA-c, 4 µM final) targeting the gene of interest using the Biorad GenePulser Xcell electroporation system at the following conditions: Voltage: 150 Volts; Pulse length: 50 milliseconds; Number of pulses: 2; Pulse intervals: 0.1 seconds. After electroporation, animals were transferred to a 10 cm diameter plastic dish containing HM, washed 16 hours later with fresh HM and dissociated tissues were discarded. Electroporation was repeated every other day, up to 5 times. All subsequent experiments were performed two days after the last electroporation.

### Q-PCR analysis

Total RNA was extracted using the E.Z.N.A.^®^ Total RNA kit (Omega) and cDNA synthesized using the qScript™ cDNA SuperMix (Quanta Bio-sciences). qPCR was performed in a 96-well format using the SYBR™ Select Master Mix for CFX (Thermo Fisher Scientific) and a Biorad CFX96™ Real-Time System. The *TATA-Box Binding protein (TBP)* gene was used as an internal reference gene; primer sequences can be found in **Table-S3**.

### Wound healing imaging

To visualize wound healing, regenerating halves were fixed in 4% PFA overnight, then washed 3x 10 minutes in PBS, labeled with Phalloidin-488 (Invitrogen, A12379) 1:40 for one hour at RT in the dark, washed 3x 10 minutes in PBS, counterstained in Hoechst for 30 minutes, washed 2x 5 minutes in PBS and 1x 5 minutes in dH2O, and finally mounted in 1% agarose with the wound up-side. Imaging was performed on the inverted spinning disk confocal Marianas 3i. Quantification of the wound size was performed with the Fiji software using the free-shape tool to calculate the surface of the full-bisected plane and that of the wound. The ratio of the wound surface over the total transversal surface of the animal was expressed as percentage.

### Regeneration experiments

Before bisection, animals were first kept in 10 cm diameter plastic dishes (Greiner, ref. 628102), with 1 ml HM per animal and then bisected. Regeneration experiments were carried out with triplicates, 10 to 30 animals per replicate and the emergence of tentacle rudiments was recorded over seven days to assess head formation. To modulate superoxide production or activity at injury time or during regeneration, animals were exposed either to the NOX inhibitor DPI 10 µM or to Tiron 15 mM for one hour prior to bisection and as indicated after bisection. For catalase inhibition, ATZ 75 mM was applied for three hours before bisection, kept after bisection and washed out after 24 hours.

### Detection of cell death on whole mounts

The In Situ Cell Death Detection Kit Fluorescein (Roche, N°11684-795-910) was used for detecting dying cells in intact *Hv_Basel*. Animals were fixed overnight in 50% ethanol, 3.7% formaldehyde, washed 3x 10 min in PBS, treated for 10 min in 0.2% sodium citrate, 0.5% Triton X-100 freshly prepared at RT, washed extensively 5x 3 min in PBS, incubated for 20 min at 70°C, washed 5 min in PBS. The kit components were mixed according to the guidelines, 50 µl is added to each 1.5 ml tube containing animals after careful removal of PBS. Samples were subsequently incubated in the dark for 90 min at 37°C, washed 3x 10 min in PBS, counterstained with 20 µM Draq5 (Biostatus) for 30 min, washed again in PBS, counter-stained with Hoechst 33342 (Thermofisher, 1 µg/ml), washed 2x 5 min in PBS, 5 min in dH2O and mounted in Mowiol. Imaging was done with the Leica SP5 confocal microscope, 40x objective.

### Cell death quantification on macerated tissues

25 *Hv_Basel* animals were bissected and the AR tips dissected at indicated regeneration time-points. Collected slices were put in 25 µl maceration solution (7% glycerol, 7% acetic acid) and macerated for 15 min, fixed in 4% PFA for 30 min, spread on electrically charged slides and dried for two days at RT. The Click-iT TUNEL Alexa Fluor488 Imaging Assay (C10245, Molecular Probes) kit was used for detecting apoptotic nuclei. The dried cells were washed 3x 10 min in PBS, treated for 2 min at RT in 0.5% sodium citrate, 0.5% Triton X-100, washed 5x 3 min in PBS, heated for 5 min at 70°C, washed 5 min in PBS, 5 min in dH_2_O. Next, we followed the guidelines of the supplier, then the slides were washed in 2% BSA, counterstained with Hoechst 33342 and MitoFluor598 (Molecular Probes, 200 nM) for 20 min, washed 2x 5 min in PBS, 5 min in dH_2_O and mounted in Mowiol. Cells from 4 or 5 distinct experiments per condition were counted with a minimum number of 2’000 cells.

### Immunodetection of phosphorylated CREB (pCREB) on *Hv_Basel* animals

The mouse monoclonal anti-pCREB antibody (Ser133, Millipore, # 05-807) was aliquoted upon arrival in the lab and stored at -20°C. Each aliquot was defrosted only once and kept at 4°C for less than 3 weeks. The Tyramide Signal Amplification kit (TSA, #40 Molecular Probes) with the HRP-conjugated anti-mouse goat antibody IgG Alexa Fluor 555 was used for detection. Animals taken at 30 min after bisection, were briefly relaxed in 2% urethane, fixed in 2% PFA diluted in HM supplemented with PhosSTOP (04906837001 Roche, one tablet/10 ml) for 15 hours at 4°C. Samples were then washed 3x 10 min in PBS, incubated in freshly prepared 2 N HCl for 30 min, washed 4x 5 min in PBS, blocked with freshly prepared 3% H_2_O_2_ for one hour at RT, washed 4x 5 min in PBS and finally blocked in 2% BSA for 1 hour at RT. Samples were subsequently incubated in anti-pCREB (1:400) diluted in 2% BSA overnight at 4°C, washed 3x 10 min in PBS and incubated for 4 hours at RT in 1:100 anti-mouse HRP in 2% BSA diluted in BSA. After 3x 10 min washes in PBS, detection was performed as indicated by the supplier with 1:100 Tyramide-555 for 18 min at RT. The reaction was stopped by first washing the samples 5 min in PBS then by incubating them in 3% H_2_O_2_ for 20 min. Samples were counter-stained with Hoechst 33342, washed 2x 5 min in PBS and 5 min in dH2O, mounted in Mowiol and imaged with a Leica SP5 confocal microscope.

## Supporting information

2022_Suknovic_SUPPLEMENTAL FILE

## CONTRIBUTIONS

NS performed majority of experiments presented in this manuscript, SR contributed with pCREB immunostaining and TUNEL staining experiments, OC performed mathematical modeling, DM contributed to the project via numerous consultations, NS prepared the figures, NS and BG wrote and revised the manuscript, BG obtained the funding and supervised the study.

## ACKNOWLEDGEMENTS

The authors thank Bert Hobmayer (Innsbrück) for providing the LifeAct-GFP transgenic strains, Joe Dan Dunn for his suggestion to use Amplex UltraRed to quantify H_2_O_2_ in vivo, Florenci Serras, Lutz Brusch and Andreas Deutsch for helpful discussions, Chrystelle Perruchoud for technical support and Denis Benoni for animal care. Nenad Suknovic was supported by an iGE3 doctoral fellowship. This work was supported by the Canton of Geneva, the Swiss National Science Foundation (SNF grants 31003A_149630, 31003_169930), the Human Frontier Science Program (HFSP grant no. RGP0016/2010) and the Claraz donation.

## REFERENCES

1. Bergmann, A. & Steller, H. Apoptosis, stem cells, and tissue regeneration. Sci Signal 3, re8 (2010).

2. Vriz, S., Reiter, S. & Galliot, B. Cell death: a program to regenerate. Curr. Top. Dev. Biol. 108, 121–51 (2014).

3. Perez-Garijo, A. & Steller, H. Spreading the word: non-autonomous effects of apoptosis during development, regeneration and disease. Development 142, 3253–62 (2015).

4. Bosch, T. C. Hydra and the evolution of stem cells. BioEssays 31, 478–86 (2009).

5. Buzgariu, W., Al Haddad, S., Tomczyk, S., Wenger, Y. & Galliot, B. Multi-functionality and plasticity characterize epithelial cells in Hydra. Tissue Barriers 3, e1068908 (2015).

6. Vogg, M. C., Buzgariu, W., Suknovic, N. S. & Galliot, B. Cellular, Metabolic, and Developmental Dimensions of Whole-Body Regeneration in Hydra. Cold Spring Harb. Perspect. Biol. a040725 (2021) doi:10.1101/cshperspect.a040725.

7. Chera, S. et al. Apoptotic Cells Provide an Unexpected Source of Wnt3 Signaling to Drive Hydra Head Regeneration. Dev. Cell 17, 279–289 (2009).

8. Galliot, B., Welschof, M., Schuckert, O., Hoffmeister, S. & Schaller, H. C. The cAMP response element binding protein is involved in hydra regeneration. Development 121, 1205–16 (1995).

9. Kaloulis, K., Chera, S., Hassel, M., Gauchat, D. & Galliot, B. Reactivation of developmental programs: The cAMP-response element-binding protein pathway is involved in hydra head regeneration. Proc. Natl. Acad. Sci. U. S. A. 101, 2363–8 (2004).

10. Chera, S., Ghila, L., Wenger, Y. & Galliot, B. Injury-induced activation of the MAPK/CREB pathway triggers apoptosis-induced compensatory proliferation in hydra head regeneration: MAPK-dependent injury-induced apoptosis in Hydra. Dev. Growth Differ. 53, 186–201 (2011).

11. Rojkind, M., Dominguez-Rosales, J. A., Nieto, N. & Greenwel, P. Role of hydrogen peroxide and oxidative stress in healing responses. Cell Mol Life Sci 59, 1872–91 (2002).

12. Niethammer, P., Grabher, C., Look, A. T. & Mitchison, T. J. A tissue-scale gradient of hydrogen peroxide mediates rapid wound detection in zebrafish. Nature 459, 996–9 (2009).

13. Moreira, S., Stramer, B., Evans, I., Wood, W. & Martin, P. Prioritization of competing damage and developmental signals by migrating macrophages in the Drosophila embryo. Curr. Biol. CB 20, 464–470 (2010).

14. Razzell, W., Evans, I. R., Martin, P. & Wood, W. Calcium flashes orchestrate the wound inflammatory response through DUOX activation and hydrogen peroxide release. Curr Biol 23, 424–9 (2013).

15. Xu, S. & Chisholm, A. D. C. elegans Epidermal Wounding Induces a Mitochondrial ROS Burst that Promotes Wound Repair. Dev Cell 31, 48–60 (2014).

16. Buchon, N., Broderick, N. A., Chakrabarti, S. & Lemaitre, B. Invasive and indigenous microbiota impact intestinal stem cell activity through multiple pathways in Drosophila. Genes Dev. 23, 2333–2344 (2009).

17. Santabárbara-Ruiz, P. et al. ROS-Induced JNK and p38 Signaling Is required for Unpaired cytokine activation during Drosophila regeneration. PLoS Genet. 11, e1005595 (2015).

18. Santabárbara-Ruiz, P. et al. Ask1 and Akt act synergistically to promote ROS-dependent regeneration in Drosophila. PLoS Genet. 15, e1007926 (2019).

19. Love, N. R. et al. Amputation-induced reactive oxygen species are required for successful Xenopus tadpole tail regeneration. Nat Cell Biol 15, 222–8 (2013).

20. Carbonell M B., Zapata Cardona, J. & Delgado, J. P. Hydrogen peroxide is necessary during tail regeneration in juvenile axolotl. Dev. Dyn. 251, 1054–1076 (2022).

21. Gauron, C. et al. Sustained production of ROS triggers compensatory proliferation and is required for regeneration to proceed. Sci Rep 3, 2084 (2013).

22. Saitoh, M. et al. Mammalian thioredoxin is a direct inhibitor of apoptosis signal-regulating kinase (ASK) 1. EMBO J. 17, 2596–2606 (1998).

23. Furuhata, M., Takada, E., Noguchi, T., Ichijo, H. & Mizuguchi, J. Apoptosis signal-regulating kinase (ASK)-1 mediates apoptosis through activation of JNK1 following engagement of membrane immunoglobulin. Exp. Cell Res. 315, 3467–3476 (2009).

24. Morita, K. -i. Negative feedback regulation of ASK1 by protein phosphatase 5 (PP5) in response to oxidative stress. EMBO J. 20, 6028–6036 (2001).

25. Kamata, H. et al. Reactive oxygen species promote TNFalpha-induced death and sustained JNK activation by inhibiting MAP kinase phosphatases. Cell 120, 649–61 (2005).

26. Chen, L., Liu, L., Yin, J., Luo, Y. & Huang, S. Hydrogen peroxide-induced neuronal apoptosis is associated with inhibition of protein phosphatase 2A and 5, leading to activation of MAPK pathway. Int. J. Biochem. Cell Biol. 41, 1284–1295 (2009).

27. Inupakutika, M. A., Sengupta, S., Devireddy, A. R., Azad, R. K. & Mittler, R. The evolution of reactive oxygen species metabolism. J. Exp. Bot. 67, 5933–5943 (2016).

28. Murphy, M. P. How mitochondria produce reactive oxygen species. Biochem. J. 417, 1–13 (2009).

29. Bedard, K. & Krause, K.-H. The NOX Family of ROS-Generating NADPH Oxidases: Physiology and Pathophysiology. Physiol. Rev. 87, 245–313 (2007).

30. Jiang, F., Zhang, Y. & Dusting, G. J. NADPH oxidase-mediated redox signaling: roles in cellular stress response, stress tolerance, and tissue repair. Pharmacol Rev 63, 218–42 (2011).

31. Landis, G. N. & Tower, J. Superoxide dismutase evolution and life span regulation. Mech. Ageing Dev. 126, 365–379 (2005).

32. Nguyen, N. H., Tran, G.-B. & Nguyen, C. T. Anti-oxidative effects of superoxide dismutase 3 on inflammatory diseases. J. Mol. Med. 98, 59–69 (2020).

33. Casareno, R. L. B., Waggoner, D. & Gitlin, J. D. The Copper Chaperone CCS Directly Interacts with Copper/Zinc Superoxide Dismutase. J. Biol. Chem. 273, 23625–23628 (1998).

34. Ambasta, R. K. et al. Direct Interaction of the Novel Nox Proteins with p22phox Is Required for the Formation of a Functionally Active NADPH Oxidase. J. Biol. Chem. 279, 45935–45941 (2004).

35. Altenhöfer, S., Radermacher, K. A., Kleikers, P. W. M., Wingler, K. & Schmidt, H. H. H. W. Evolution of NADPH Oxidase Inhibitors: Selectivity and Mechanisms for Target Engagement. Antioxid. Redox Signal. 23, 406–427 (2015).

36. Zana, M. et al. Interaction between p22phox and Nox4 in the endoplasmic reticulum suggests a unique mechanism of NADPH oxidase complex formation. Free Radic. Biol. Med. 116, 41–49 (2018).

37. Wenger, Y., Buzgariu, W., Perruchoud, C., Loichot, G. & Galliot, B. Generic and context-dependent gene modulations during <em>Hydra</em> whole body regeneration. bioRxiv 587147 (2019) doi:10.1101/587147.

38. Mukhopadhyay, P., Rajesh, M., Yoshihiro, K., Haskó, G. & Pacher, P. Simple quantitative detection of mitochondrial superoxide production in live cells. Biochem. Biophys. Res. Commun. 358, 203–208 (2007).

39. Mohanty, J. G., Jaffe, J. S., Schulman, E. S. & Raible, D. G. A highly sensitive fluorescent micro-assay of H2O2 release from activated human leukocytes using a dihydroxyphenoxazine derivative. J. Immunol. Methods 202, 133–141 (1997).

40. Fu, X. et al. cAMP-response Element-binding Protein Mediates Acid-induced NADPH Oxidase NOX5-S Expression in Barrett Esophageal Adenocarcinoma Cells. J. Biol. Chem. 281, 20368–20382 (2006).

41. Wang, X. et al. Early controlled release of peroxisome proliferator-activated receptor β/δ agonist GW501516 improves diabetic wound healing through redox modulation of wound microenvironment. J. Controlled Release 197, 138–147 (2015).

42. Venkatachalam, K. et al. Motor Deficit in a Drosophila Model of Mucolipidosis Type IV due to Defective Clearance of Apoptotic Cells. Cell 135, 838–851 (2008).

43. Xiong, Y., Contento, A. L., Nguyen, P. Q. & Bassham, D. C. Degradation of Oxidized Proteins by Autophagy during Oxidative Stress in Arabidopsis. Plant Physiol. 143, 291–299 (2007).

44. Aufschnaiter, R., Wedlich-Soldner, R., Zhang, X. & Hobmayer, B. Apical and basal epitheliomuscular F-actin dynamics during Hydra bud evagination. Biol. Open 6, 1137–1148 (2017).

45. Livshits, A., Shani-Zerbib, L., Maroudas-Sacks, Y., Braun, E. & Keren, K. Structural Inheritance of the Actin Cytoskeletal Organization Determines the Body Axis in Regenerating Hydra. Cell Rep 18, 1410–1421 (2017).

46. Ledenev, A. N., Konstantinov, A. A., Popova, E. & Ruuge, E. K. A simple assay of the superoxide generation rate with Tiron as an EPR-visible radical scavenger. Biochem. Int. 13, 391–396 (1986).

47. Yamada, J. et al. Cell permeable ROS scavengers, Tiron and Tempol, rescue PC12 cell death caused by pyrogallol or hypoxia/reoxygenation. Neurosci. Res. 45, 1–8 (2003).

48. Taiwo, F. A. Mechanism of tiron as scavenger of superoxide ions and free electrons. Spectroscopy 22, 491– 498 (2008).

49. Nicholls, P. The reaction between aminotriazole and catalase. Biochim. Biophys. Acta 59, 414–420 (1962).

50. Buzgariu, W., Wenger, Y., Tcaciuc, N., Catunda-Lemos, A. P. & Galliot, B. Impact of cycling cells and cell cycle regulation on Hydra regeneration. Dev. Biol. 433, 240–253 (2018).

51. Kowaltowski, A. J., de Souza-Pinto, N. C., Castilho, R. F. & Vercesi, A. E. Mitochondria and reactive oxygen species. Free Radic. Biol. Med. 47, 333–343 (2009).

52. Muliyil, S. & Narasimha, M. Mitochondrial ROS Regulates Cytoskeletal and Mitochondrial Remodeling to Tune Cell and Tissue Dynamics in a Model for Wound Healing. Dev. Cell 28, 239–252 (2014).

53. Fuhrmann, D. C. & Brüne, B. Mitochondrial composition and function under the control of hypoxia. Redox Biol. 12, 208–215 (2017).

54. Wenger, Y., Buzgariu, W., Reiter, S. & Galliot, B. Injury-induced immune responses in Hydra. Semin. Immunol. 26, 277–94 (2014).

55. Bibb, C. & Campbell, R. D. Tissue healing and septate desmosome formation in hydra. Tissue Cell 5, 23–35 (1973).

56. Romero, M. M. G., McCathie, G., Jankun, P. & Roehl, H. H. Damage-induced reactive oxygen species enable zebrafish tail regeneration by repositioning of Hedgehog expressing cells. Nat. Commun. 9, 4010 (2018).

57. Fujisawa, T., David, C. N. & Bosch, T. C. Transplantation stimulates interstitial cell migration in hydra. Dev. Biol. 138, 509–12 (1990).

58. Boehm, A. M. & Bosch, T. C. Migration of multipotent interstitial stem cells in Hydra. Zool. Jena 115, 275–82 (2012).

59. McCubrey, J. A., LaHair, M. M. & Franklin, R. A. Reactive Oxygen Species-induced activation of the MAP Kinase signaling pathways. Antioxid. Redox Signal. 8, 1775–1789 (2006).

60. Galliot, B. Injury-induced asymmetric cell death as a driving force for head regeneration in Hydra. Dev. Genes Evol. 223, 39–52 (2013).

61. Tursch, A. et al. Injury-induced MAPK activation triggers body axis formation in Hydra by default Wnt signaling. Proc. Natl. Acad. Sci. 119, e2204122119 (2022).

62. Han, P. et al. Hydrogen peroxide primes heart regeneration with a derepression mechanism. Cell Res. 24, 1091–1107 (2014).

63. Bai, H. et al. Hydrogen peroxide modulates the proliferation/quiescence switch in the liver during embryonic development and posthepatectomy regeneration. Antioxid. Redox Signal. 22, 921–937 (2015).

64. Jaenen, V. et al. Reactive oxygen species rescue regeneration after silencing the MAPK–ERK signaling pathway in Schmidtea mediterranea. Sci. Rep. 11, 881 (2021).

65. Suknovic, N., Tomczyk, S., Colevret, D., Perruchoud, C. & Galliot, B. The ULK1 kinase, a necessary component of the pro-regenerative and anti-aging machinery in Hydra. Mech. Ageing Dev. 194, 111414 (2021).

66. Guindon, S. et al. New algorithms and methods to estimate maximum-likelihood phylogenies: assessing the performance of PhyML 3.0. Syst Biol 59, 307–21 (2010).

67. Letunic, I. & Bork, P. Interactive Tree Of Life (iTOL) v5: an online tool for phylogenetic tree display and annotation. Nucleic Acids Res. 49, W293–W296 (2021).

