## Supplementary material for "Apical-to-basal graded ROS metabolism in intact *Hydra* leads to distinct levels of injury-induced ROS signaling in apical and basal regenerating tips": 2022_Suknovic_SUPPLEMENTAL FILE

### *Supplementary information*

### Supplementary Methods- Modeling

#### Parameter optimization

Algebraically, the problem of fitting consists in the exploration of the free parameter space in order to minimize a given function (*f*), in this case, the sum of the squares of the residues between the experimental data corresponding to the density of given cell types and those simulated by the model:

$$f = \sum_{i=1}^{N} \sum_{j=1}^{2} \sum_{k=1}^{3} \frac{\left(D_{j,k}^{exp}(t_i) - D_{j,k}^{theor}(t_i)\right)^2}{\left(\Delta D_{j,k}^{exp}(t_i)\right)^2}$$

Where $D_{j,k}^{exp}(t_i)$, □$D_{j,k}^{exp}(t_i)$ and $D_{j,k}^{theor}(t_i)$ are the experimental density of cell of type *k* in the space region *j* at the time $t_i$ , the standard deviations, and the corresponding values calculated from the model in the same conditions by averaging over the simulated spatial profile for the given region:

$$D_{j,k}^{theor} = \frac{1}{L_j} \sum_{l=1}^{L_j} \rho(x_l)$$

Where $\rho(x_l)$ is the density of interstitial, early or advanced apoptotic cells in the discretized cell $x_l$. $D_{j,k}$ =1, 2 and 3 correspond to interstitial (*I*), early apoptotic ($A_e$) and late apoptotic cell densities ($A_l$), respectively, in the space region *j*. Region 1 and 2 were previously defined as 0 ≤ x ≤ 100 µm and 100 µm ≤ x ≤ 300 µm from the amputation plane, respectively.

#### Typical values of the cellular densities

The model variables are the concentrations [*U*], $[M]_i$, $[M]_a$, [*W*] as well as the density of interstitial cells (*I*), early apoptotic cells ($A_e$) and late apoptotic cells ($A_l$). We performed a non-dimensionalization of the model using the typical values $I^*$, $A_e^*$, $A_l^*$, $[M]_i^*$, $[M]_a^*$ and $[W]^*$ estimated from literature.

Based on the percentages of interstitial and apoptotic cells observed in intact and amputated *Hydra* (ref[1]), $I^*$ and $A_e^* = A_l^*$ are estimated as 3.5 $10^4$ and 4.4$10^4$ cell.cm$^{-1}$, for a *Hydra* having a length of 1 cm and containing a total of 75,000 cells. We assume that the total concentration of MAPK was $[M]^*$, regardless the phosphorylation state of the enzyme: i.e., $[M]_i^* + [M]_a^* = [M]^*$. Since it was reported that a cell could contain 360,000 molecules of MAPK (ref[2]) and that *Hydra* contains between 10.8 % and 46.8 % of interstitial cells [1], in which MAPK are mainly expressed, we estimated that $[M]^*$ ranges between 4.8 $10^{-15}$ -6.2 $10^{-14}$ mol.cm$^{-1}$. The Wnt3 concentration range was reported within $10^{-15}$ – $10^{-10}$ mol.cm$^{-3}$ (ref[3]). Hence, assuming that the *Hydra* transversal width is 200 µm, $[W]^*$ was estimated as $10^{-19}$ - $10^{-14}$ mol.cm$^{-1}$.

### Supplemental Tables

| Species code | Species name | Cu-Zn SOD1 (SODC) | Cu-Zn SOD3 (SODS) | Mn SOD2 (SODM) | Copper Chaperone SOD (CCS) |
|---|---|---|---|---|---|
| **Acrmi** | *Acropora millepora* (Anthozoa, coral) | XP_029201721.1 | | XP_044179376.1 | XP_029179807.1 |
| **Actte** | *Actinia tenebrosa* (Anthozoa, anemone) | XP_031571030.1 | | XP_031555730.1 | A0A6P8IFH6_ACTTE |
| **Aedal** | *Aedes albopictus* (Arthropoda, tiger mosquito) | XP_029715437.1 | XP_019537847.1 | XP_019544173.1 | KXJ73503.1 |
| **Ampqu** | *Amphimedon queenslandica* (Demosponge) | XP_003388880.1 | | XP_003389045.1 | XP_003386470.2 |
| **Anevi** | *Anemonia viridis* (Anthozoa, anemone) | AAN85727.2 | AAN85728.2 | nd | nd |
| **Aplca** | *Aplysia californica* (Mollusca, sea slug) | Q8MUT8_APLCA | XP_005097192.1 | XP_005104703.1 | XP_005109651.1 |
| **Arath** | *Arabidopsis thaliana* (Viridipantae, | O78310 | | Q9LYK8 | Q9LD47 |
| **Brafl** | *Branchiostoma floridae* (Cephalochordata, amphioxus) | C3ZM91_BRAFL<br>C3ZAI5_BRAFL | C3YDS5_BRAFL<br>C3YVA5_BRAFL | XP_035687345.1<br>C3Z5Z4_BRAFL | XP_035688703.1 |
| **Caeel** | *Caenorhabditis elegans* (Nematoda, round worm) | NP_001021956.1 | P34461.2<br>(NP_001255003.1) | NP_492290.1 | nd |
| **Capow** | *Capsaspora owczarzaki* (Filasterea) | KJE97912.1 | | XP_004365015.1 | XP_004342913.2 |
| **Capte** | *Capitella teleta* (Annelida) | ELU08058.1 | ELT99613.1 | ELU11436.1 | ELU12420.1 |
| **Chlre** | *Chlamydomonas reinhardtii* (Viridiplantae, chlorophyta) | XP_001699077.1 | | Q42684_CHLRE | nd |
| **Clapu** | *Claviceps purpurea* (Fungi, ergot fungus) | Q96VL0_CLAP2 | | KAG6238627.1 | CCE27547.1 |
| **Cragi** | *Crassostrea gigas* (Mollusca, oyster) | UNY46907.1 | K1RVZ4_CRAGI | NP_001295847.1 | XP_011446778.1 |
| **Cripl** | *Cristaria plicata* (Mollusca, freshwater mussel) | ACI28282.1 | | QED22039.1 | nd |
| **Cyaca** | *Cyanea capillata* (Scyphozoa, giant jellyfish) | ALQ81852.1 | | nd | nd |
| **Danre** | *Danio rerio* (Actinopterygii, zebrafish) | NP_571369.1 | NP_001092706.1 | Q6P980_DANRE | F1QCS3_DANRE |
| **Dicdi** | *Dictyostelium discoideum* (Amoebozoa) | XP_647129.1<br>XP_635813.1 | | XP_645815.1 | XP_635464.1 |
| **Escco** | *Escherichia coli* (Bacteria) | RAX32182.1 | | MHO04530.1 | nd |
| **Exadi** | *Exaiptasia diaphana* (Anthozoa, sea anemone) | XP_020895630.1<br>XP_020895631.1 | | XP_020892835.1 | XP_020892029.1 |
| **Haldu** | *Halisarca dujardinii* (Demosponge) | QEH04774.1 | | QSX72305.1 | nd |
| **Halpa** | *Halichondria panicea* (Demosponge) | QSX72243.1 | | QSX72244.1 | nd |
| **Hensa** | *Henneguya salminicola* (Cnidaria, myxozoa) | KAF0986005.1 | | KAF0987259.1 | nd |
| **Human** | *Homo sapiens* (Mammalia) | P00441 or<br>NP_000445.1 | P08294 | NP_000627.2 | Sp O14618 |
| **Hydvu** | *Hydra vulgaris* (Hydrozoa, freshwater polyp) sequences from Uniprot (NCBI, HydrAtlas) | I3V7W8_HYDVU<br>(NP_001274724.1<br>seq37733_loc14321<br>c19962_g1_i01) | A0S5T9_HYDVU<br>(NP_001296642.1<br>seq12246_loc05612<br>c12652_g1_i01) | T2MGY4_HYDVU<br>(NP_001296665.1<br>seq69497_loc23238<br>c20941_g1_i02) | T2MJ72_HYDVU<br>(XP_002169753.3<br>seq71352_loc23686<br>c20965_g1_i02) |
| **Kleni** | *Klebsormidium nitens* (Klebsormidiales, green alga) | GAQ90353.1 | | GAQ87983.1 | GAQ90354.1 |
| **Lamsa** | *Lamellibrachia satsuma* (Annelida) | KAI0225630.1 | | KAI0243194.1 | KAI0236612.1 |
| **Mescr** | *Mesembryanthemum crystallinum* (Viridiplantae, ice plant) | P93258_MESCR | | nd | nd |
| **Myxsq** | *Myxobolus squamalis* (Cnidaria, myxozoa) | KAF1743046.1 | | KAF1742605.1 | nd |
| **Nauca** | *Naumovozyma castellii* (Fungi, budding yeast) | XP_003677709.1 | | XP_003677335.1 | nd |
| **Nemve** | *Nematostella vectensis* (Anthozoa, sea anemone) | EDO42041.1<br>EDO42040.1 | | XP_001641079.1 | XP_048590406.1 |
| **Orbfa** | *Orbicella faveolata* (Anthozoa, star coral) | XP_020600523.1 | XP_020600523.1 | XP_020624989.1 | XP_020604328.1 |
| **Pecma** | *Pecten maximus* (Mollusca, king scallop) | XP_033740917.1 | | XP_033740122.1 | XP_033734618.1 |
| **Phama** | *Phallusia mammillata* (Urochordata, | CAB3266471.1 | | CAB3266472.1 | CAB3228946.1 |
| **Phypa** | *Physcomitrium patens* (Viridiplantae, earthmoss) | XP_024383742.1 | | XP_024403365.1<br>XP_024399093.1 | XP_024401734.1 |
| **Pocda** | *Pocillopora damicornis* (Anthozoa, cauliflower coral) | RMX58501.1 | | RMX45412.1 | XP_027050048.1 |
| **Sacce** | *Saccharomyces cerevisiae* (Fungi, brewer's yeast) | NP_012638.1 | | P00447_SACCE | PTN40728.1 |
| **Sacko** | *Saccoglossus kowalevskii* (Hemichordata, acorn worm) | XP_002737020.1 | XP_006822462.1 | XP_002741802.1 | XP_006818904.1 |
| **Strpu** | *Strongylocentrotus purpuratus* (Echinodermata, sea urchin) | A0A7M7RB15_STRPU | A0A7M7N089_STRPU | A0A7M7RGD5_STRPU<br>A0A7M7RGR2_STRPU | XP_011674044.2 |
| **Stycl** | *Styela clava* (Urochordata, sea squirt) | XP_039262007.1 | | XP_039262481.1 | XP_039273736.1 |
| **Stypi** | *Stylophora pistillata* (Anthozoa, stony coral) | XP_022790361.1 | | XP_022799031.1 | A0A2B4SUW6_STYPI |
| **Tetth** | *Tetrahymena thermophila* (Ciliophora) | XP_001032187.1 | | XP_001010506.1 | nd |
| **Triad** | *Trichoplax adherens* (Placozoa) | XP_002116945.1 | | XP_002110053.1 | XP_002114197.1 |
| **Trica** | *Tribolium castaneum* (Arthropoda, red flour beetle) | XP_015835141.1 | KYB25410.1 | XP_972440.1 | XP_975577.1 |
| **Trisp** | *Trichinella spiralis* (Nematoda, round worm) | XP_003382001.1 | | KRY27591.1 | nd |
| **Verba** | *Verrucomicrobia bacterium* (Bacteria) | MBI4664285.1 | PYJ47568.1<br>PYJ46137.1 | MBS0615710.1 | nd |
| **Xentr** | *Xenopus tropicalis* (Amphibia, Western clawed frog) | NP_001016252.1<br>NP_001123681.1 | NP_001106630.1 | Q6DJC5_XENTR | Q63ZZ7_XENTR |
| **Zeama** | *Zea mays* (Planta, angiosperms, corn) | XP_008651855.1 | | P09233.1 | NP_001150157.1 |

#### Table-S1: Species code and accession numbers of the SOD sequences

Abbreviations: nd: not detected after Blastp on NCBI; SOD: Super Oxide Dismutase; SODC : cytoplasmic SOD; SODM: mitochondrial SOD; SODS: secreted SOD. Accession numbers written black correspond to the sequences used to produce the phylogenetic trees shown in **Figure S3** ; accession numbers written grey correspond to sequences identified as orthologous after Blastp on NCBI with e-values lower than $10^{-12}$.

| Species code | Species name | Phylum | NOX-like | NOXA-like | CyBB/NOX2 | NOX4 | NOX5 | DUOX |
|---|---|---|---|---|---|---|---|---|
| **Acama** | *Acaryochloris marina* | BACTERIA | | | | | YP_001517464.1 | |
| **Acrmi** | *Acropora millepora* | CNIDARIA (Anthoz) | | | XP_044181176.1<br>XP_029189986.2 | XP_029207032.2 | | |
| **Actte** | *Actinia tenebrosa* | CNIDARIA (Anthoz) | | XP_031569869.1 | XP_031549425.1 | XP_031559628.1 | | |
| **Ampqu** | *Amphimedon queenslandica* | PORIFERA | XP_019850557.1<br>(orphan) | | XP_003386282.1 | | XP_019850316.1<br>XP_011406986.2<br>XP_019858698.1<br>XP_019856436.1 | XP_003384066.1 |
| **Arath** | *Arabidopsis thaliana* | ANGIOSPERMAE | NP_001190833.1 | | | | | |
| **Batde** | *Batrachochytrium dendro.* | FUNGI | | | EGF78576.1 | | | |
| **Blyhe** | *Blyttiomyces helicus* | FUNGI | | RKO91938.1 | | | | |
| **Brafl** | *Branchiostoma floridae* | CEPHALO-CHORDATA | | | XP_002612231.1 | XP_035686017.1 | XP_002611039.1 | XP_002595019.1<br>XP_035660518.1<br>XP_035668371.1 |
| **Caebr** | *Caenorhabditis briggsae* | NEMATODA | | | | | | XP_045092860.1 |
| **Capow** | *Capsaspora owczarzaki* | FILASTEREA | | | XP_004345248.1 (EFW41367.1) | | | |
| **Capte** | *Capitella teleta* | ANNELIDATA | | | | | ELU05067.1<br>ELU05066.1 | ELU06497.1<br>ELU01203.1 |
| **Chlre** | *Chlamydomonas reinhar.* | CHLOROPHYTA | | XP_001691856.1 | | | | |
| **Cioin** | *Ciona intestinalis* | UROCHORDATA | | | NP_001121595.1 | NP_001029001.1 | | XP_018670917.1<br>XP_026693992.1 |
| **Cragi** | *Crassostrea gigas* | MOLLUSCA | | | XP_034331878.1 | | XP_019919960.2<br>XP_034322015.1<br>XP_034322027.1 | XP_034317672.1 |
| **Cyaca** | *Cyanea capillata* | CNIDARIA (Scypho) | | | | | | |
| **Danre** | *Danio rerio* | VERTEBRATA | | | NP_956708.1<br>AAI60633.1 | XP_005173476.4 | XP_001921894.1 | BAF33370.1 |
| **Dappu** | *Daphnia pulex* | ARTHROPODA | | | | XP_046448810.1 | XP_046447198.1<br>XP_046447189.1 | XP_046437872.1 |
| **Dengi** | *Dendronephthya gigantea* | CNIDARIA (Anthoz) | | | XP_028415268.1<br>XP_028415287.1 | | XP_028413422.1<br>XP_028405374.1<br>XP_028401239.1<br>XP_028394802.1 | XP_028409178.1 |
| **Dicdi** | *Dictyostelium discoideum* | AMOEBOZOAN | XP_635387.1<br>(orphan) | | XP_636064.1<br>XP_637386.1 | | | |
| **Exadi** | *Exaiptasia diaphana* | CNIDARIA (Anthoz) | | XP_020894978.1<br>KXJ28514.1 | XP_020896149.1<br>XP_020910621.1 | XP_028513814.1 | | |
| **Horvu** | *Hordeum vulgare* | ANGIOSPERMAE | BAK03598.1 | | | | | |
| **Human** | *Homo sapiens* | VERTEBRATA | | | NP_000388.2<br>NP_008983.2<br>NP_056533.1 | Q9NPH5.2 | Q96PH1 | Q9NRD9<br>NP_054799.4 |
| **Hydvu** | *Hydra vulgaris* | CNIDARIA (Hydroz) | | XP_002169684.3<br>(seq42412) | XP_012556948.1<br>(seq48686) | XP_012560964.1<br>(seq32691) | | |
| **Kleni** | *Klebsormidium nitens* | STREPTOPHYTA | | GAQ85633.1 | | | | |
| **Lamsa** | *Lamellibrachia satsuma* | ANNELIDA (polychaete) | | | KAI0229492.1 | KAI0209124.1 | KAI0231383.1<br>KAI0213699.1<br>KAI0213701.1<br>KAI0224797.1<br>KAI0210947.1 | KAI0221520.1<br>KAI0232396.1<br>KAI0231914.1<br>KAI0231915.1 |
| **Nemve** | *Nematostella vectensis* | CNIDARIA (Anthoz) | | XP_032234487.1 | XP_032235205.1<br>XP_001630388.1 | XP_032219431.1 | | |
| **Orbfa** | *Orbicella faveolate* | CNIDARIA (Anthoz) | | XP_020631551.1 | XP_020612067.1 | | | |
| **Pecma** | *Pecten maximus* | MOLLUSCA | | | XP_033741148.1 | XP_033739342.1 | XP_033725578.1 | XP_033724658.1<br>XP_033749554.1 |
| **Phama** | *Phallusia mammillata* | UROCHORDATA | | | CAB3235208.1 | | | |
| **Phyin** | *Phytophthora infestans* | FUNGI | | XP_002905918.1 | | | | |
| **Pocda** | *Pocillopora damicornis* | CNIDARIA (Anthoz) | | | XP_027047510.1<br>XP_027043844.1 | XP_027051542.1 | | |
| **Ricco** | *Ricinus communis* | ANGIOSPERMAE | XP_002516222.1 | | | | | |
| **Reine** | *Reinekea sp.* | BACTERIA | | | | | ZP_01113405.1 | |
| **Sacko** | *Saccoglossus kowalevskii* | HEMICHORDATA | | | XP_002738796.2 | XP_006823190.1 | XP_002731084.1<br>XP_002740169.1 | XP_002735792.1<br>XP_006813412.1 |
| **Scham** | *Schistocerca americana* | ARTHROPODA | | | | XP_046979754.1 | XP_047000632.1 | XP_046994195.1 |
| **Sorbi** | *Sorghum bicolor* | ANGIOSPERMAE | XP_002450884.1 | | | | | |
| **Strpu** | *Strongylocentrotus purpuratus* | ECHINODERMATA | | | XP_030850793.1 | XP_030851094.1 | XP_030829321.1 | XP_030832128.1<br>XP_030833017.1 |
| **Stycl** | *Styela clava* | UROCHORDATA | | | XP_039270448.1<br>XP_039256385.1 | XP_039251724.1 | | XP_039267420.1 |
| **Stypi** | *Stylophora pistillata* | CNIDARIA (Anthoz) | | | XP_022804580.1<br>XP_022794957.1 | XP_022795525.1 | | |
| **Tetth** | *Tetrahymena thermophila* | CILIOPHORA | | XP_001015231.1 | | | | |
| **Triad** | *Trichoplax adherens* | PLACOZOA | | | XP_002112911.1 | | XP_002115436.1 | |
| **Trica** | *Tribolium castaneum* | ARTHROPODA | | | | | XP_972375.2 | XP_970848.2<br>XP_008200778.1 |
| **Trisp** | *Trichinella spiralis* | NEMATODA | | | | | | XP_003376791.1 |
| **Vitvi** | *Vitis vinifera* | ANGIOSPERMAE | XP_002282296.1<br>XP_002277540.1 | | | | | |
| **Xenia** | *Xenia* | CNIDARIA (Anthoz) | | | XP_046849036.1 | | XP_046849727.1<br>XP_046863973.1<br>XP_046850046.1<br>XP_046846021.1 | XP_046845727.1<br>XP_046845733.1 |

**Table-S2: Species code and accession numbers of the NOX sequences**

See the corresponding phylogenetic analyses in **Figure S6; supplementary data** to **Figure 2D**.

| *H. vulgaris Jussy AEP* | *H. vulgaris* | Gene name: | Forward primer: | Reverse primer: |
|---|---|---|---|---|
| seq37733 | c19962_g1_i01 | ***SOD1 (CuZn-SOD)*** | TCAGGTGGTTGTATGAGTACTGG | CAATGTTACCTAGATCACCTGCAT |
| seq69497 | c20941_g1_i02 | ***SOD2 (Mn-SOD)*** | GGAGGAAAACCAACAGGTGA | GCACTGCAACAGCTGAAGAC |
| seq12246 | c12652_g1_i01 | ***SOD3 (CuZn-SOD)*** | GTAAACTGCGCTCCCGTATC | GCAATTCGACAAGTGCAACA |
| seq49098 | c19292_g1_i01 | ***CYBA*** | GAATGGGCTATGTGGGCTAA | GCATGAATGGAAGAGCATGA |
| seq42412 | c16180_g1-i04 | ***NOXA-like*** | CTTTGGCGTGTCGAAATTCT | GCCAACAAGAACACCTAACCA |
| seq48684 | c15638_g1_i01 | ***NOX2 / CYBB*** | GCTGGAGATTGGACTGAAGC | GGACCATCAACTGCCAGACT |
| seq32691 | c19363_g1_i03 | ***NOX4*** | GGGGTTGTCATGCTTGTTCT | TGTCAAATGGTGTGTGTACCAA |
| seq18159 | c18424_g1_i01 | ***Catalase*** | ACAGGCTGCTTTTTCTCCTG | ATCGATGGCGATGTGTATCA |
| seq65236 | c16197_g1_i01 | ***TBP*** | AAGCGATTTGCAGCAGTTAT | GCTCTTCACTTTTTGCTCCA |

| Gene name: | siRNA-a: | siRNA-b: | siRNA-c: |
|---|---|---|---|
| ***Scramble*** | AGGUAGUGUAAUCGCCUUG | / | / |
| ***SOD1*** | UCGACAUGCAGGUGAUCUA | AUCUGCUAUUUGUGUGUUA | / |
| ***SOD2*** | GAAUGUUCGUCUUGAUUAU | CAAUCAGCGGUCAAAUUAU | / |
| ***SOD3*** | UAACAUCAAAGGCGAAAUA | ACGAUAUCCUUACAUAGUA | ACUAUCAGAUGGCUGCAAA |
| ***CYBA*** | AUCUGGCCGUACAAUAGAA | AACAUUAACUAGAGCUCAA | / |
| ***NOXA-like*** | UCAGACAUAUGGUCGACAA | CAAGAGACCUGUUAACUUA | GUAUGGUAAAUGAUGUUGA |
| ***NOX2 / CYBB*** | ACAUUGGUAUCAAGUUGGA | UGUGGACACUUCGAAGAUA | CAUUGCAUCUUCUCAUAAA |
| ***NOX4*** | CAGCUUGCGGUUUCAUUUA | UGAUGGUGGUGAUGAAACA | UGAAGCGUAUGCAUUUGGU |
| ***Catalase*** | AAAUGGUGUUGGAUCGUAA | UGUGUUGGAUGAUAAAGAA | GCAUGUUGGUGCAAAUUUC |

### Table S3: Sequences of the primers and siRNAs used in this study

Primers used for Q-PCR analysis (upper table) and siRNAs electroporated in animals for gene knock-down (lower table). The *SOD1* and *SOD2* sequences are too short to design three distinct siRNAs. Accession numbers on HydrAtlas (hydratlas.unige.ch) are given.

| Drugs / RNAi Hv_AEP2 | | $mtO_2^{\cdot-}$ levels | $mbO_2^{\cdot-}$ levels | $H_2O_2$ levels | SOD activity | Catalase activity | Cell death | Wound healing | Apical Regener. | Basal Regener. |
|---|---|---|---|---|---|---|---|---|---|---|
| Control | BRt | ↗ | ↗ | ↗ | ↗ | ↗↗ | ↗ | 4 hrs | | 30 hrs |
| *Hv_AEP* | ARt | ↗ | ↗ | ↗↗ | ↗↗ | ↗ | ↗↗ | 4 hrs | 48 hrs | |
| Tiron | | ↘ 90% | ↘ | ↘ | nd | nd | ↘ 90% | delayed | delayed | nd |
| *SOD1*(RNAi) | | ↗↗ | ↗↗ | ↘ | | nd | nd | faster | nd | nd |
| *SOD2*(RNAi) | | ↗↗ | ↗↗ | ↘ | | | nd | faster | nd | nd |
| *SOD3*(RNAi) | | ↗↗ | ↗↗ | ↘ | | | nd | faster | nd | nd |
| *Catalase(RNAi)* | | | | ~ | | ↘ 90% | nd | nd | 48 hrs | 30 hrs |
| Ac-DEVD-CHO | | nd | | ↘ | nd | nd | inhibited | nd | nd | nd |
| DPI | | → | ↘ | ↘ | nd | nd | nd | 4 hrs | blocked | nd |
| HU | | → | | ↘ | | | reduced | nd | delay ++ | |
| ATZ | | | | ↘ | | ↘ 65% | nd | nd | blocked | nd |
| *NOX2/CYBB*(RNAi) | | | ↘ | | | | | | delay ++ | |
| *NOX4*(RNAi) | | | ↘ | | | | | | delay ++ | |
| *NOXA-like*(RNAi) | | | ↘ | | | | | | delay + | |

### Table S4: Table summarizing the various effects induced by modulations of ROS metabolism after drug treatments or gene silencing in *Hv_AEP2*

AR: Apical Regeneration ; ARt: apical-regenerating tips ; BR: Basal Regeneration ; BRt: basal-regenerating tips ; hpa: hours post-amputation ; $mtO_2^{\cdot-}$: mitochondrial superoxide; $mbO_2^{\cdot-}$: membrane superoxide; pa: post-amputation. Black arrows indicate observed modulations, red arrows indicate expected modulations in the absence of experimental data.

### Supplemental Figures

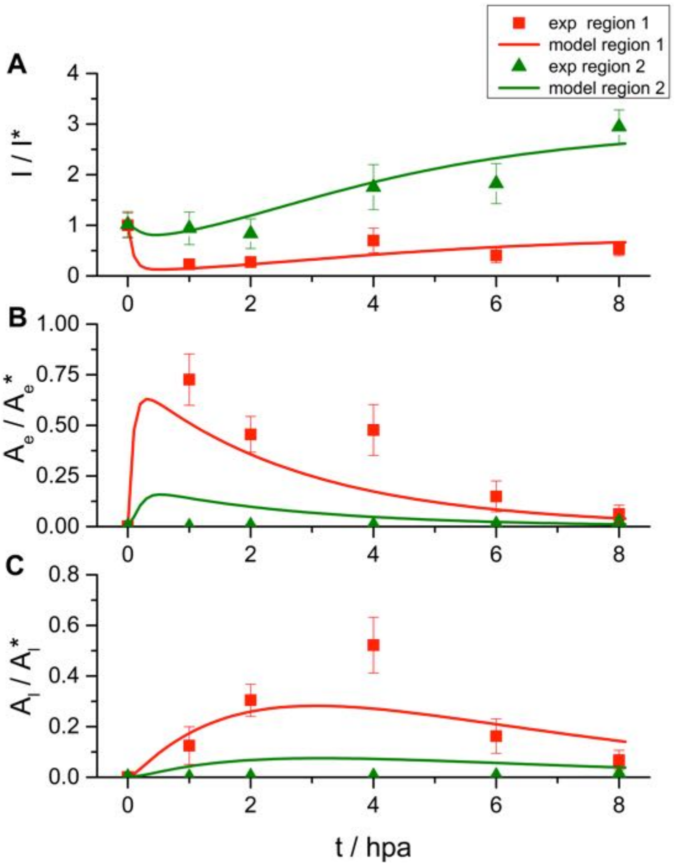

**Figure S1: Fitting the model to previous experimental data.**

Time course of density of interstitial (I, panel A), early apoptotic ($A_e$, panel B) and late apoptotic cells ($A_l$, panel C) in region 1 (closer than 100 µm to the amputation plane, in red) and region 2 (200 µm underneath, in green). Each density is normalized by the typical density values (see Supplementary Methods section). Dots and error bars correspond to the experimental data (average and standard deviation, respectively) while continuous curves depict the best fitting model simulation.

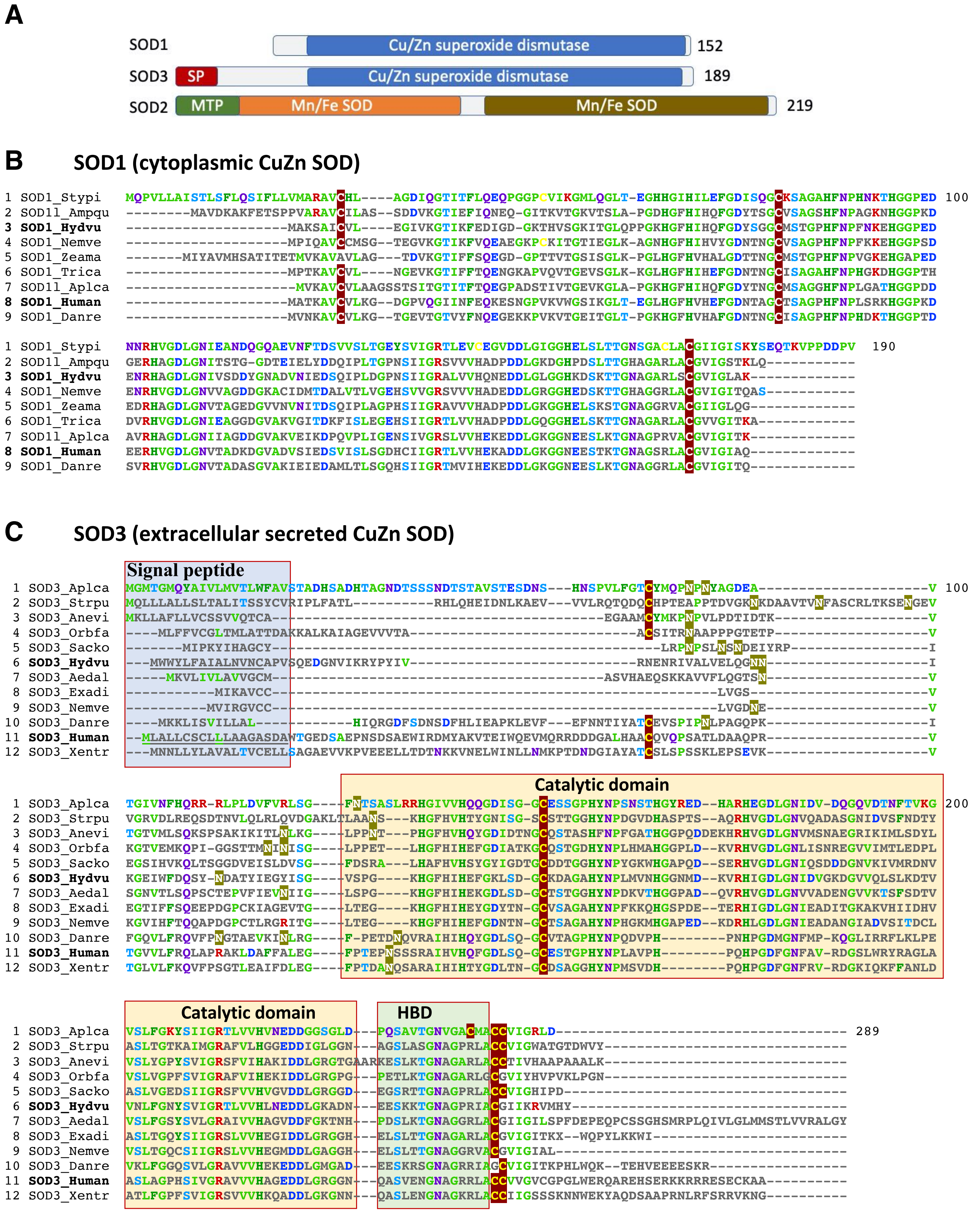
A
SOD1
Cu/Zn superoxide dismutase
152
SOD3
SP
Cu/Zn superoxide dismutase
189
SOD2
MTP
Mn/Fe SOD
Mn/Fe SOD
219
B
SOD1 (cytoplasmic CuZn SOD)
C
SOD3 (extracellular secreted CuZn SOD)
Signal peptide
Catalytic domain
Catalytic domain
HBD

**D SOD2 (mitochondrial Mn SOD)**

```
                 Mitochondrion transit peptide
1 MnSOD_Trica   ---------MFALRRLATAP---LAGSRAAHSLPELPYAYEALEPVISRDIMCLHHSKHHQTYVTNLNAAEEKLKAALSKGDISTAISLEPALRFNGGGH  100
2 SOD2_Hydvu    MFS-------FGIHHLSVFRKISRIAFANKHTLPELGYEYNALEPTISGQIMEIHHRKHHQAYVNNLNTAEEQLAEAQHKGDTSKIISLAPALKFNGGGH
3 SOD2_Human    MLSRA----VCGTSR-QLAPVLGYLGSRQKHSLPDLPYDYGALEPHINAQIMQLHHSKHHAAYVNNLNVTEEKYQEALAKGDVTAQIALQPALKFNGGGH
4 MnSod_Ampqu   MLSATRSFLLFGGKN-SLSS-LSSISRRHKHVLPELPYGYKALEPVISGDIMELHHTKHHATYVNNLNATEGKMKECLEAGDVSGAVALEGAYRFNGGGH

1 MnSOD_Trica   LNHSIFWQNLSPE-STQPSDALKKAIEDSFGGVQQLKDQLSASSIGVQGSGWGWLGYCSKSGKLKIATCANQDPLQATTGLVPLLGIDVWEHAYYLQYKN  200
2 SOD2_Hydvu    INHSIFWTNLSPNGGGKPTGELLEAIIKDFGSFEAMKTRLSSSAVAVQGSGWGWLGYDSVTKRLAITALPNQDPLQATTGLIPLLGIDVWEHAYYLQYKN
3 SOD2_Human    INHSIFWTNLSPNGGGEPKGELLEAIKRDFGSFDKFKEKLTAASVGVQGSGWGWLGFNKERGHLQIAACPNQDPLQGTTGLIPLLGIDVWEHAYYLQYKN
4 MnSod_Ampqu   INHSIFWNNLSPNGGGTPQGKLMEAIERDFGSFDEFKSQLTARTVAIQGSGWGWLGFNQVTGRLQIATCPNQDPLQATTGLVPLLGIDVWEHAYYLQYRN

1 MnSOD_Trica   VRADYVKAIFEIVNWKDVSQRFEKAKSC  228
2 SOD2_Hydvu    VRLDYVNAIFNIIDWKNVSARFVAAK--
3 SOD2_Human    VRPDYLKAIWNVINWENVTERYMACKK-
4 MnSod_Ampqu   VRPDYVKAIWDVINWDDVSKRLP-----
```

**E CCS (Copper Chaperone SOD)**

```
1 CCS_Trica   --------MPESQIEFAVQMTCNSCVEAVKKSLAGDPNIKNVNVDLEKGSVVVTSTLPTLQIQEKLESTGRKVVVRGYAGS--------SAGVAIVDTGK  100
2 CCS_Human   MASDSGNQGTLCTLEFAVQMTCQSCVDAVRKSLQGVAGVQDVEVHLEDQMVLVHTTLPSQEVQALLEGTGRQAVLKGMGSGQ---LQNLGAAVAILG-GP
3 CCS_Hydvu   ------MENLFDKMEFSVNMTDSSCVDKVSSSLDQLQGIKSFEVDLDKQSIIVTTNLPSSIVQESLESTGMLAVYRGQGEN----SVNLGAAVAILK-ND
4 CCS_Ampqu   --------MADTKMEFAVQMTCKSCEEAVKAALN-VPGINSVYVDVPNEVVIVETSLPSSNVHKLLESTGKLIVFRGFGGQEQAPTSHQGAAVVVMK-GS

1 CCS_Trica   GNIQGVVRFVQVAPSSCIIDGTIDGLEPSAEYKLTVNECGDISQGCESVGGVYQPLQDGEGRP-------YGDLGVFKTGNEGRATFKKEDNVLNLPEII  200
2 CCS_Human   GTVQGVVRFLQLTPERCLIEGTIDGLEPGL-HGLHVHQYGDLTNNCNSCGNHFNPDGASHGGP-QDSDRHRGDLGNVRADADGRAIFRMEDEQLKVWDVI
3 CCS_Hydvu   HQTYGLVRFVQKDLNSCIIDGSISKLSPFCKHAVHIHELGDLSNGCESTGDVYNPMPSTNEKV-------VGDLGNISXDXKGNSIFKYIDHXIKVWDVI
4 CCS_Ampqu   GPVNGLLRMVQVSSNECVIEGTIDGLTPNKEHLLKIHDHGDLSNGCESCGDVYNVMMSKNGKSVSHSLPPVGDIAALQSDGSGRISFQTKSERVKVYDVI

1 CCS_Trica   GRSLVVNSGE-------------------KRLVCGIIARSSGLFQNPKTICACDGVTIWDEK------------IPPRAVL   280
2 CCS_Human   GRSLIIDEGEDDLGRGGHPLSKITGNSGERLACGIIARSAGLFQNPKQICSCDGLTIWEERGRPIAGKGRKESAQPPAHL
3 CCS_Hydvu   GRSVCLHEKDVDLKTSKHS------DAGESIACGIIARSAGMLENSKKVCTCSGKTLWEER------------EETRVH-
4 CCS_Ampqu   GRSMILHSSIPSV------------YGTRRLMCGIIARSAGLFQNTKKVCTCDGVTIWDEA------------ASQRQK-
```

**F Catalase (Cata)**

```
1 Cata_Hydvu   -MANRPKSANQLQEYRDK---NHAAALTTEAGSPMETNTASMTVGPNGPLLIQDNNFIENMAHFDRERIPERVVHAKGAGAFGYFEVTHDVSKYTKLAAL  100
2 Cata_Trica   -MSFVDKAANQLVDYKNN--SKNDSAATTGHGTPIGEKDATLTVGPRGPILLQDNNFIEELAHFDRERIPERVVHAKGAGAFGYFEVTHDITKYTAAKVF
3 Cata_Ampqu   MSSKRPACFSQLEEYAKK--TQPGEVLTTTHGNPIDFKTAIQTFGPRGPMLMQDGVYLDEMAHFDRERIPERVVHAKGAGAYGVFEVTHDITKYCCAKLF
4 CATA_Human   MADSRDPASDQMQHWKEQRAAQKADVLTTGAGNPVGDKLNVITVGPRGPLLVQDVVFTDEMAHFDRERIPERVVHAKGAGAFGYFEVTHDITKYSKAKVF

1 Cata_Hydvu   NGVGKKTRMVVRFSSVGGESGSADTARDPRGFALKFYTEEGNWDLVGNNTPIFFIRDPILFPSFIHTQKRNPKTHLKDPDMFWDFISLRPETTHQVMFLF  200
2 Cata_Trica   SNIGKRTPIAIRFSTVAGESGSADTVRDPRGFAMKFYTEDGIWDLVGNNTPIFFIRDPILFPHFIHSQKRNPQTHLKDANMFWDFITLRPETTHQTI---
3 Cata_Ampqu   SEVGKKTDLFIRFSTVGGESGSADTARDPRGFAVKFYTEDGNWDLVGNNTPIFFIRDPFLFPSFIHTQKRNPVTHLKDPDMFWDFISLRPETTHQVSFLF
4 CATA_Human   EHIGKKTPIAVRFSTVAGESGSADTVRDPRGFAVKFYTEDGNWDLVGNNTPIFFIRDPILFPSFIHSQKRNPQTHLKDPDMVWDFWSLRPESLHQVSFLF

1 Cata_Hydvu   SNRGTPASYRHMNGYGSHTFKLVNKDGEACWCKFHFKTNQGIKNLSDEEAGLLAGQNPDYYIQDLYDSIEEGNFPSWTMYLQIMSLEQAKKHPVNPFDIT  300
2 Cata_Trica   ----------HMNGYGSHTFKLVNAEGKAVYCKFHHKTDQGIRNLDVQKAAQLASTDPDYGLRDLYNAIANGNYPSWTTYIQVMTMEQAAACKFNPFDLT
3 Cata_Ampqu   SDRGIPDGYRHMNGYGSHTFKLVNSKGEPVYCKFHYKTDQGIKNMPVGKAAELAGTNPDYSIQDLYEAIATGNFPSWTLSIQVMTYEQAEKCSFNPFDLT
4 CATA_Human   SDRGIPDGHRHMNGYGSHTFKLVNANGEAVYCKFHYKTDQGIKNLSVEDAARLSQEDPDYGIRDLFNAIATGKYPSWTFYIQVMTFNQAETFPFNPFDLT

1 Cata_Hydvu   KVWSHSEYPLIQVGKMVIDRNPENYFADVEQAAFSPAHMPPGVETSPDKMLQGRLFSYGDTHRHRLGCNYQQLPVNCPLKGQ-HNYQRDGPQAFN-NQGS  400
2 Cata_Trica   KVWPHGEYPLIPVGKFTLNRNPENYFAEVEQIAFSPAHMIPGIEPSPDKMLLGRLFSYGDTHRHRLGANYLQLPVNCPFKVR--NYQRDGPQAIN-NQGG
3 Cata_Ampqu   KVWPHADYPLIPVGKITLNRNPSNYFFDVEQSAFSPAHMPPGIEASPDKMLQGRLFSYDDTHFHRLGPNFQMIPVNCPYAGKPRNYVRDGPMCVDGNQGG
4 CATA_Human   KVWPHKDYPLIPVGKLVLNRNPVNYFAEVEQIAFDPSNMPPGIEASPDKMLQGRLFAYPDTHRHRLGPNYLHIPVNCPYRARVANYQRDGPMCMQDNQGG

1 Cata_Hydvu   APNYFPNSFNGPQEQRQFSNHVDSY--NGECRRYQSGDEDNFSQPKLFWENVLDDKEKSDLISNIAGHLKNAQEFIRKRVIKNFSDVHQDFGRRLNEAID  500
2 Cata_Trica   APNYHPNSFNGPDGDSRAKALNVPFAVSGDANRYDSGDEDNYSQSRVFYQKTLDQGARDRLIANIVGNLKDAADFIQERAIRNFTQVDAGLGQKIAEGLK
3 Cata_Ampqu   APNYYPNSFNGPKDMGKHDVTIFPGP-AGDVKRYNAADDDNFSQVGIFYNKVLNEEERTRLAQNIAGHMKNASPKIQERAIANFSKADPDYGARIKKYIS
4 CATA_Human   APNYYPNSFGAPEQQPSALEHSIQY--SGEVRRFNTANDDNVTQVRAFYVNVLNEEQRKRLCENIAGHLKDAQIFIQKKAVKNFTEVHPDYGSHIQALLD

1 Cata_Hydvu   ELVKG-----------------TTSAKANL   529
2 Cata_Trica   K----------------------QKRANL
3 Cata_Ampqu   Q-----------------------------
4 CATA_Human   KYNAEKPKNAIHTFVQSGSHLAAREKANL
```

**Figure S2: Alignment of superoxide dismutases (SOD) and catalase protein sequences**

**(A)** Schematic representation of the structure of the *Hydra* SOD1 , SOD3 and SOD2 proteins that are expressed at the cytoplasmic, extracellular and mitochondrial levels respectively. SOD1 and SOD3 belong to the Copper Zinc SOD protein family (CuZnSOD), SOD2 to the manganese SOD protein family (MnSOD). **(B)** Alignment of SOD1 sequences. **(C)** Alignment of SOD3 sequences with the catalytic domain (yellow frame) and a short Heparin Binding Domain (HBD, green frame) at the C-terminus. The signal peptide at the N-terminus (blue frame) was identified with SignalP-6.0; its sequence is underlined in the human and *Hydra* sequences. **(D)** SOD2 sequences encode a mitochondrion transit peptide (MTP, underlined) at the N-terminus identified with TargetP-2.0. **(E)** Alignment of Copper Chaperone for SOD1 (CCS) sequences. **(F)** Alignment of catalase sequences retrieved from Uniprot (www.uniprot.org): *Hydra* A0S5U0_HYDVU, Human P04040, *Tribolium* A0A139WLA9_TRICA, *Amphimedon* A0A1X7VNW3_AMPQE. In each panel, sequences were aligned with Muscle align, subsequently viewed with Mview at EBI. In each alignment, the reference sequence is (1), identities are normalised by aligned length and residues colored by identity. Accession numbers to SOD and CCS sequences are given in **Table-S1**.

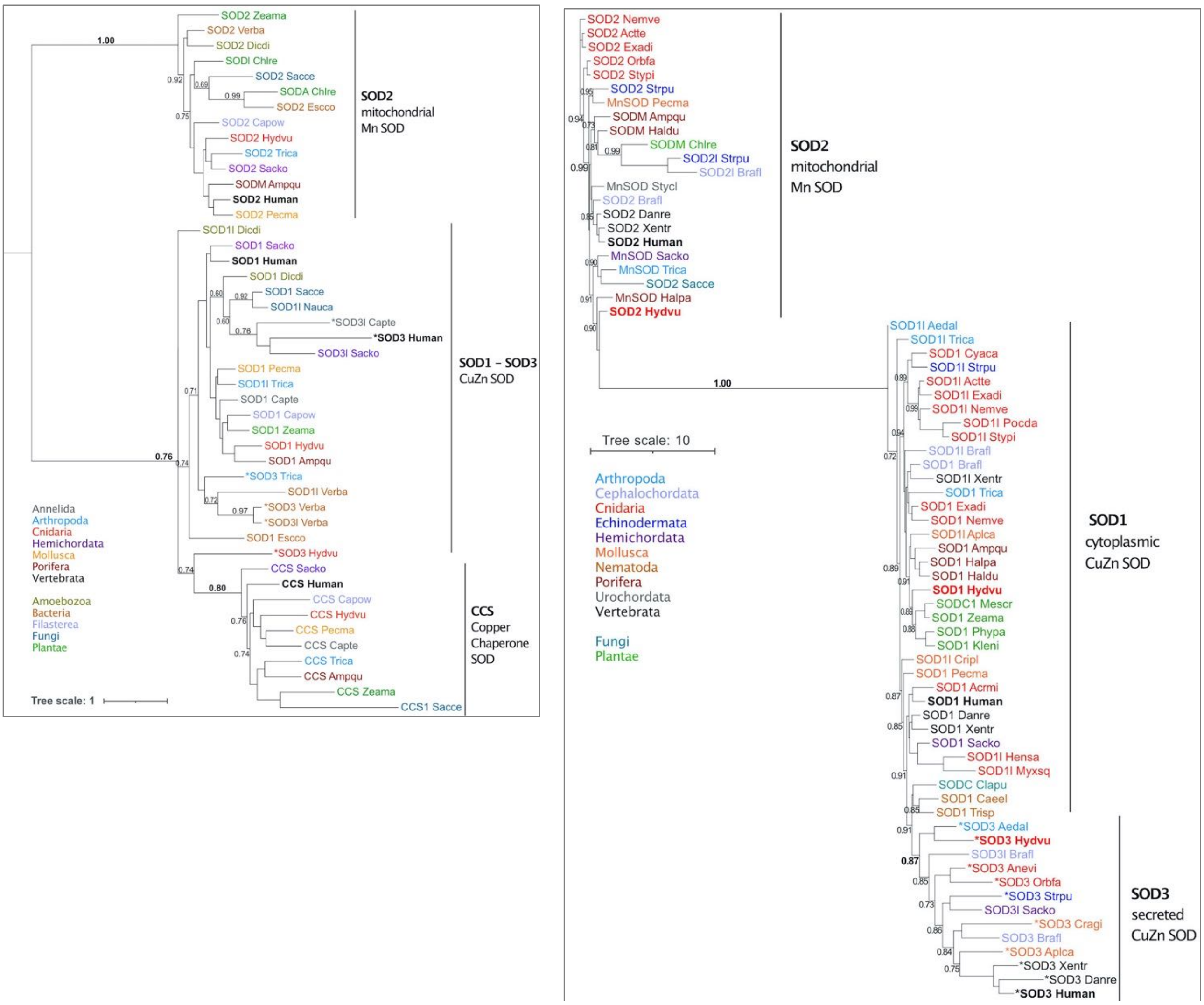

**Figure S3: Phylogenetic analysis of the *Super Oxide Dismutase (SOD)* gene families**

Phylogenetic reconstruction performed with PhyML3.0[4] on the ATGC facility using the LG substitution model on sequences aligned with Muscle Align. The branch support was assessed with the aLRT SH-like fast likelihood method[5]. The trees show the phylogenetic position of 46 SOD1, SOD2, SOD3, CSS sequences (tree on the left hand) and 77 SOD1, SOD2, SOD3 sequences including all available sequences from cnidarian species (written red), either anthozoan (*Acrmi, Actte, Nemve, Orbfa, Stypi)* or medusozoan (*Hydvu, Cyaca*) (tree on the right hand). See species code and accession numbers in **Table-S1**.

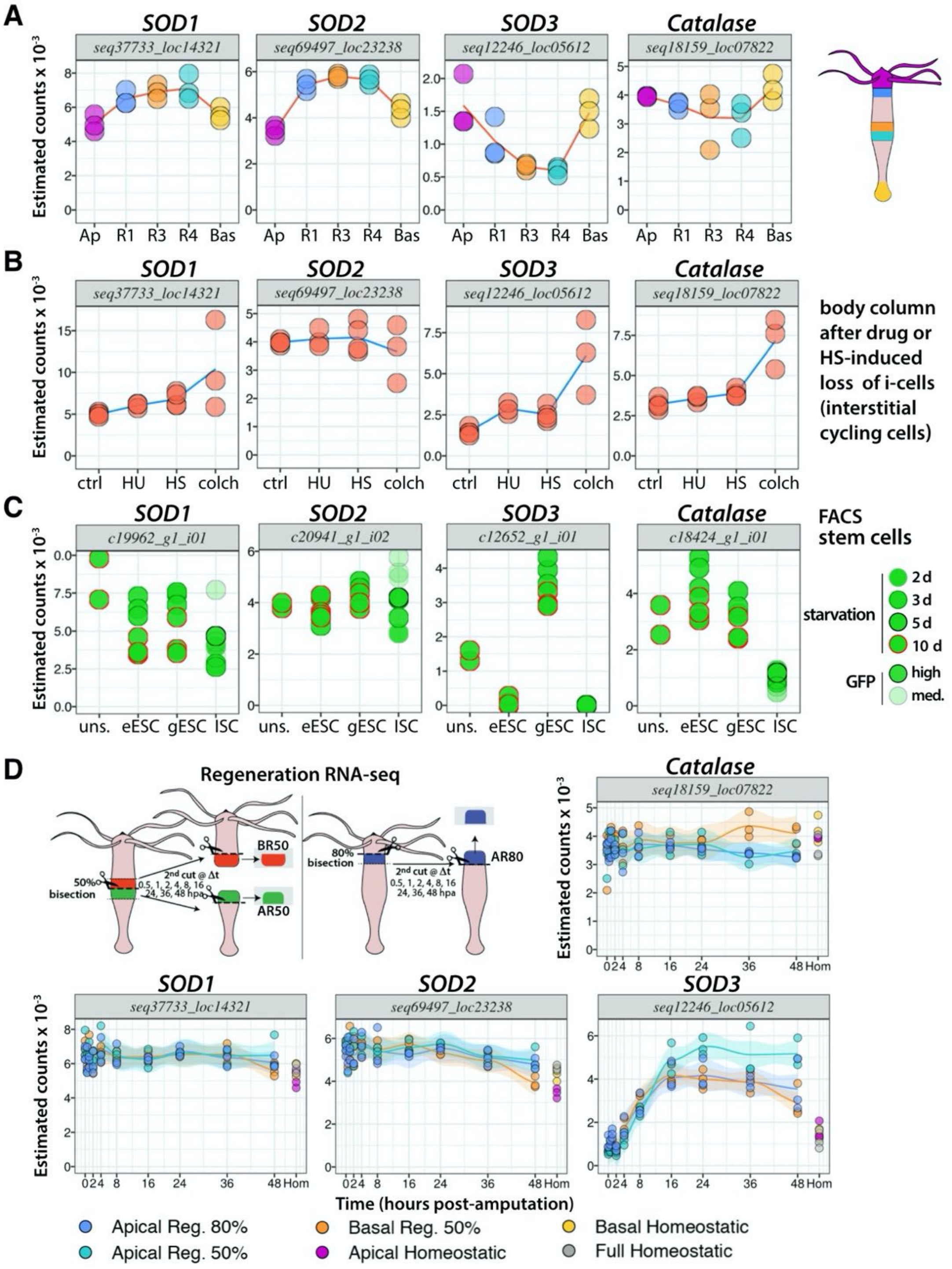

**Figure S4: RNA-seq expression profiles of the *Hydra SOD1, SOD2, SOD3* and *catalase* genes**

**(A)** Estimated counts at five positions along the body axis of *Hv_Jussy* animals: Ap: apical; R1: upper body column; R3 and R4: upper and lower mid-gastric regions; Bas: basal region. **(B)** Estimated counts in the body column of *Hv_sf1* animals 7 days after drug (HU: hydroxyurea, colch: colchicine) or heat-shock (HS) treatments that eliminate the cycling interstitial cells (ISCs and interstitial progenitors); ctrl: control body columns from untreated animals. **(C)** Estimated counts in flow cytometry sorted stem cells from transgenic *Hv_AEP* animals that constitutively express GFP, either in their epidermal Epithelial Stem Cells (eESC), or gastrodermal Epithelial Stem Cells (gESC), or Interstitial Stem Cells (ISC); uns: unsorted cells from the body column of non-transgenic *Hv_AEP* animals. **(D)** Estimated counts from apical or basal-regenerating tips (AR, BR respectively) collected at indicated time-points after mid-gastric bisection (50%) or decapitation (80%); hpa: hours post-amputation. All sequences and RNA-seq profiles are available on HydrAtlas. (hydratlas.unige.ch/blast/blast_link.cgi). For methodological details see in refs.[6,7].

**A** **CYBA (Cytochrome b-245 light or alpha chain)** Alternative names: alpha-subunit p22, p22phox

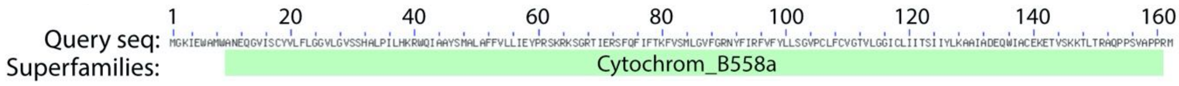

Reference sequence (1): CYBA_Hydvu - Identities normalised by aligned length. Colored by: identity

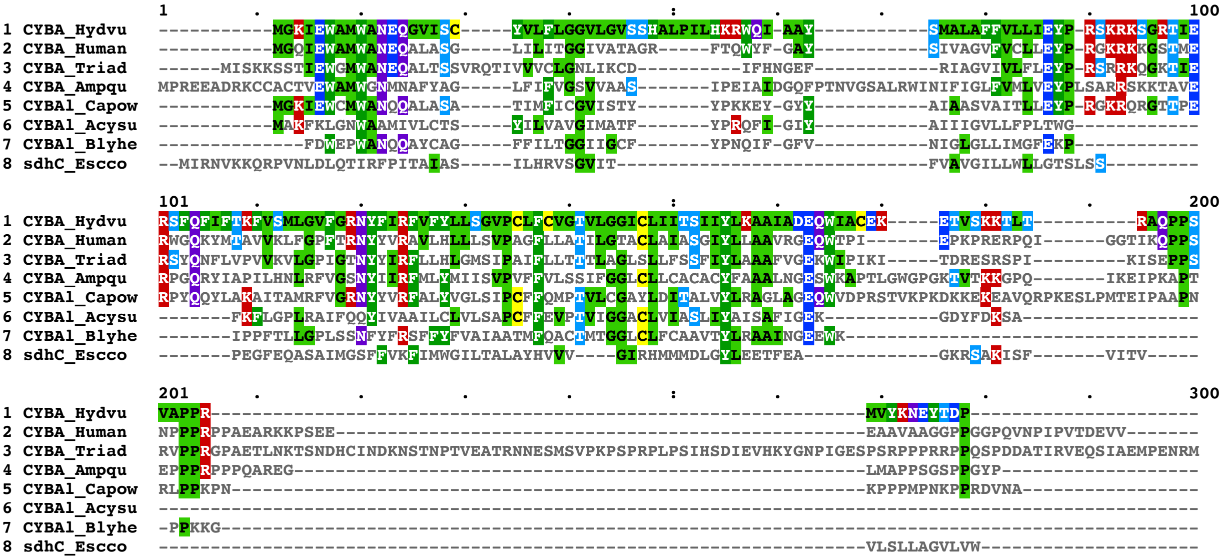

**B** **NOXA-like**

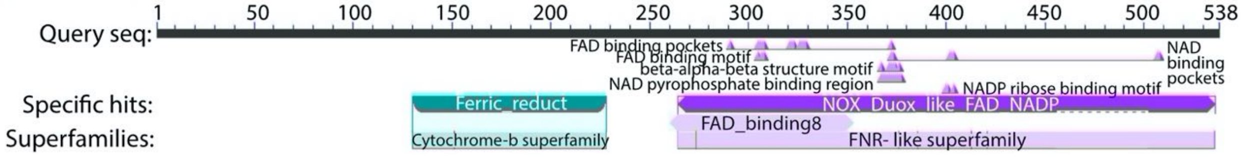

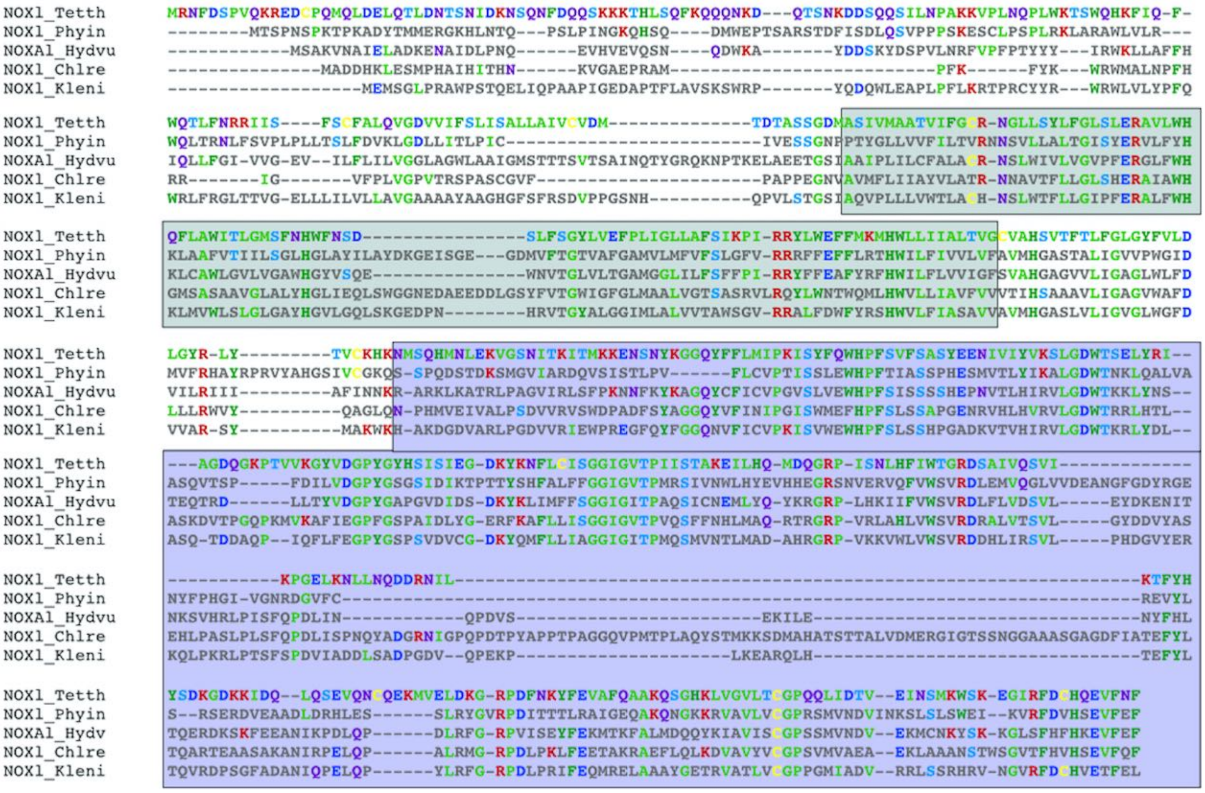

### C NOX2/CYBB (Cytochrome b-245 heavy chain) and NOX4

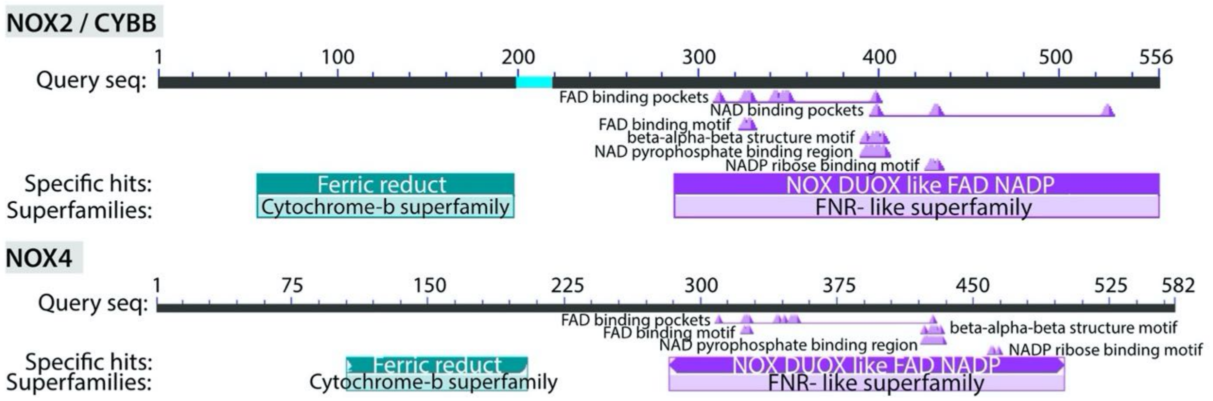

```
              1         .         .         .         .         :         .         .         .         .       100
NOX4_Hydvu    ----------------------------MGINIRSMVQNNFLKTIILAGWLTINIYLLTSTYLLYYKSDKYHYLYMVLKESICVSRACAMAINFNLALIL
NOX4_Human    ----------------------------MAVSWRSWLANEGVKHLCLFIWLSMNVLLFWKTFLLYNQGPEYHYLHQMLGLGLCLSRASASVLNLNCSLIL
NOX1l_Dicdi   ------------------------MRLPTKEEIQRYWVNEGNKLILVILYTLGNIAAFVYTFVHYYNSPAF----EVVGYGVCFARGCAQLLKLNCALIL
CYBB_Ampqu    ----------------------------MSLLPSGWIINEFPKWLVFGTWFLLNIVLFIYQFNLFQTSNTQFYMRFYLKEALAFARGPALPINFNVILIL
NOX1_Batde    MPNSRPQQTSFQNELSLGAVRPRQRKSGWMVTFNNWMVNEGRSRLFFALFIASQLGYFAWSYYQLWTSPTLVTFRTVLQHGLPMARAAANVINLDCGIIL
NOX1_Capow    ---------------------MVFGLPEGKAEWTKFLWNEGPALLFFVTWLCANIGLFFSTYSSYQTNEDYKCPHDIMGAGLSVARGAAMILNLNCALIL
CYBB_Hydvu    ---------------------------------MGWFVNEFPKYFVMVIWVGLNLMLFLLTYFQYTDGEKYYYLRYIIKGGLPVARGAAAMLNFNCMLIL
CBB_Human     --------------------------------MGNWAVNEGLSIFVILVWLGLNVFLFVWYYRVYDIPPKFFYTRKLLGSALALARAPAACLNFNCMLIL

              101       .         .         .         .         :         .         .         .         .       200
NOX4_Hydvu    IPMCKTIISWIRTKLLKYLHSNPRRLV--NHLKGLHILCAFTMCILSVIHTLSHLVNSLRF-----HLNFT------------EAIEAINVASSKK-ETT
NOX4_Human    LPMCRTLLAYLRGS-QKVPSRRTRRLL--DKSRTFHITCGVTICIFSGVHVAAHLVNALNF-----SVNYS------------EDFVELNAARYRD-EDP
NOX1l_Dicdi   VPVLRNLLSFLRGT-------FLNNYVPFDKNIVFHKLIAWVICFATFGHVMAHFNNFRLY-----QDITP------------QEYKRILGIDYPN-LTP
CYBB_Ampqu    LPVCRNIISVIRGC-TRVVPRSIRRVL--DKNLTFHKAIAWMLVVSSTMHVVAHYYNYERL-----ARYSPPNSLPPGDQATSIPTQALPFNGSVGQLNP
NOX1_Batde    FTVCRNVISLMRTT-------FLNRIIPFDKNITFHIWIAYSIAFWTFVHVIAHYFNYNNV--------------------------RIALG-----VSA
NOX1_Capow    LPVCRNLLSLVRGS----IRLSFIRLL--DHNITMHKLIAWTICLAAVVHTGAHFFNFLNL----------------------EKTGSLQCGGQPG-TSA
CYBB_Hydvu    LPVCRNLLSFVREL--PFLPHSIGRIM--DKNITFHRNIAYMICFATALHICAHVFNIETLVSSW-QSKDPVIQYLNHLDDNILNQTRLNPIRGPE-LDP
CBB_Human     LPVCNLLSFLRGS-SACCSTRVRRQL--DRNLTFHKMVAWMIALHSAIHTIAHLFNVEWCVNARVNNSDPYSVALSELGDR-QNESYLNFARKRI-KNP

              201       .         .         .         .         :         .         .         .         .       300
NOX4_Hydvu    ---LWLVLSKVPGWTGVVMLVLLAIIVLTSFQAVRRQNYEVFWYTHHLTIVFLILACFHGFGKITK-FQ---TNLDKNPRECHSLVRKMW----------
NOX4_Human    ---RKLLFTTVPGLTGVCMVVVLFLMITASTYAIRVSNYDIFWYTHNLFFVFYMLLTLHVSGGLLK-YQ---TNLDTHPPGCISLNRTSSQNISLPEYFS
NOX1l_Dicdi   ---IKYAFATLAGWTGHVVCIVMVLMYTSAVESIRRPMFEGFWYTHHLFVVFFGLLVVHGLHSILE----------------------------------
CYBB_Ampqu    ---IHACFITYAGVTGHIITLVLFLMITSSVTYIRRSFFEVFWYTHQLFIVFLVGLGIHQYGRLLPVFRSASQLQDVGLGVCAYAPTS------------
NOX1_Batde    ---EYLSLVSGPGLTGQVISVSFFLIFTSAMEAVRRKYFEIFWFTHHLFLVFFGALLMHGSFCFIK-GD--------SGDPCRG----------------
NOX1_Capow    ---EDYAFATIAGSTGHIVVLVMFLMFTSSMSIIRRSYFEIFWYTHHLFVVFFVLLCIHGLGGLVK-KR-----VVIDPTDPANDV--------------
CYBB_Hydvu    ---IIELLKTVAGVTGVIITIALIVMVSSSTELIRRSYFEVFWFTHHLFVIFFVLLVFHGFGNLVR-SQ--NNVAQSNPEKCIKNFT-------------
CBB_Human     EGGLYLAVTLLAGITGVVITLCLILIITSSTKTIRRSYFEVFWYTHHLFVIFFIGLAIHGAERIVR-GQTAESLAVHNITVCEQKIS-------------

              301       .         .         .         .         :         .         .         .         .       400
NOX4_Hydvu    -------------------KENSTICLEAPSFESNVPQTWKWIVGPLCLYIIDRIIRLFRAVKDCQVANVILHEDKMIELHMLKKGFKSKPGQYILLQCP
NOX4_Human    EHFHEPFPEGFSKPAEFTQHKFVKICMEEPRFQANFPQTWLWISGPLCLYCAERLYRYIRSNKPVTIISVMSHPSDVMEIRMVKENFKARPGQYITLHCP
NOX1l_Dicdi   -----------------------------------PTSFWKWVIGPCALYIVERLIRLLRSKKTTMLIQARIHPSRVIEVRMKTERFKYKPGQYLFLNCP
CYBB_Ampqu    -------------------SENYTRFCTNLNFAPAGPTSWMFLLCGLLIYGLERILRVVRGFYSVVIIKVVQHPSNTIEIQMRRKGFHAEAGQYVFVNCP
NOX1_Batde    -----------------------------------GPMFWKFWVGSAAFYLIERLWREISGRRKTYIFKVVQHPSKVVEVQIKKNGWKMQAGQYIFICCP
NOX1_Capow    -------------------------------YESQPPNFWKFVVGPFGVYLIERGLRFYRYTQNTIVQKVVQHPSKVVEIQLVRRGFHSKPGQYIFVHCP
CYBB_Hydvu    -------------------------MCPLPQFSAKGPETWKWIVGPMFLYLIERCLRFYRSFETCEIKKVVKHPSRVVEFQLRKPGFKASAGQYIFLHCP
CBB_Human     -------------------EWGKIKECPIPQFAGNPPMTWKWIVGPMFLYLCERLVRFWRSQQKVVITKVVTHPFKTIELQMKKKGFKMEVGQYIFVKCP

              401       .         .         .         .         :         .         .         .         .       500
NOX4_Hydvu    DISDLEWHAFTLTKCPTNDGGDETFSVNMKVAGDWTKKLKAKFFKCINTKGIFMQRVGDNCIINIDKSIVYDNENEENDENVLPIIRVDGPYGSPNEDCF
NOX4_Human    SVSALENHPFTLTMCPTET--KATFGVHLKIVGDWTERFRDLLPP---------------------SSQDS-EILPFIQSRNYPKLYIDGPFGSPFEESL
NOX1l_Dicdi   TIAQNEWHPFTITSAPEED----FVSCHINVVGNWTGKLSTLLNPD--------------------KKMGIVQENVLKSPDGKPILRIDGPFGAASEEVF
CYBB_Ampqu    SVAWFEWHPFTLTSAPEED----YFSVHIRILGDWTRGIADKL-GM--------------------------NRNDFQQSWELPSIYIDGPFGTASEDAF
NOX1_Batde    EIGLFEWHPFTLTSSPHEE----FLSIHIRVVGDWTEKFAERV-GC--------------------RFGG--SSDNMPAPDTLPYVMVDGPYGSASEDVF
NOX1_Capow    AVSWFEWHPFTLTSAPEED----FASVHIRVAGDWTTKFAEAL-GC--------------------NWSNKGDAGEIQASATLPRVAFDGPFGTASEDVF
CYBB_Hydvu    SISPLEWHPFTLTSCPEED----FFSVHIRQAGDWTEALAKYCDGN--------------------------SNQKTVDTSKIPSLAVDGPFGTASTDVF
CBB_Human     KVSKLEWHPFTLTSAPEED----FFSIHIRIVGDWTEGLFNAC-GC--------------------------DKQEFQDAWKLPKIAVDGPFGTASEDVF

              501       .         .         .         .         :         .         .         .         .       600
NOX4_Hydvu    RYPVNISVAAGIGITPFAAVLNKL--RSLDPKLVDKNFKM-KRMHLVWACRDIQEFTWFLQLMFDTVKHLNSINMGDLLVCHLFVTRGANLTTEDPSMAD
NOX4_Human    NYEVSLCVAGGIGVTPFASILNTL----LDD---WKPYKL-RRLYFIWVCRDIQSFRWFADLLCMLHNKFWQENRPDYVNIQLYLSQTDGIQKIIGEKYH
NOX1l_Dicdi   KYKQVILVGAGIGVTPFASILKHIKYQMART---YNTTPLIDKVHFYWICRDRNSFEWFSGLIGELEME----NHNNFLEIHPYLTGALSAQEIRDVMYG
CYBB_Ampqu    EHPVALLVGAGIGVTPFASVLKSLYYRITEE---GSGLVL-KKLYFVWICPETYAFEWFANLFQELEQQLIDKGIDDFLTSWIYLTRGWKDNQAFYLMLN
NOX1_Batde    DYEAAVLVGAGIGVTPFASILKTIWFRINNP---TRAVPL-KKVYFFWICRDKDAFEWFQDLLSTIEDE----NISNFLEIHTYLTQKLKIFEVKNIVIN
NOX1_Capow    KYEVAVCVGTGIGVTPFASILKSIWYRLLNP---SIQLKL-RKVYFIWVCREKEAFEWFADLLEALEQQMEQQNLYDFLDARIYLTGELNSNEVRNVMLN
CYBB_Hydvu    YHENLMCIGAGIGITPFASILKSIWYRHHRG---DTTLKI-KKVYFYWICPDTNAFEWFSDLLQSLEDQMTEKGLLDFLEYNIYLTQGWGKGDVRNIVLH
CBB_Human     SYEVVMLVGAGIGVTPFASILKSVWYKYCNN---ATNLKL-KKIYFYWLCRDTHAFEWFADLLQLLESQMQERNNAGFLSYNIYLT-GWDESQANHFAVH

              601       .         .         .         .         :         .         .         .    687
NOX4_Hydvu    Y--------------KIQWLQKRLRYGRPDFEDILKKISLNHYKTRVGVFFCGPGPLGKVLYDACKSVNFR----GNSFIFHKESFA
NOX4_Human    A------------------LNSRLFIGRPRWKLLFDEIAKYNRGKTVGVFCCGPNSLSKTLHKLSNQNNSY----GTRFEYNKESFS
NOX1l_Dicdi   D----------EEKDLITGFTTPTQFGRPKWDEIFADHALRYAEKDVGVFFCGPKLLSKSLYKASTHYTKT---TTCRFHYNKENF-
CYBB_Ampqu    EGRVTAEHATDDMVDAITGLRAETTFGRPEWEKIFEKVALAHPNTDVGVFFCGPSVLSHNLHKQCNAFTKKGGSAGARFYYNKENF-
NOX1_Batde    --------EDGRDAITGLKSRTQYGRPNWDQIFEALRVKHRATDIGVFFCGPKVLSRTLHHTCNKWTEA-TEDGTRFYYGKENF-
NOX1_Capow    D---------EEEADAITGLRAKTNFGRPQWGRIYEQLVQDHPSTNIGVFYCGPKVVAKELGRLSKQFTNP-RPGGTIFNFNKENF-
CYBB_Hydvu    E---------SDSADVITGLRQKTNYGRPNWDRIFSNIASSHKGSNIGVFFCGPAVLSHVLHKMSNKYSGD----SVRFYYNKENF-
CBB_Human     H---------DEEKDVITGLKQKTLYGRPNWDNEFKTIASQHPNTRIGVFLCGPEALAETLSKQSISNSES-GPRGVHFIFNKENF-
```

**Figure S5: Structure of the *Hydra vulgaris* NOX-like proteins: NOXA-like, NOX2/CYBB, NOX4**

The three *Hydra* NOX-related proteins belong to the group of CYBA-dependent NADPH-oxidases. **(A)** Structure and alignment of **CYBA** (Cytochrome b(558) alpha chain) sequences from *Hydra* (T2MGH1_Hydvu), *Human* (P13498), *Trichoplax* (RDD47224.1), *Amphimedon* (XP_003387341.1), *Capsaspora* (XP_004344260.1), *Blyttiomyces* (RKO85547.1) and *Acytostelium* (XP_012748157.1). **(B)** Structure and alignment of **NOXA-like** sequences found in plants (*Klebsormidium, Chlamydomonas*), ciliophoran (*Tetrahymena*), fungi (*Phytophthora*) and cnidarians (*Hydra*) but not in bilaterians. **(C)** Structure and alignment of the **NOX2/CYBB** and **NOX4** sequences. Sequences were aligned with Muscle align, subsequently viewed with Mview at EBI. Supplementary data to Fig. 2B, Fig. 6E-6F See species code and accession numbers in **Table-S2**.

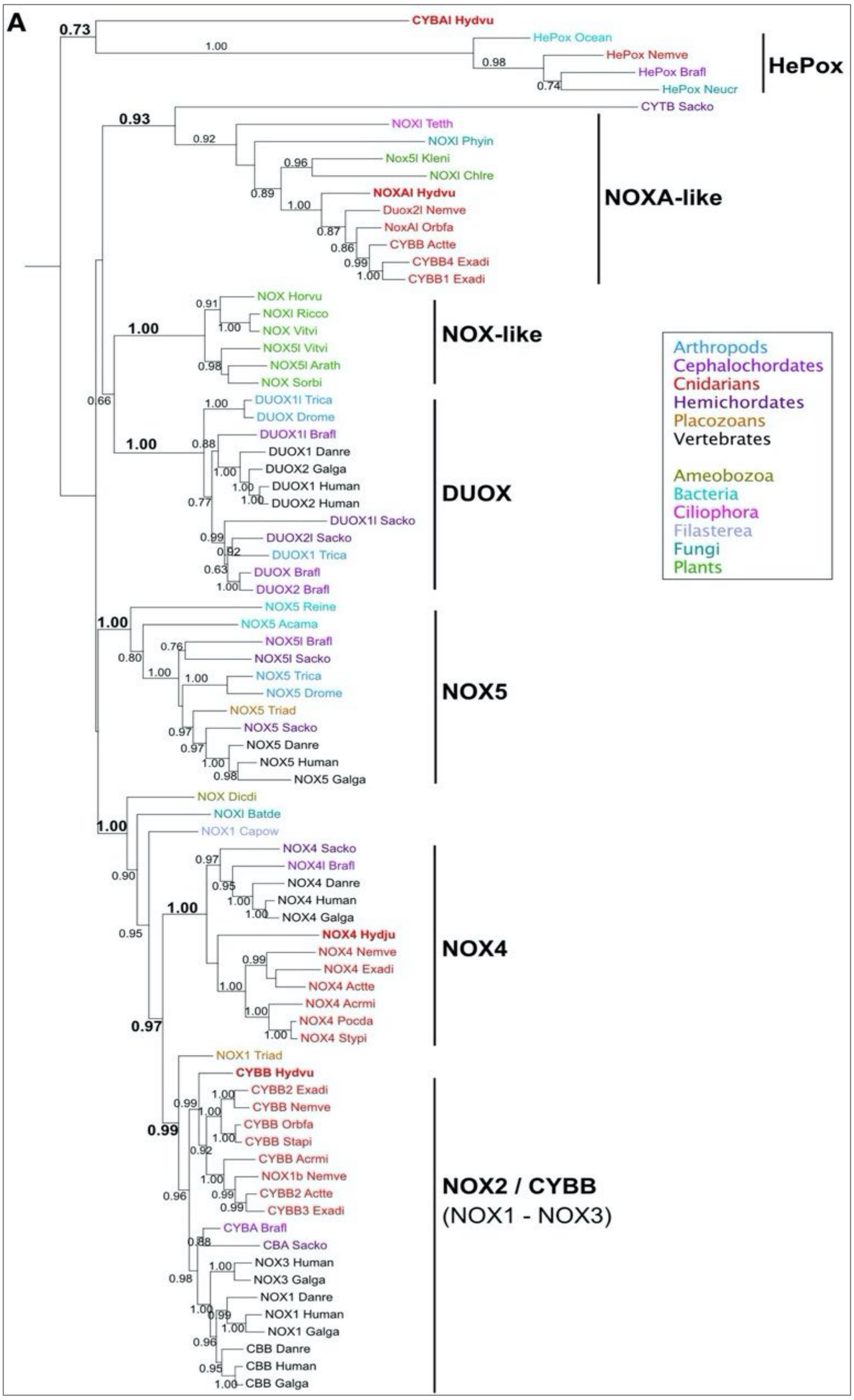

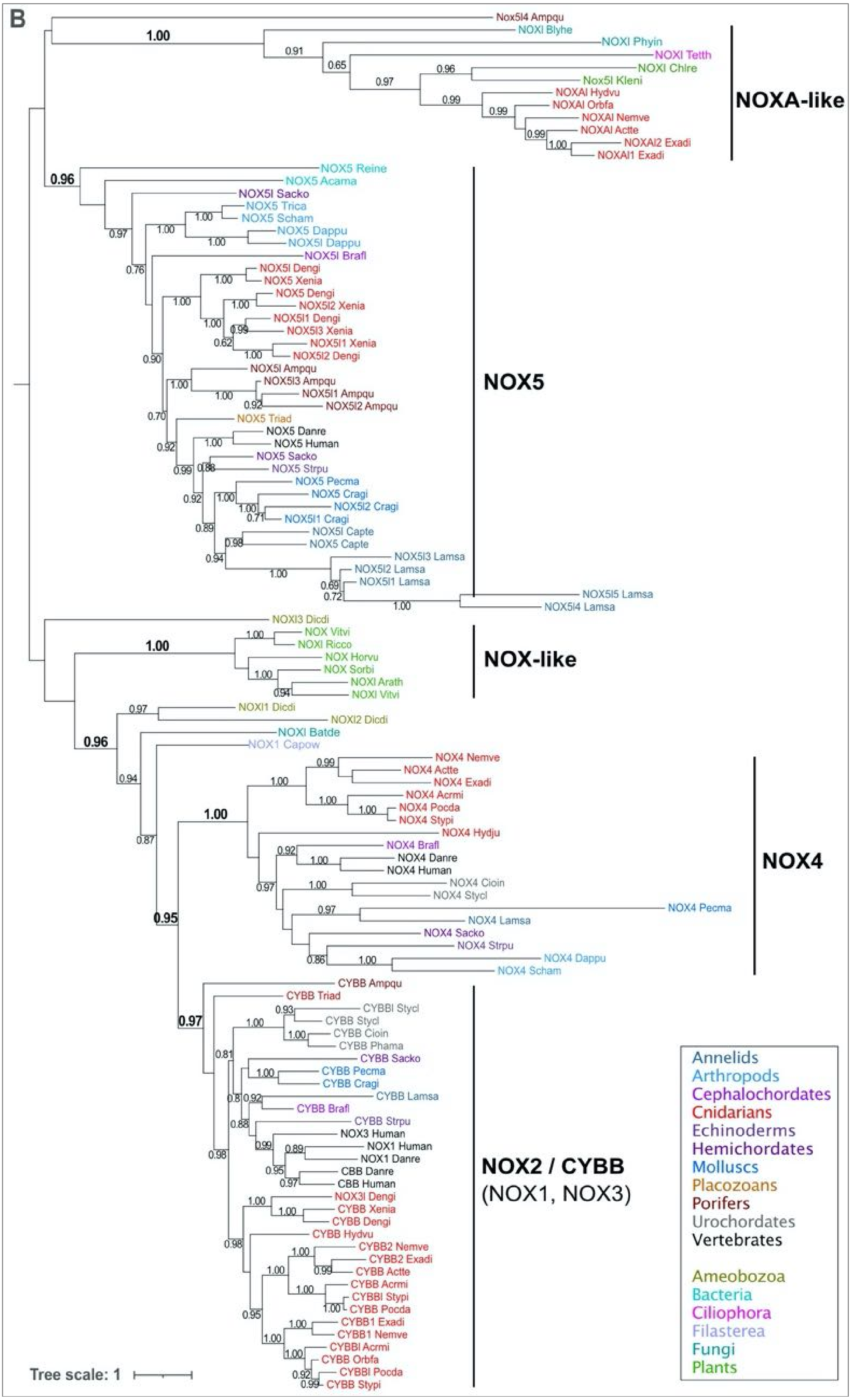

B
Nox5l4 Ampqu
NOXl Blyhe
NOXl Phyin
NOXl Tetth
NOXl Chlre
Nox5l Kleni
NOXAl Hydvu
NOXAl Orbfa
NOXAl Nemve
NOXAl Actte
NOXAl2 Exadi
NOXAl1 Exadi
NOXA-like
NOX5 Reine
NOX5 Acama
NOX5l Sacko
NOX5 Trica
NOX5 Scham
NOX5 Dappu
NOX5l Dappu
NOX5l Brafl
NOX5l Dengi
NOX5 Xenia
NOX5 Dengi
NOX5l2 Xenia
NOX5l1 Dengi
NOX5l3 Xenia
NOX5l1 Xenia
NOX5l2 Dengi
NOX5l Ampqu
NOX5l3 Ampqu
NOX5l1 Ampqu
NOX5l2 Ampqu
NOX5 Triad
NOX5 Danre
NOX5 Human
NOX5 Sacko
NOX5 Strpu
NOX5 Pecma
NOX5 Cragi
NOX5l2 Cragi
NOX5l1 Cragi
NOX5l Capte
NOX5 Capte
NOX5l3 Lamsa
NOX5l2 Lamsa
NOX5l1 Lamsa
NOX5l5 Lamsa
NOX5l4 Lamsa
NOX5
NOXl3 Dicdi
NOX Vitvi
NOXl Ricco
NOX Horvu
NOX Sorbi
NOXl Arath
NOXl Vitvi
NOX-like
NOXl1 Dicdi
NOXl2 Dicdi
NOXl Batde
NOX1 Capow
NOX4 Nemve
NOX4 Actte
NOX4 Exadi
NOX4 Acrmi
NOX4 Pocda
NOX4 Stypi
NOX4 Hydju
NOX4 Brafl
NOX4 Danre
NOX4 Human
NOX4 Cioin
NOX4 Stycl
NOX4 Pecma
NOX4 Lamsa
NOX4 Sacko
NOX4 Strpu
NOX4 Dappu
NOX4 Scham
NOX4
CYBB Ampqu
CYBB Triad
CYBBl Stycl
CYBB Stycl
CYBB Cioin
CYBB Phama
CYBB Sacko
CYBB Pecma
CYBB Cragi
CYBB Lamsa
CYBB Brafl
CYBB Strpu
NOX3 Human
NOX1 Human
NOX1 Danre
CBB Danre
CBB Human
NOX3l Dengi
CYBB Xenia
CYBB Dengi
CYBB Hydvu
CYBB2 Nemve
CYBB2 Exadi
CYBB Actte
CYBB Acrmi
CYBBl Stypi
CYBB Pocda
CYBB1 Exadi
CYBB1 Nemve
CYBBl Acrmi
CYBB Orbfa
CYBBl Pocda
CYBB Stypi
NOX2 / CYBB
(NOX1, NOX3)
Annelids
Arthropods
Cephalochordates
Cnidarians
Echinoderms
Hemichordates
Molluscs
Placozoans
Porifers
Urochordates
Vertebrates
Ameobozoa
Bacteria
Ciliophora
Filasterea
Fungi
Plants
Tree scale: 1

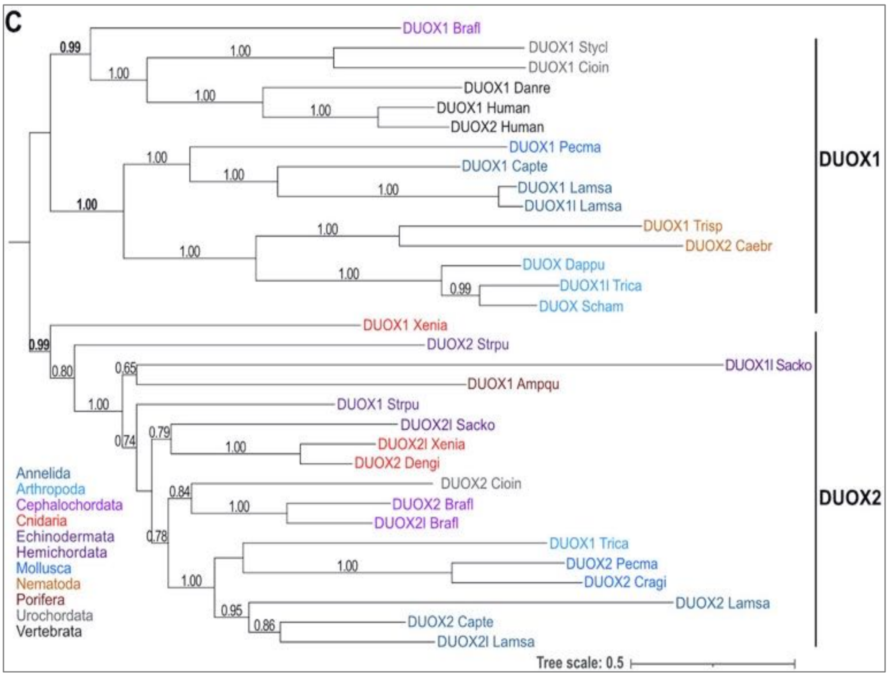

### Figure S6: Phylogenetic analysis of NOX sequences

Phylogenetic analysis of NOX-related sequences aligned with Muscle Align and submitted to PhyML3.0[4] at ATGC facility in Montpellier using the LG substitution model. The branch support was assessed with the aLRT SH-like fast likelihood method[5]; only values above 0.60 are shown. **(A)** Tree built on 81 NOX-related sequences that include DUOX sequences, rooted with Heme peroxidase (HePox) sequences from *Brafl* (XP_002605323.1), *Nemve* (XP_001637870.1), *Ocean* (ZP_01306662.1), *Neucra* (XP_001728273.1). **(B)** Tree built on 110 NOX-related sequences but no DUOX-related ones, rooted at midpoint. **(C)** Tree built on 32 DUOX-related sequences rooted at midpoint. Well supported nodes show aLRT values >0.80. See accession numbers and species code in **Table-S2**. Supplementary data to Fig. 2D

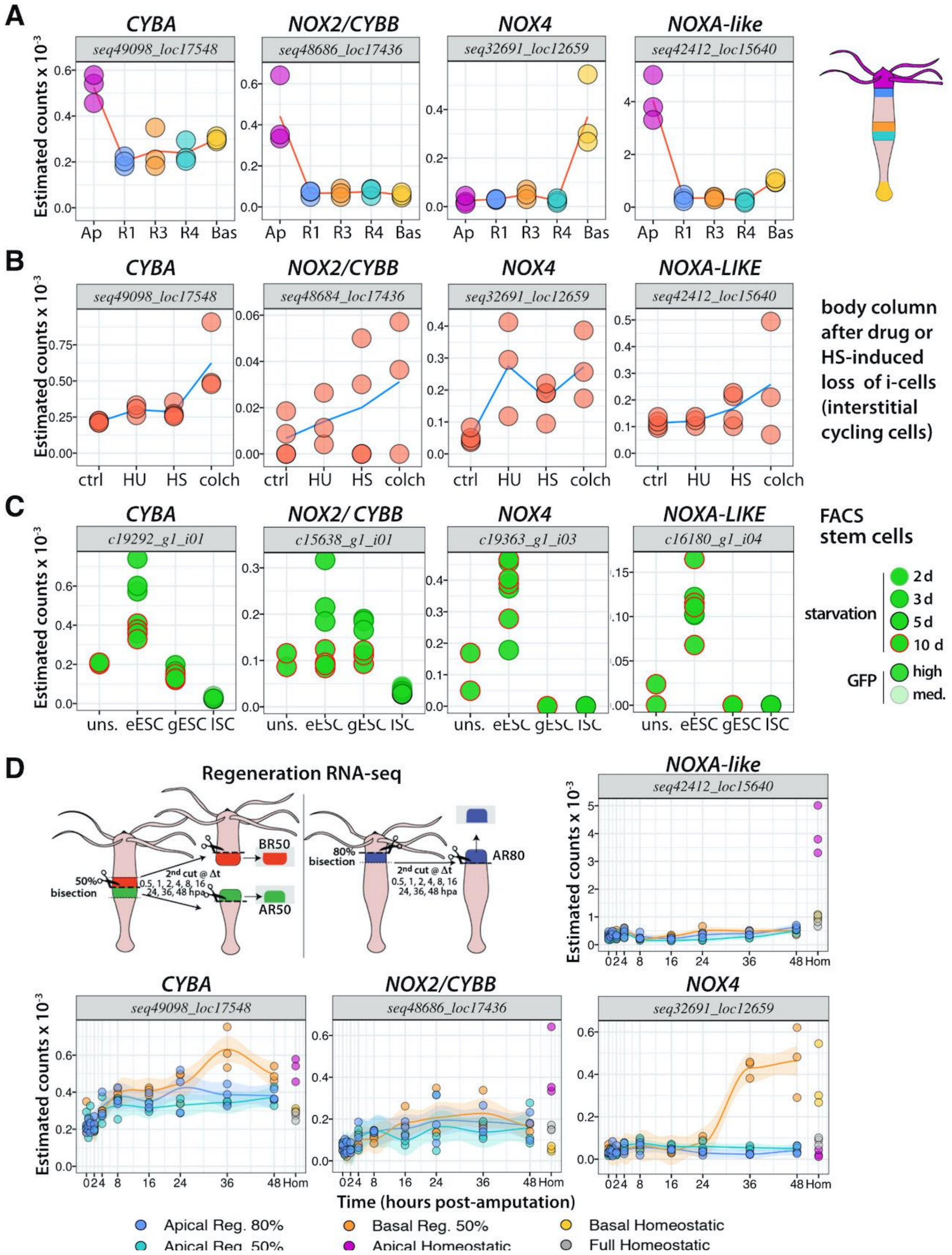

**Figure S7: Expression profiles of *Hydra* NOX-like sequences: *CYBA; CYBB/NOX2, NOX4, NOXA-like***

Note the higher level of *NOXA-like* expression when compared to *NOX4, CYBA, CYBB/NOX2* and the predominantly apical expression of CYBA, NOX2/CYBB and NOXA-like when NOX4 expression is predominantly basal and together with CYBA up-regulated during basal regeneration.

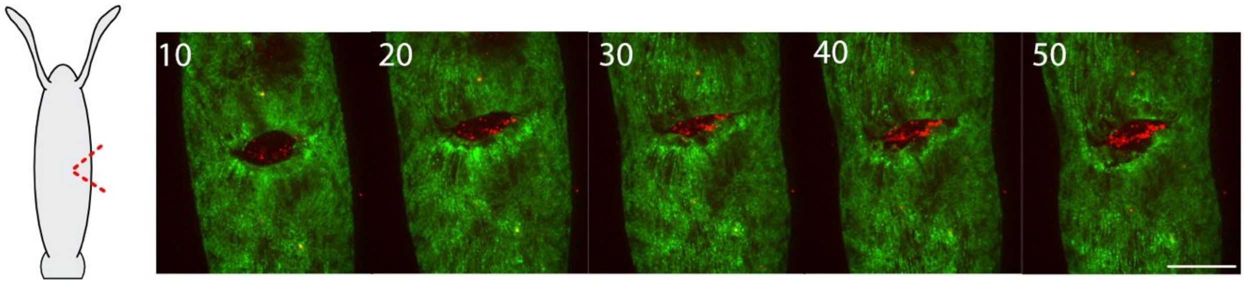

**Figure S8: MitoSOX dots detected in a LifeAct-eGFP animal injured in a non-regenerative context**

MitoSOX dots (red) detected in a LifeAct-eGFP animal injured in a non-regenerative context (lateral nick). Scale bar: 200 µm. Supplementary data to Fig. 3F.

**A**

| | Ctrl | Diphenyleneiodonium (DPI) | | |
|---|---|---|---|---|
| **Time:** | *0.2% DMSO* | *1µM* | *5µM* | *10µM* |
| 0.3 hpa | stage 1 | stage 1 | stage 1 | stage 1 |
| 1 hpa | stage 1 | stage 1 | stage 2 | stage 2 |
| 3 hpa | stage 1 | stage 2 | stage 2 | stage 2 |
| 24 hpa | stage 1 | stage 3 | stage 3 | stage 3 |

**B**

| | Ctrl | Tiron | | |
|---|---|---|---|---|
| **Time:** | *0.2% DMSO* | *5mM* | *15mM* | *20mM* |
| 0.3 hpa | stage 1 | stage 1 | stage 2 | stage 3 |
| 1 hpa | stage 1 | stage 1 | stage 1 | stage 3 |
| 3 hpa | stage 1 | stage 1 | stage 1 | stage 3 |
| 24 hpa | stage 1 | stage 1 | stage 2 | stage 4 |

**C**

| | Ctrl | 3-amino-triazole (ATZ) | | |
|---|---|---|---|---|
| **Time:** | *HM* | *10mM* | *50mM* | *100mM* |
| 0.5 hpa | stage 1 | stage 1 | stage 1 | stage 2 |
| 3 hpa | stage 1 | stage 1 | stage 1 | stage 2 |
| 24 hpa | stage 1 | stage 1 | stage 1 | stage 2 |
| 96 hpa | stage 1 | stage 1 | stage 1 | stage 4 |

**D**

| | |
|---|---|
| stage 1 | healthy with extended tentacles |
| stage 2 | healthy with shorter tentacles - reversible |
| stage 3 | contracted body column - reversible |
| stage 4 | dissociation (dying animals) - irreversible |

**Figure S9: Toxicity tests (A) DPI, (B) Tiron and (C) ATZ**

Ten Hv_AEP2 animals per condition were maintained in 5 ml in the continuous presence of drugs at indicated concentration. The toxicity stages were defined as reported in (ref[8]): **stage 1** - healthy; **stage 2** - healthy with shorter tentacles, which can reverse to the stage 1; **stage 3** - contracted body column and ball-shape tentacles, which can as well reverse to stage 1; **stage 4** - irreversible dissociation (dying animals).

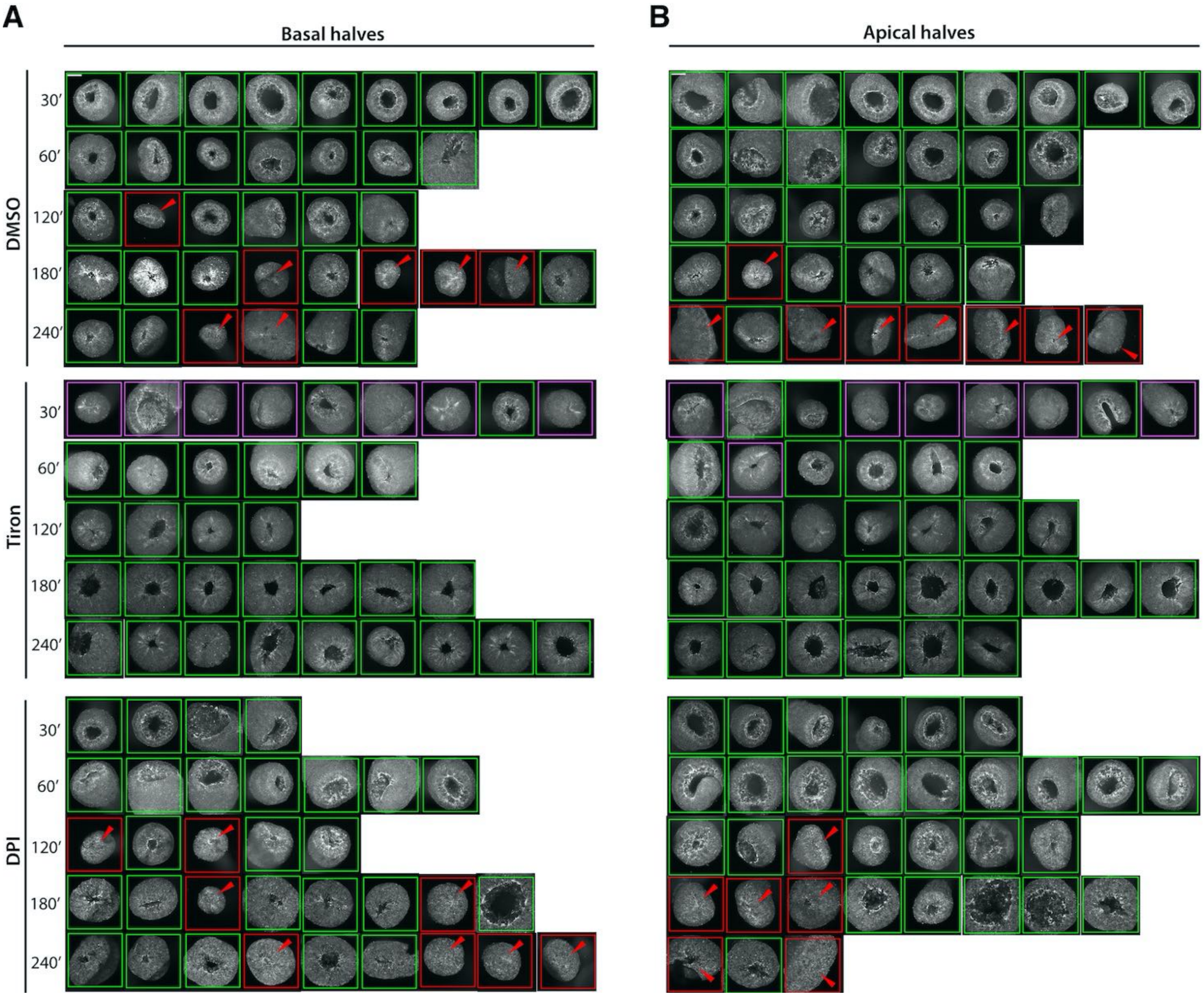

#### Figure S10: Wound closure of AR and BR halves exposed to Tiron or DPI

Wound closure of apical-regenerating **(A)** and basal-regenerating **(B)** halves from *Hv_AEP2* animals either untreated (DMSO) or treated with Tiron (15 mM) or DPI (10 µM) for one hour before mid-gastric bisection and continued up to 4 hours. Phalloidin stained animals were–pictured at indicated time points with the spinning disc microscope. Green frames indicate open wounds, red ones indicate closed wound, magenta ones indicate transiently contracted wounds. Scale bars: 200 µm. Supplementary data to Figure 4E, 4F

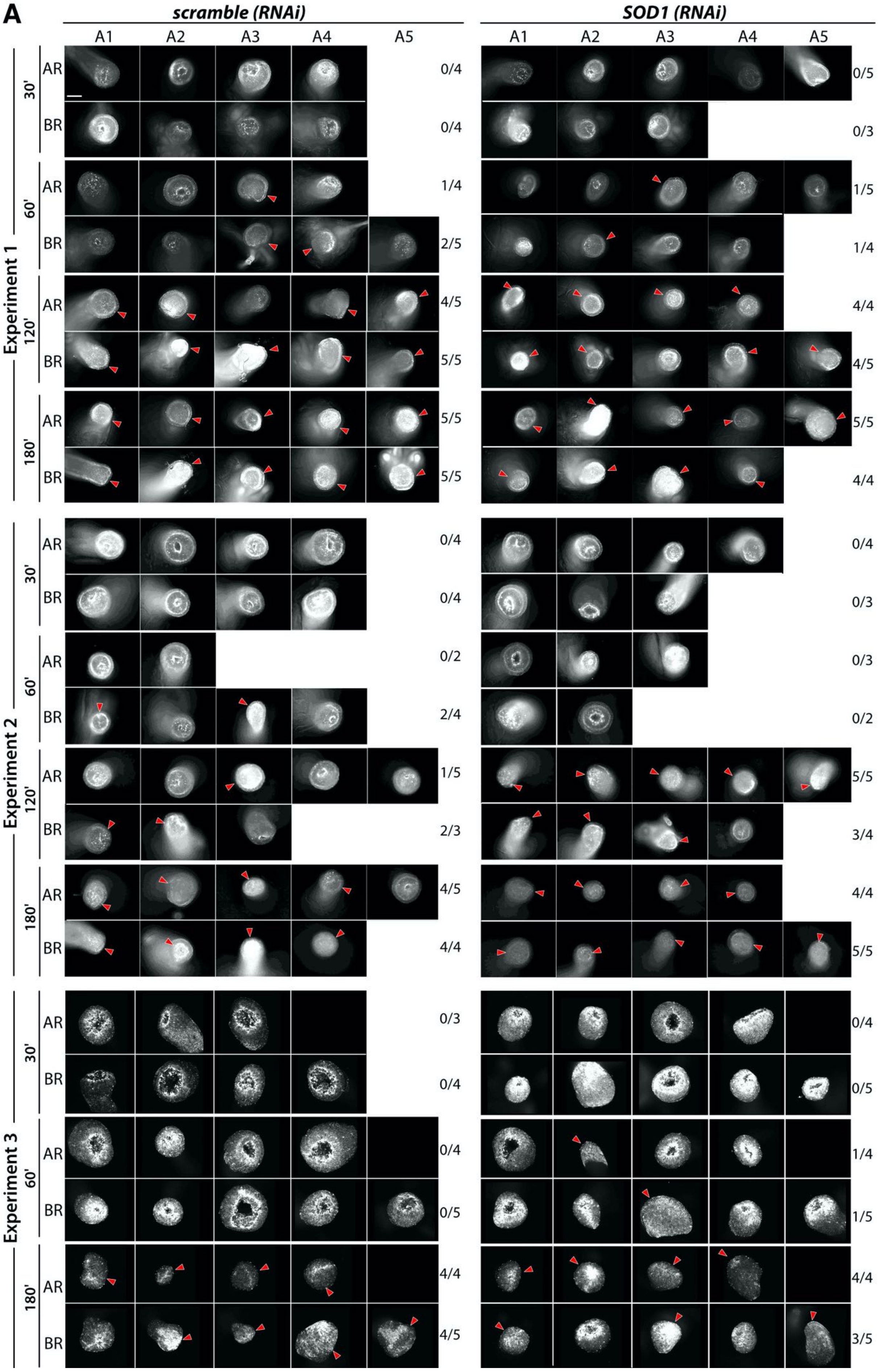

A
scramble (RNAi)
SOD1 (RNAi)
A1 A2 A3 A4 A5
A1 A2 A3 A4 A5
Experiment 1
30' AR 0/4 0/5
BR 0/4 0/3
60' AR 1/4 1/5
BR 2/5 1/4
120' AR 4/5 4/4
BR 5/5 4/5
180' AR 5/5 5/5
BR 5/5 4/4
Experiment 2
30' AR 0/4 0/4
BR 0/4 0/3
60' AR 0/2 0/3
BR 2/4 0/2
120' AR 1/5 5/5
BR 2/3 3/4
180' AR 4/5 4/4
BR 4/4 5/5
Experiment 3
30' AR 0/3 0/4
BR 0/4 0/5
60' AR 0/4 1/4
BR 0/5 1/5
180' AR 4/4 4/4
BR 4/5 3/5

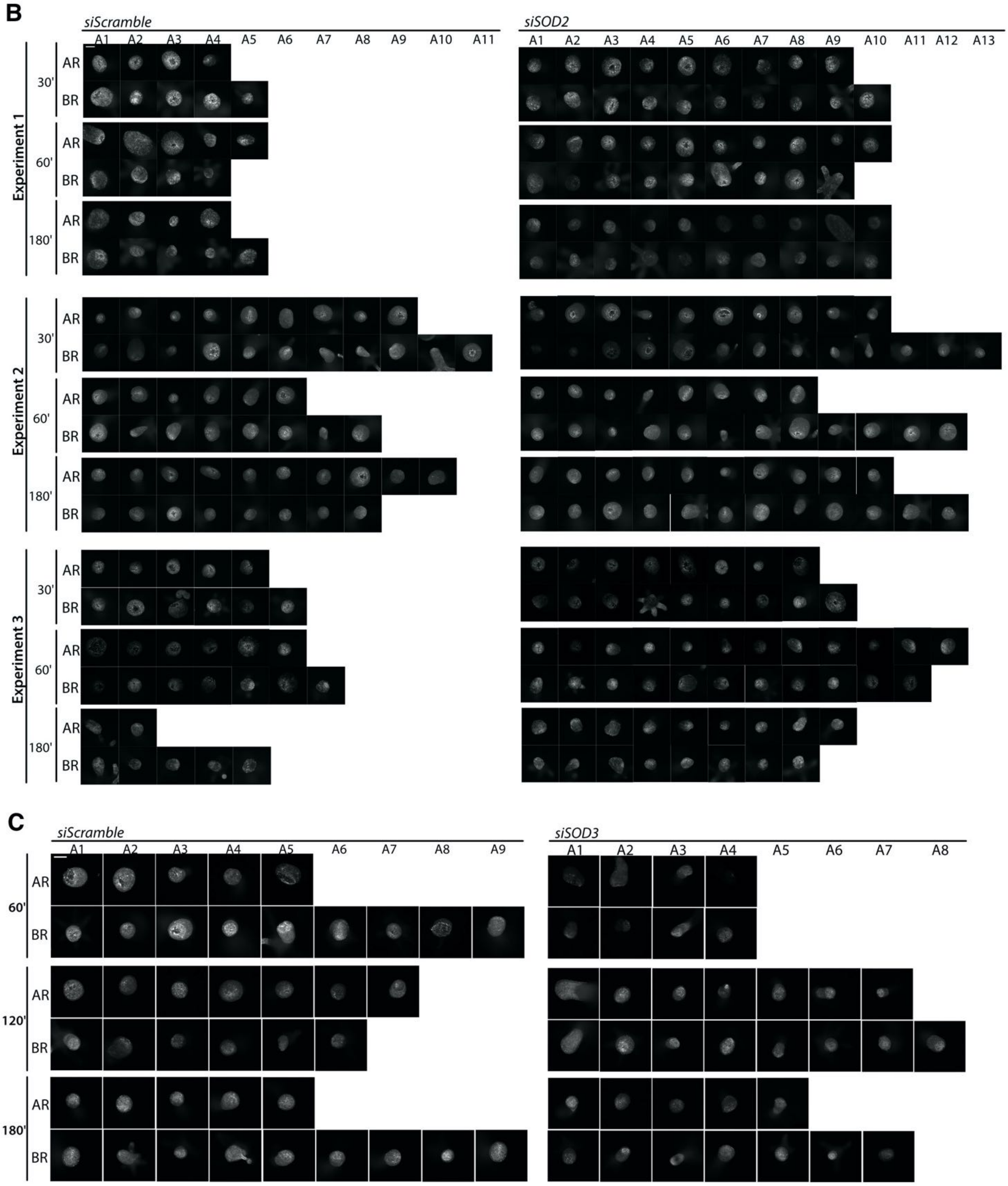

**Figure S11: Wound closure in animals knocked-down for *SOD1*, *SOD2 and SOD3***

Wound closure of apical-regenerating halves and basal-regenerating halves from *Hv_AEP2* animals exposed to siRNAs targeted against *SOD1* **(A)**, *SOD2* **(B)** and *SOD3* **(C)**. *SOD1*(RNAi) and *SOD2*(RNAi) experiments were repeated three times (experiment 1-3). Scale bar: 500 µm. Animals collected at indicated time point after mid-gastric bisection were stained with Phalloidin and pictured either with a Leica DM5500 wide-field fluorescent microscope (*SOD1*(RNAi) experiments 1 and 2) or with a Marianas 3i spinning disk confocal microscope (all other experiments). Images acquired on the wide-field fluorescent microscope tend to appear oversaturated post processing, therefore we applied dynamic fluorescence range. Images treated that way are the following: From experiment 1 *scramble* group: 30' A2-AR, 120' A1-BR ; 180' : A1-AR, A3-AR; and from *SOD1* RNAi group: 120' : A1-AR, A2-BR, A5-BR ; 180' : A3-AR, A1-AR ; from experiment 2 *scramble* group: 60' A1-BR; and from *SOD1* RNAi group: 30' A4-AR, 120' A3-BR. Supplement to Figure 5E-5J

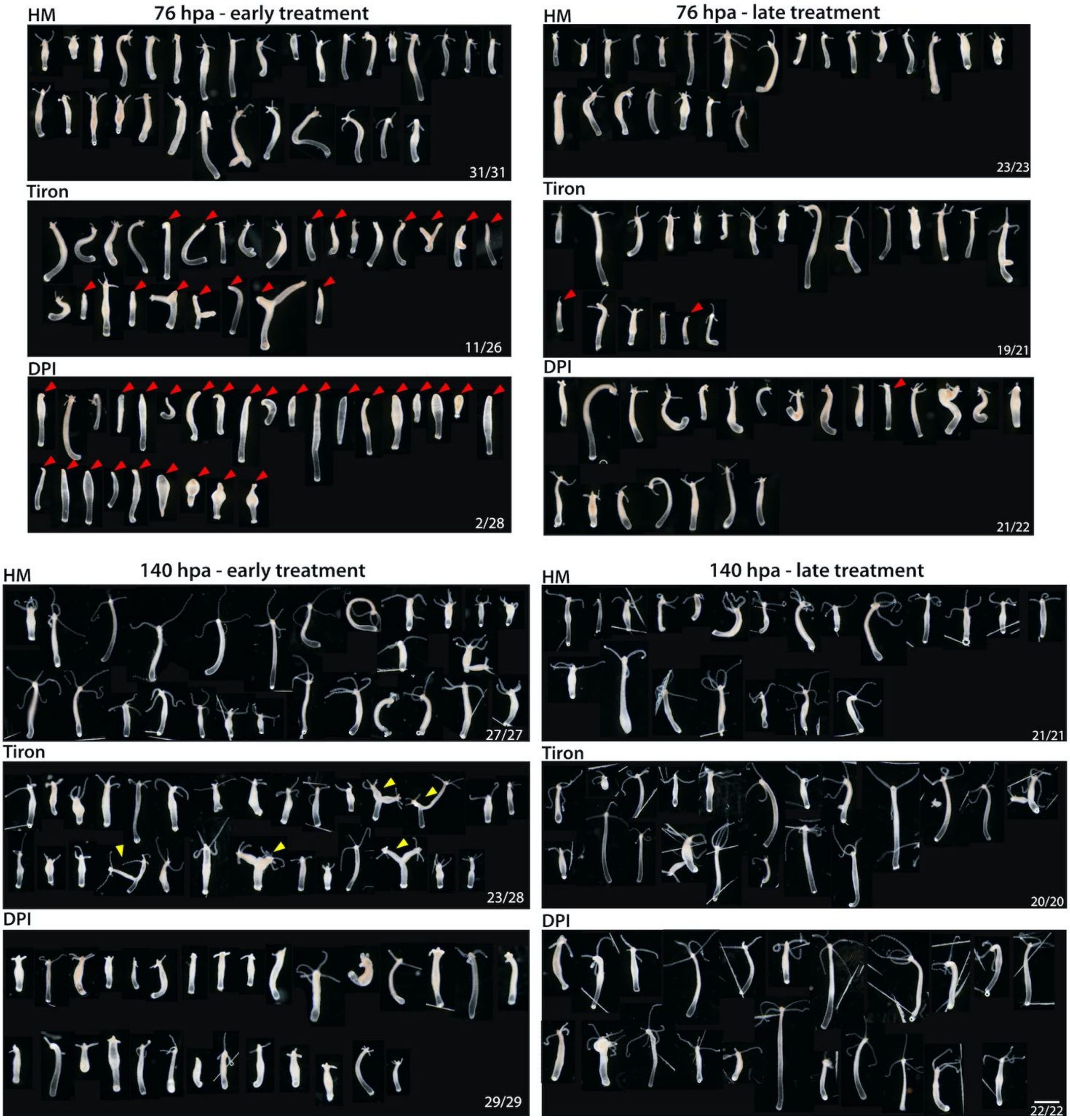

**Figure S12: Apical regeneration of animals exposed to Tiron or DPI and pictured 76 and 140 hours after mid-gastric bisection.**

Apical regeneration of animals exposed to Tiron (15 mM) or DPI (10 µM) for 25 hours either early (from -1 up to 24 hpa), or late (from 24 up to 48 hpa) pictured at 76 hpa and 140 hpa as indicated in Fig. 5A. Each animal was imaged separately. Red arrowheads point to non-regenerated extremities, yellow arrows point to abnormal regenerated heads. For each condition, the number of animals with regenerated heads is indicated. Scale bar: 1 mm. Supplementary data to Figure 6A, 6B

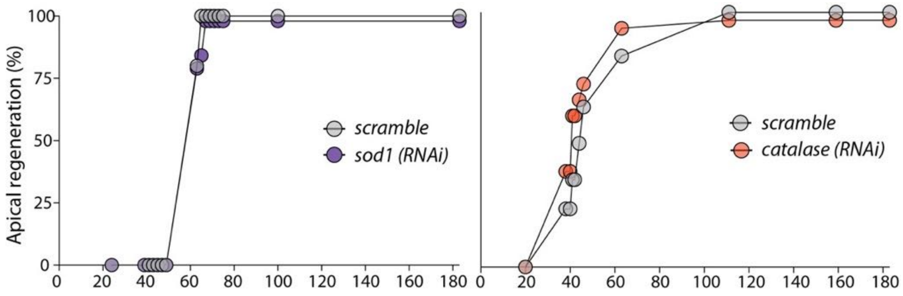

**Figure S13: Kinetics of apical regeneration in animals knocked-down for *SOD1* or *catalase***
Animals knocked-down for *SOD1* or *catalase* do not exhibit any delay in apical regeneration.

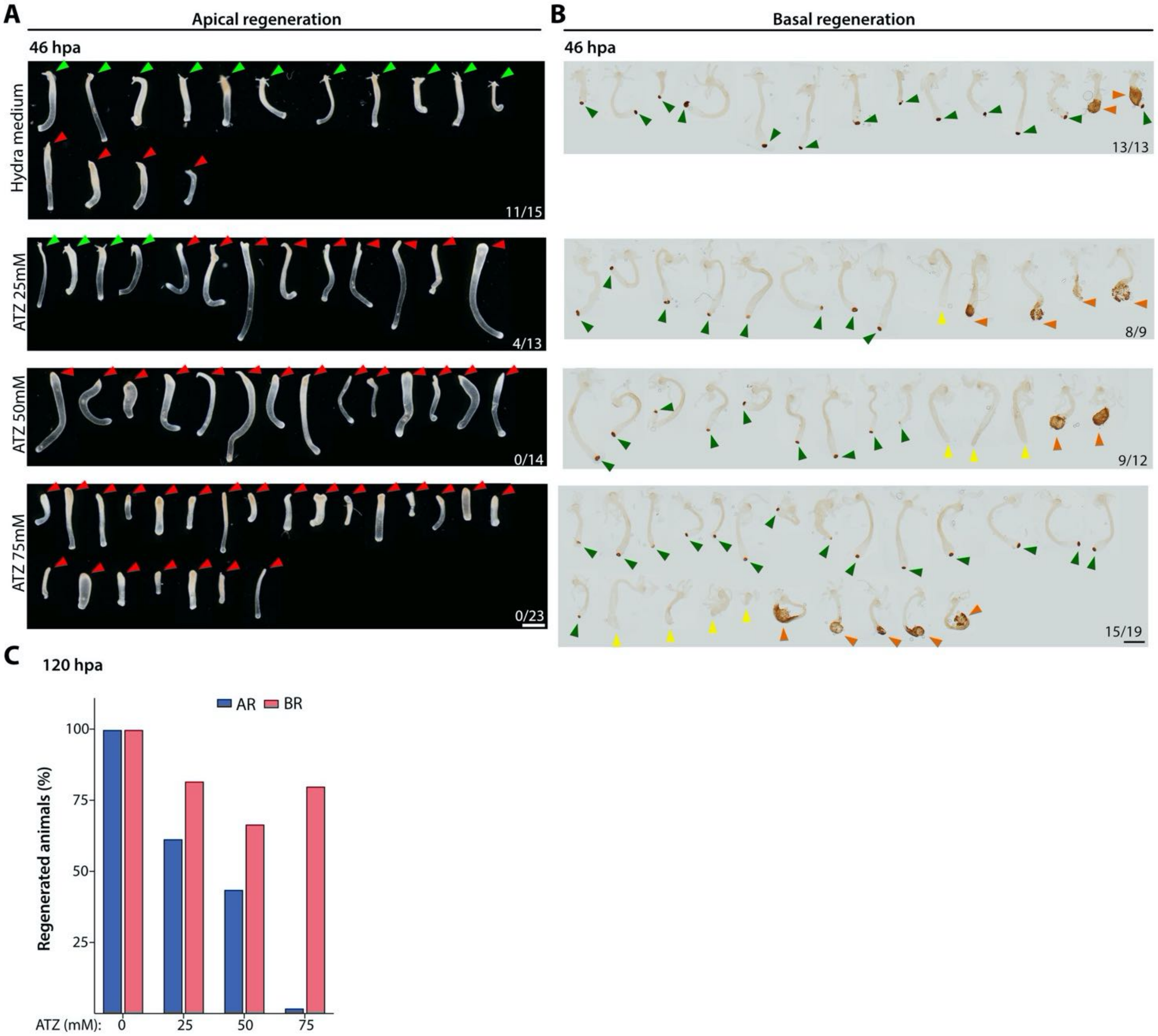

**Figure S14: Apical and basal regeneration in ATZ-treated animals**
**(A)** Apical morphology of all ATZ-treated regenerating animals pictured at 46 hpa. Green arrows indicate regenerated animals and red arrows are pointing to a defect in apical regeneration. Each animal from each condition was imaged separately. Scale bar: 1mm. **(B)** Basal morphology of all ATZ-treated regenerating animals pictured at 46 hpa. BR halves were stained for peroxidase. Green arrows indicate regenerated animals, yellow arrows are pointing to an absence of basal regeneration and orange arrows are showing the animals with an enlarged peroxidase+ basal region. Note that for each condition animals that are marked with orange arrows were excluded for the final quantification. Each animal from each condition was imaged separately. Scale bar: 1mm and 76 hpa. **(C)** Quantification for both apical and basal regeneration at 120 hours post amputation. Supplementary data to Figure 6D

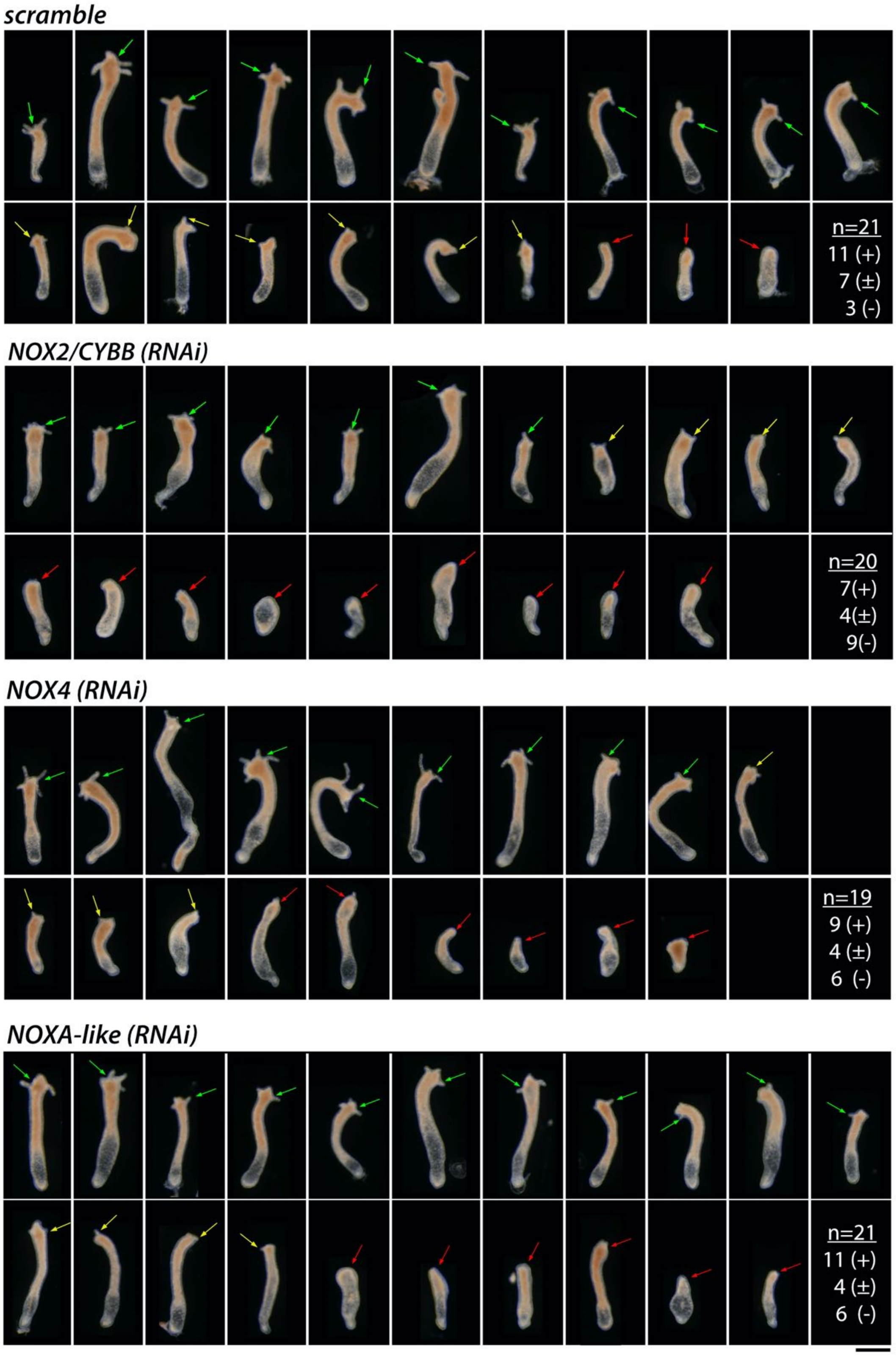

**Figure S15: Apical regeneration of *Hv_AEP2* animals knocked-down for *NOX2/CYBB, NOX4* and *NOXA-like***
Apical morphology of all siRNA-treated regenerating animals pictured at 48 hpa. Animals were electroporated 4x with siRNAs, once every other day, then bisected two days after EP4 and imaged 48 hours later. Arrows indicate apical extremities with visible growing tentacles (green (+)), emerging tentacle rudiments (yellow (±)) or without any sign of regeneration (red, (-)).Green arrows indicate regenerated animals and red arrows are pointing to a defect in apical regeneration. Scale bar: 500 µm. Supplementary data to Figure 6E

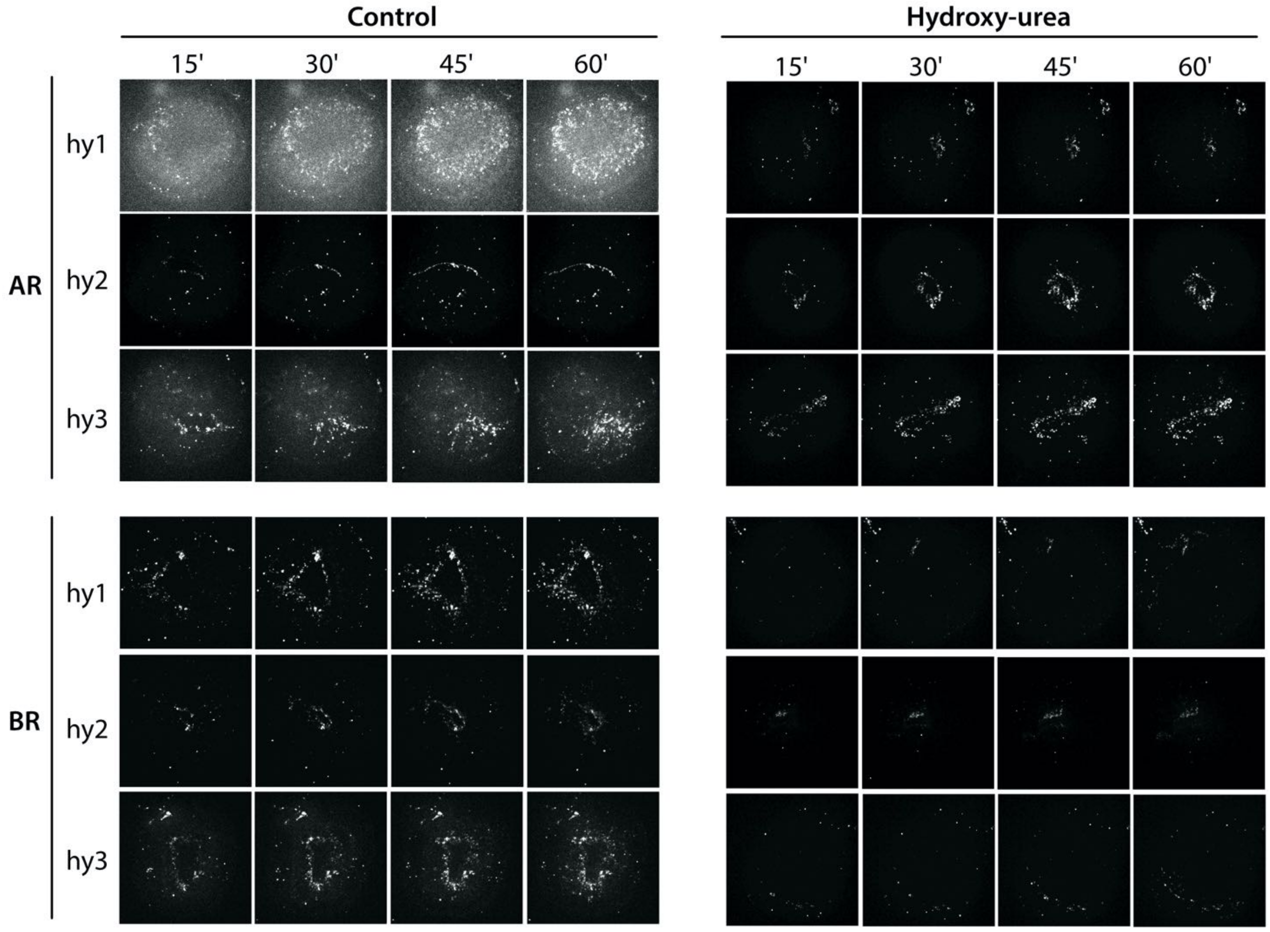

**Figure S16: MitoSOX detection of mtO2 in wounds of AR and BR halves from untreated and HU-treated animals.**

MitoSOX dots were counted as described in the Materials & Methods section. Although the images from HU-treated and control animals differ in intensity, the counting of MitoSOX dots provides similar values. Supplementary data to Figure 7F

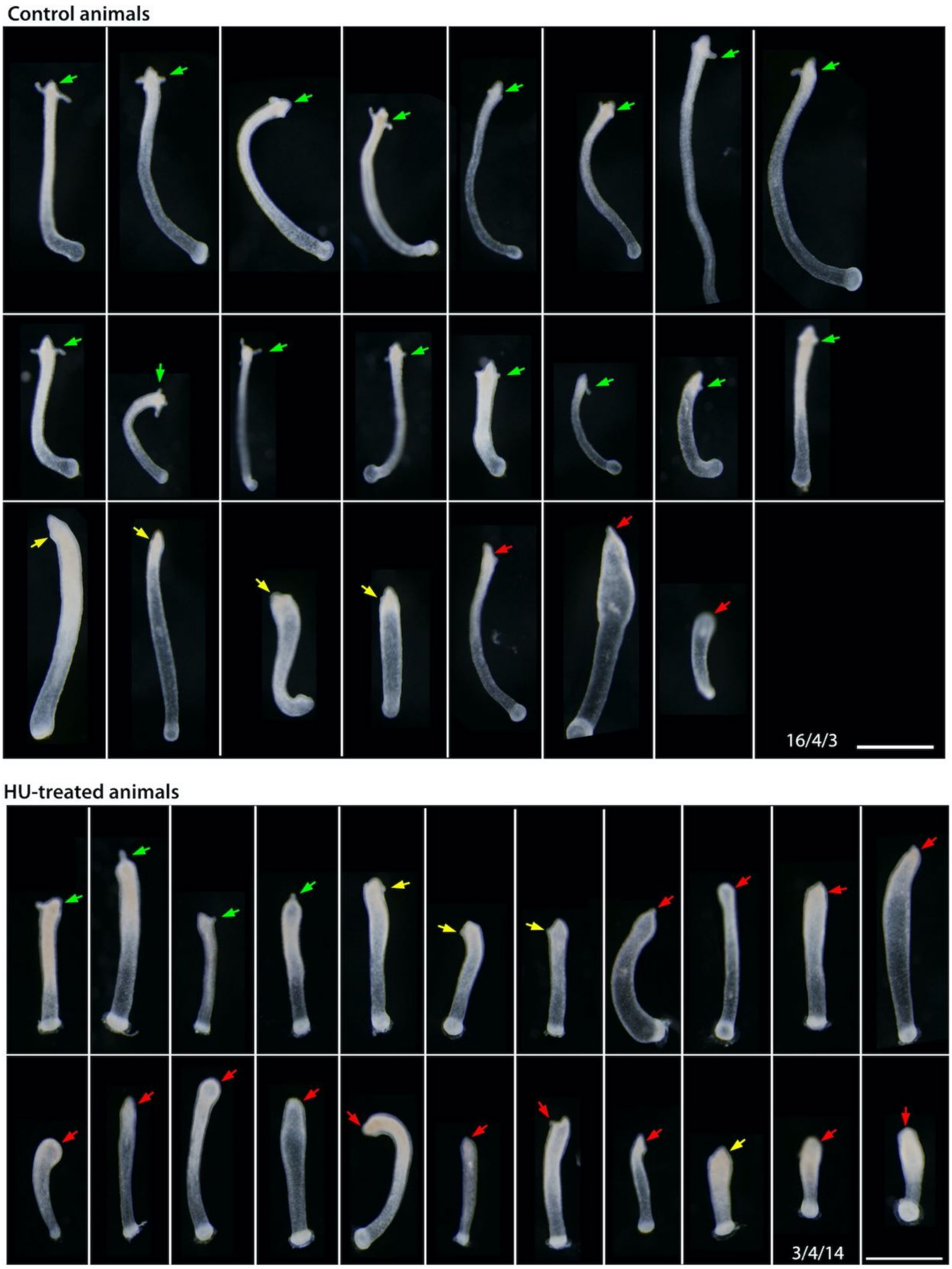

**Figure S17: Delay in apical regeneration in *Hv_AEP2* animals bisected 5 days after HU treatment**
Control and HU-treated regenerating animals were pictured at 46 hpa. Arrows indicate apical extremities with visible growing tentacles (green), emerging tentacle rudiments (yellow), abnormal or no sign of regeneration (red). The three numbers correspond to these three categories with n= 23 control animals and n= 21 HU-treated animals. Scale bar: 1 mm. Supplementary data to Figure 7H

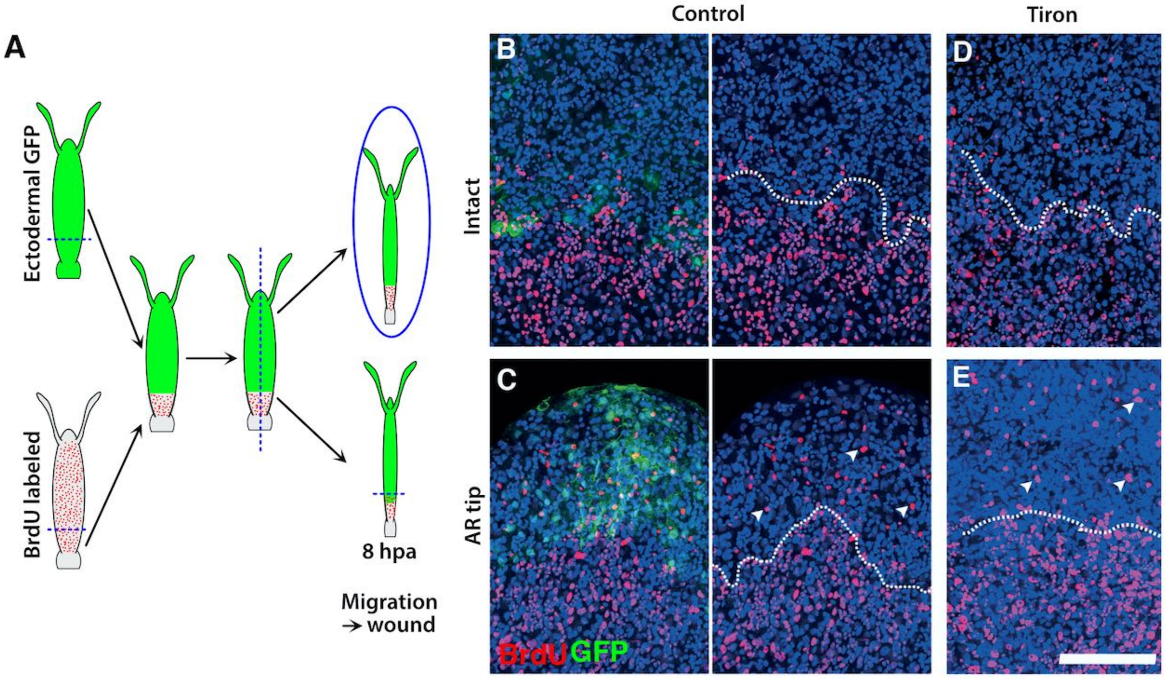

**Figure S18: Cell migration towards the wound is not affected upon Tiron treatment**

**(A)** Scheme depicting the axial grafting procedure performed between *Hv_AEP* animals that constitutively express actin::GFP in the epidermal cell layer (AEP-ECTO ref[9]) (green) and non-transgenic *Hv_AEP* animals pulse-labeled with BrdU for 14 hours before grafting (grey with red dots). **(B-E)** To observe the migration of BrdU-labeled cells from the basal region towards the central region of the body column, AEP-ECTO and non-transgenic AEP BrdU animals were bisected at 40% body length and the upper part of an AEP-ECTO animals was grafted onto the lower part of a wt *Hv_AEP* animal previously exposed to BrDU. Such chimeric *Hydra* were used to monitor cell migration in homeostatic and regenerative conditions: chimeric animals were bisected longitudinally 8 hours after grafting, providing two identical twin thin animals after closing. One of them was left intact (B, D), when the other was bisected at mid-gastric level (C, E). The dashed white line represents the grafting boundary. (B, D) Note the low number of BrdU-labeled cells (pink) that migrate along the body column of intact animals, either untreated (B) or exposed to 20 mM Tiron for 9 hours (D). (C, E) Numerous BrdU-labeled cells (white arrows) migrate in animals either left untreated (C) or exposed to 20 mM Tiron for 9 hours from one hour before bisection and fixed at 8 hpa (E). Scale bar 100 µm.

### REFERENCES

1. Chera, S. *et al.* Apoptotic Cells Provide an Unexpected Source of Wnt3 Signaling to Drive Hydra Head Regeneration. *Dev. Cell* **17**, 279–289 (2009).
2. Suresh Babu, C. V., Babar, S. M. E., Song, E. J., Oh, E. & Yoo, Y. S. Kinetic analysis of the MAPK and PI3K/Akt signaling pathways. *Mol. Cells* **25**, 397–406 (2008).
3. Wawrzak, D. *et al.* Wnt3a binds to several sFRPs in the nanomolar range. *Biochem. Biophys. Res. Commun.* **357**, 1119–1123 (2007).
4. Guindon, S. & Gascuel, O. A simple, fast, and accurate algorithm to estimate large phylogenies by maximum likelihood. *Syst Biol* **52**, 696–704 (2003).
5. Anisimova, M. & Gascuel, O. Approximate Likelihood-Ratio Test for Branches: A Fast, Accurate, and Powerful Alternative. *Syst. Biol.* **55**, 539–552 (2006).
6. Wenger, Y., Buzgariu, W. & Galliot, B. Loss of neurogenesis in Hydra leads to compensatory regulation of neurogenic and neurotransmission genes in epithelial cells. *Philos Trans R Soc Lond B Biol Sci* **371**, 20150040 (2016).
7. Wenger, Y., Buzgariu, W., Perruchoud, C., Loichot, G. & Galliot, B. Generic and context-dependent gene modulations during <em>Hydra</em> whole body regeneration. *bioRxiv* 587147 (2019) doi:10.1101/587147.
8. Bossert, P. & Galliot, B. How to use Hydra as a model system to teach biology in the classroom. *Int. J. Dev. Biol.* **56**, 637–52 (2012).
9. Wittlieb, J., Khalturin, K., Lohmann, J. U., Anton-Erxleben, F. & Bosch, T. C. Transgenic Hydra allow in vivo tracking of individual stem cells during morphogenesis. *Proc. Natl. Acad. Sci. U. S. A.* **103**, 6208–11 (2006).
